# Rational Design and Optimization of a Potent IDO1 Proteolysis Targeting Chimera (PROTAC)

**DOI:** 10.1101/2025.01.07.631731

**Authors:** Paige J. Monsen, Prashant V. Bommi, Arabela A. Grigorescu, Kristen L. Lauing, Yingyu Mao, Payton Berardi, Lijie Zhai, Oluwatomilayo Ojo, Manon Penco-Campillo, Taylor Koch, Michael Egozi, Sonam V. Jha, Sara F. Dunne, Hong Jiang, Guiqin Song, Fang Zhang, Steven Kregel, Ali Vaziri-Gohar, Sean Fanning, Pilar Sanchez-Gomez, Jacob M. Allen, Bakhtiar Yamini, Rimas V. Lukas, Derek A. Wainwright, Gary E. Schiltz

## Abstract

Indoleamine 2,3-dioxygenase 1 (IDO1) is a potently immunosuppressive protein that inhibits antitumor immunity through both tryptophan metabolism and non-enzymatic functions. Pharmacological therapies targeting IDO1 enzyme activity have generally failed to improve the overall survival of patients with cancer. Developing new therapeutic agents that are capable of neutralizing both enzyme-and non-enzyme-derived immunosuppressive IDO1 effects is therefore of high interest. We previously described the development of a novel Proteolysis Targeting Chimera (PROTAC), NU223612, that degrades IDO1 in cultured human glioblastoma (GBM) cells, as well as in well-established brain tumors, *in vivo*. In this study, we rationally optimized the composition, rigidity, and linker orientation of the PROTAC structure to create NU227326, which degrades IDO1 with a DC_50_ of 5 nM in human GBM cells. Mechanistic studies showed that IDO1 degradation occurred through the ubiquitin-proteasome system and was sustained for at least 2 days, supporting NU227326 as a highly potent IDO1 PROTAC suitable for further studies in GBM and other human cancers.

## INTRODUCTION

Glioblastoma (GBM; IDHwt) is the most common primary central nervous system cancer in adults and accounts for >50% of all malignant gliomas.^1^ Standard of care treatment for patients with GBM involves surgical resection of the tumor followed by radiation and adjuvant temozolomide chemotherapy.^2^ Despite this aggressive treatment regimen, the survival rate for GBM patients remains low with a median overall survival (mOS) rate of less than 20 months and a 5-year survival rate of less than 10%.^3^ The use of cancer immunotherapy to reinvigorate the antitumor immune response is an active area of research and has been a breakthrough for many types of cancers.^4^ Over the last 15 years, the development of immune checkpoint inhibitors (ICIs) targeting PD-1, PD-L1, and CTLA-4 have provided notable survival benefits to patients with a variety of different cancer types.^5, 6, 7^ However, treatment with ICIs has failed to improve the mOS of GBM patients across broad patient populations studied in phase 3 clinical trials to-date.^8, 9, 10^

The kynurenine (Kyn) pathway of tryptophan (Trp) metabolism has been studied for its role in tumor-induced immunosuppression.^11, 12, 13^ Indoleamine 2,3-dioxygenase 1 (IDO1) is a rate-limiting enzyme that converts the essential amino acid, Trp, into downstream Kyn metabolites. Low Trp or high Kyn levels suppress T cell function(s) and/or induce cell death when evaluated *in vitro*.^14, 15^ Expression of IDO1 is prevalent in a number of cancers including GBM^16, 17^ and high expression is associated with poor survival outcomes. ^17, 18, 19, 20, 21, 22^ ^23–25^ Several IDO1 enzyme inhibitors including epacadostat (**1**),^26, 27^ navoximod (**2**),^28, 29^ PF-0684003 (**3**),^30^ and BMS-986205 (**4**)^31, 32^ have been widely studied over the past decade (Figure 1). To-date, small molecule IDO1 enzyme inhibitor treatment has generally failed to provide a survival benefit to patients with cancer. This has resulted in questioning whether IDO1 is a valid target for cancer immunotherapy.^27, 33^ Beyond IDO1 enzyme-dependent effects, we and others have demonstrated that IDO1 also possesses non-enzymatic functions that mediate immunosuppression.^23–25, 34^ The data collectively support the hypothesis that one of the reasons IDO1 enzyme inhibitors have failed to improve cancer patient survival among clinical trials to-date is due to IDO1 non-enzyme-mediated mechanisms. Therefore, the simultaneous targeting of both IDO1 enzyme and non-enzyme activities are required to effectively inhibit immunosuppression and achieve immunotherapeutic efficacy for glioblastoma.

**Figure 1.**
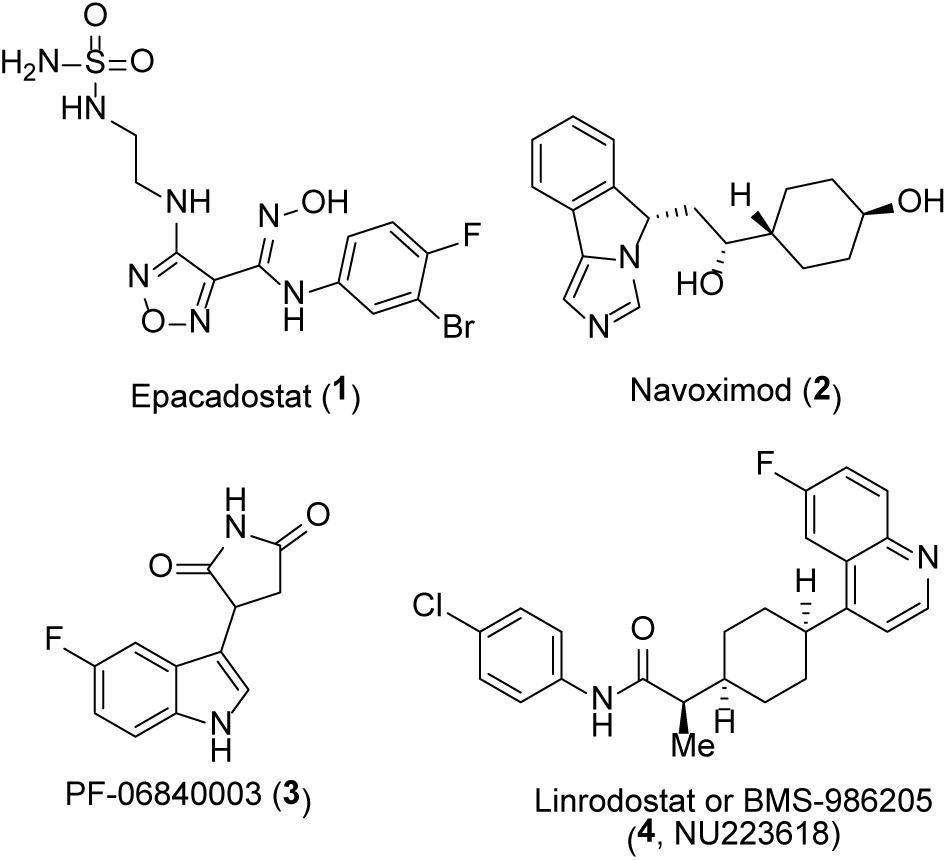
Representative IDO1 inhibitors.

Targeted protein degradation (TPD) is a powerful approach to therapeutically regulate proteins that are termed “undruggable”, embody undesirable non-enzymatic functions, or have the potential to develop mechanisms of resistance when treated with small molecule targeted enzyme inhibitors.^35, 36^ Proteolysis targeting chimeras (PROTACs) are heterobifunctional small molecules that exploit the ubiquitin-proteasome system for targeted protein degradation.^37^ Structurally, a PROTAC consists of three components: a targeting ligand that is specific for recruiting and binding to the protein of interest (POI), a warhead ligand to recruit and bind to an E3 ligase, and a linker to tether the two binding ligands (Figure 2a). Over the past 20 years, the development and optimization of PROTACs to become drug candidates has validated this approach in medicinal chemistry and drug discovery.^35^ In addition to PROTACs providing a unique drug discovery approach, they can be utilized as a chemical knockdown tool to further study the biology and non-catalytic roles of proteins. Therefore, the investigation into developing PROTACs to degrade IDO1 protein and inhibit IDO1 enzyme and non-enzyme activities represents a promising strategy to fully inhibit IDO1-mediated immunosuppression as well as further study the role of IDO1 in cancer.

**Figure 2.**
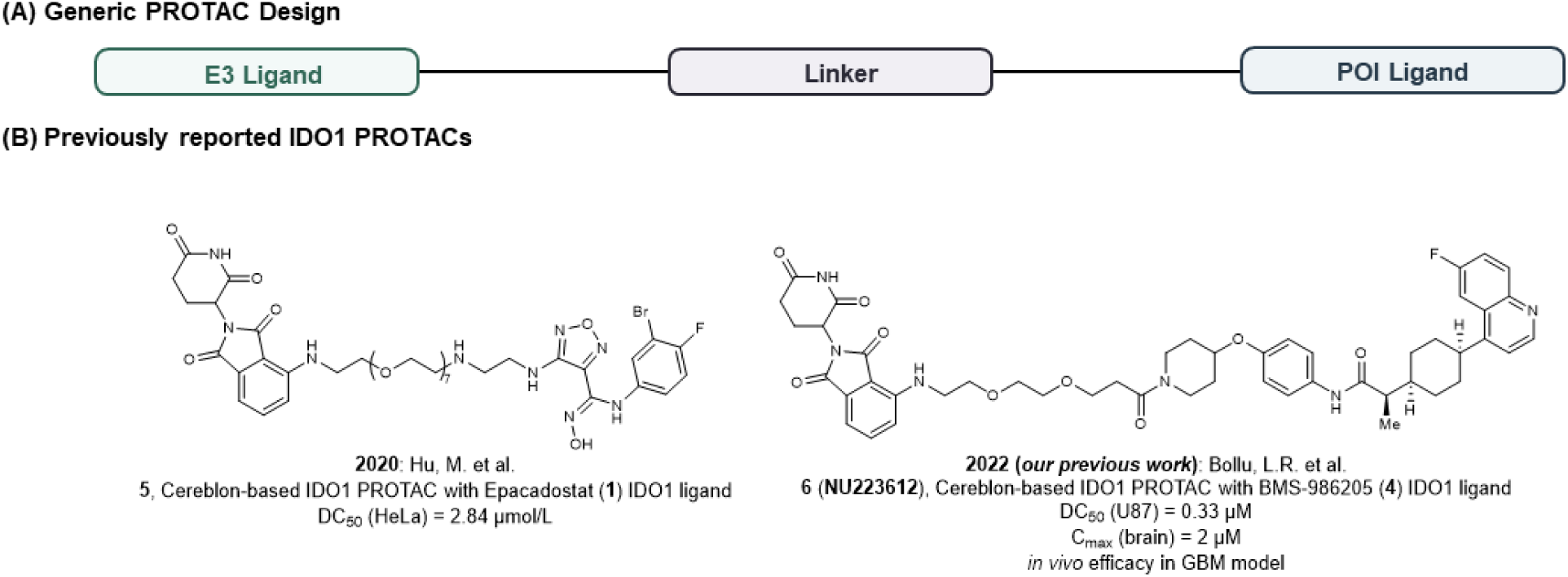
(A) General PROTAC design. (B) Structures of reported IDO1 PROTACs to date.

To date, the characterization of two potent IDO1 PROTACs have been reported (Figure 2b). In 2020, Hu *et al*. developed the first IDO1 degrader, **5**, consisting of epacadostat (**1**) as the IDO1 targeting ligand and the cereblon (CRBN) ligand, pomalidomide, as the E3 ligase recruiting ligand.^38^ The epacadostat-based degrader displayed a DC_50_ = 2.8 µM in cultured HeLa cells however no *in vivo* validation studies were performed. In 2022, our group reported the development of an IDO1 PROTAC, **6** (**NU223612**), incorporating the BMS-986205 (**4**) enzyme inhibitor as the IDO1 targeting ligand into a CRBN-based PROTAC for subjects with an intracranial primary brain tumor.^39^ Our work focused on using BMS-986205 as the IDO1-targeting ligand because this compound had fewer hydrogen-bond donors than epacadostat, making it more likely to have better cell permeability and brain penetration. BMS-986205 also possesses high selectivity for IDO1 over related enzymes IDO2 and TDO, it advanced into Phase 3 clinical trials suggesting it possesses a favorable off-target and toxicity profile, and importantly, X-ray crystal structures of it bound to IDO1 suggested clear opportunities to attach a solvent-exposed linker.^40^ Degrader **6** possessed a DC_50_ = 0.33 µM in cultured U87 human GBM cells, reached a concentration of 2 µM in brain tissue (IP), and was able to degrade IDO1 in established brain tumors that yielded a survival benefit in mice with intracranial GBM.^39^

Our previous work described the first report of a PROTAC possessing sufficient brain exposure to produce *in vivo* efficacy in a GBM model and established the therapeutic potential of developing IDO1 PROTACs for the treatment of GBM.^39^ Although encouraged by these results, we recognized an opportunity to further optimize our lead series of degraders to (i) enhance the IDO1 degradation potency and efficacy and (ii) improve the brain exposure and pharmacokinetic (PK) characteristics of the IDO1 degrader. Recently, there has been increasing reports of structural factors and physicochemical properties of PROTACs, such as molecular weight (MW), rigidity or number of rotatable bonds (RB), and the number of hydrogen bond donors (HBDs) and acceptors (HBAs), playing influential roles in potency, permeability and ADME (absorption, distribution, metabolism and excretion) properties.^41–43^ Herein, we report the optimization of lead IDO1 PROTAC, **6** (**NU223612**), in which structural modifications to the CRBN ligand, linker, and IDO1 ligand were performed in a modular fashion to study the influence on potency, efficacy, and bioavailability.

## RESULTS

### Design of potent 2^nd^ Generation IDO1 PROTACs

Previously, we reported the development of a 1^st^ generation series of degraders to establish IDO1 PROTAC proof of concept and obtain structure-activity relationship (SAR) data.^39^ Molecular docking studies revealed the phenyl ring of the parental IDO1 inhibitor, **4** (**NU223618**, BMS-986205), is solvent exposed and provided a suitable anchor for the attachment of linkers. Therefore, IDO1 degraders were designed with an ether connection of the linker to the phenyl ring at either the 3-position to retain the chlorine atom native to the inhibitor or at the 4-position, thereby replacing the chlorine atom. Docking studies were performed with truncated analogs of both substitution options to allow for further investigation into the influence the linker substitution to the warhead has on binding to the IDO1 protein (Figure 3A). When overlaid with the bound IDO1 inhibitor (BMS-116), the analog with the ether-piperidine group at the 4-position results in movement of the targeting ligand to accommodate the exiting of the linker into the solvent (Figure 3B and Figure 3D). This movement leads to a loss in interaction between the amide NH of the ligand and Serine 167 (Figure 3D). Conversely, attachment of the ether-piperidine group at the 3-position allows for a direct overlap with the parental inhibitor and restores the key interaction between the amide *N-H* and Serine 167, presumably providing greater binding affinity for IDO1 (Figure 3CD and Figure 3E). In addition to restoring this interaction, the orientation of the exit vector out of the binding site is slightly modified when the linker is attached to the 4-chloro-3-position versus the 4-position and therefore may allow for the PROTAC to facilitate a unique ternary complex upon binding to the IDO1 and E3 ligase proteins. These observations suggest the 4-chloro-3-substituted position may be a preferred connection site for linkers in comparison to the 4-substituted position and was therefore explored further during the initial optimizations of lead degrader, **6** (**NU223612**) (Figure 4).

**Figure 3.**
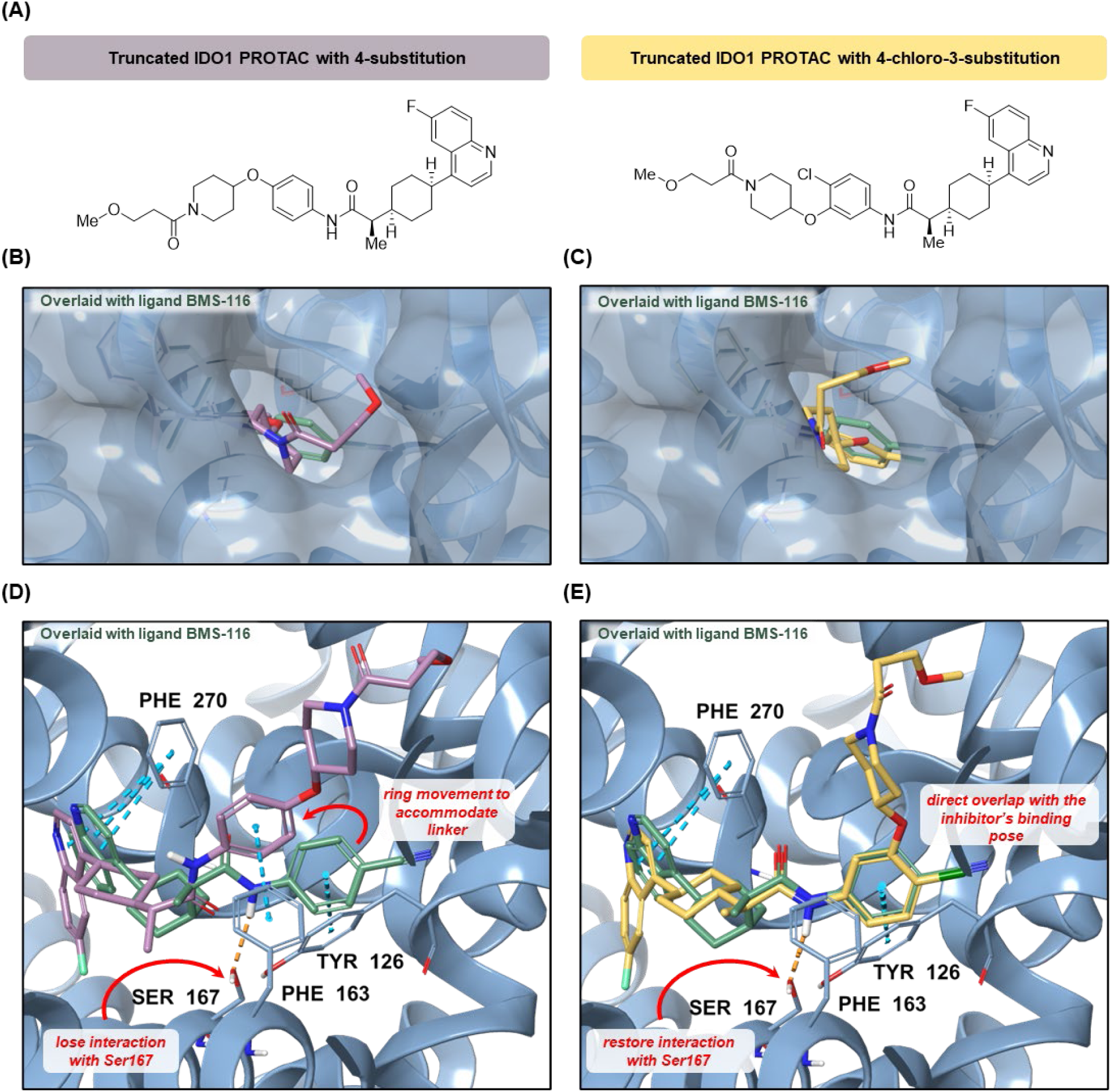
(A) Structures of truncated 4-substituted and 4-chloro-3-substituted type IDO1 PROTACs used in docking studies. (B and C) Surface representations of the co-crystal structure of IDO1 (blue PDB; 6AZW) with the ligand BMS-116 (green) overlaid with either the 4-substituted type analog (purple) or the 4-chloro-3-substituted type analog (gold) demonstrating the orientation of the exit vector and linker. (D and E) Binding pose of BMS-116 (green) modeled with either the 4-substituted type analog (purple) or the 4-chloro-3-substituted type analog (gold) in the IDO1 active site with representative interactions displayed to demonstrate the influence in altering the substitution position of the linker.

**Figure 4.**
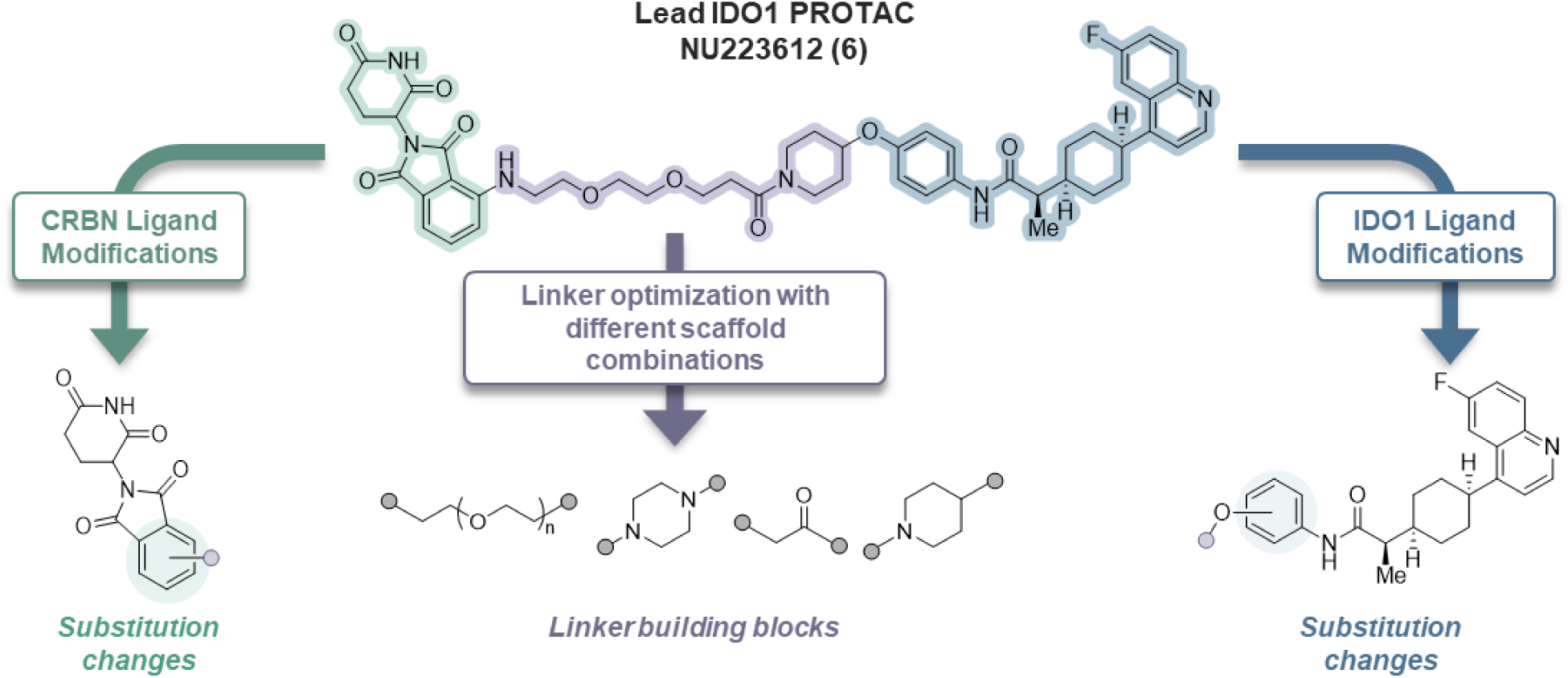
Design of novel IDO1 PROTACs based on modular structural optimizations of lead IDO1 PROTAC **6** (**NU223612**).

As previously reported, a variety of E3 ligase ligands were investigated in the initial screening and SAR analysis revealed both CRBN and Von Hippel-Lindau (VHL)-type ligands can be used as recruiting warheads in the PROTAC design to induce degradation of IDO1. To improve the PK profiles and blood brain barrier (BBB) penetration of the IDO1 degraders, CRBN-based PROTACs were of more interest as they have both decreased MW and fewer RBs compared to VHL-based PROTACs. Consequentially, optimization of degrader **6** (**NU223612**) involves simple modifications to the CRBN ligand, specifically through altering the substitution position to the linker scaffold (Figure 4). Numerous reports provide increasing evidence that linker length and composition play a major role in the bioactivity of a PROTAC.^35, 44^ Since SAR data from the original degrader library suggested amine-connected linkers to the CRBN ligand are preferred in comparison to ether or oxyacetamide type functional groups, we further investigated the amine-type connectivity to the E3 warhead. Utilizing various linker building blocks, the flexible polyethylene glycol (PEG) linker could be replaced entirely or in part with short, rigid heterocycles, which would allow us to evaluate the degradation of IDO1 with less-flexible linkers. Therefore, taking into consideration the SAR data established in our initial report, we developed a new series of IDO1 degraders.

### Optimization of IDO1 Degraders

Altering the substitution positions of the linker to both the CRBN and IDO1 ligands was undertaken to investigate the effect on IDO1 degradation. To quantitatively determine the degradation potency (DC_50_) and efficiency (D_max_) of the designed IDO1 degraders, a HiBiT degradation assay for IDO1 protein in the U87 glioblastoma cell line was developed and employed. HiBiT was inserted by CRISPR/Cas9 to the N-terminus of IDO1 protein and degradation was obtained through measuring the loss of luminescence following a 24 h treatment with the synthesized PROTACs.^45^

The initial IDO1 degraders contained 4-substituted thalidomide-type CRBN ligands, therefore altering the linker substitution to the 5-position was explored. Furthermore, investigation into the IDO1-ligand substitution preference was performed through analyzing both the 4-substituted and 4-chloro-3-substituted phenyl rings. Therefore, while keeping the amino-PEG2-amide-piperidine linker consistent, novel analogs **7** – **9** were synthesized to embody differential substitution patterns to both the CRBN and IDO1 ligands and were evaluated for IDO1 degradation (**Table 1**). Changing the substitution position to the CRBN ligand from the 4-to the 5-postion resulted in inactive degrader, **7**. Similarly, modifying the substitution of the IDO1 ligand from the 4-substitution to the 4-chloro-3-substitution position yielded inactive analog, **8**. Altering both the substitution to the CRBN and IDO1 ligands provided PROTAC, **9**, with decreased potency (DC_50_ >10 μM degradation).

**Table 1.** Initial optimization of IDO1 PROTACs.

| Compound | CRBN ligand | IDO1 ligand | IDO1 Degradation<sup>a</sup> |  | Linker |
| --- | --- | --- | --- | --- | --- |
|  |  |  | DC<sub>50</sub> (μM) | D<sub>max</sub> (%) |  |
| **6** | 4 | 4-substituted | 0.43 | 79 |  |
| **7** | 5 | 4-substituted | >10 | <10 |  |
| **8** | 4 | 4-chloro-3-substituted | >10 | <10 |  |
| **9** | 5 | 4-chloro-3-substituted | >10 | <10 |  |
| **10** | 4 | 4-substituted | >10 | <10 |  |
| **11** | 5 | 4-substituted | 0.34 | 26 |  |
| **12** | 4 | 4-chloro-3-substituted | >10 | <10 |  |
| **13** | 5 | 4-chloro-3-substituted | 0.008 | 48 |  |
| **14** | 4 | 4-substituted | >10 | <10 |  |
| **15** | 5 | 4-substituted | >10 | <10 |  |
| **16** | 4 | 4-chloro-3-substituted | >10 | <10 |  |
| **17** | 5 | 4-chloro-3-substituted | >10 | <10 |  |
| **18** | 5 | 4-substituted | 0.034 | 21 |  |
| **19** | 5 | 4-chloro-3-substituted | 0.007 | 51 |  |

<sup>a</sup>IDO1 degradation activity was tested using HiBiT assay with 24 h treatment time. DC<sub>50</sub> = dose that reduces IDO1 protein by 50%. D<sub>max</sub> = maximum reduction of IDO1 protein achieved by the compound.

The observed loss in potency resulting from small changes of the connectivity to the CRBN and IDO1 ligands lead us to focus our efforts on optimization of the linker moiety of the IDO1 degrader series. Therefore, we replaced the entire flexible amino-PEG2 with a rigidified linker scaffold to provide PROTACs **10** -**13**. To retain some flexibility, a methylene unit was utilized as a connecting scaffold between the piperazine and piperidine ring systems. Replacement of both the hydrogen bond donating amine and the entire PEG linker resulted in inactive analog **10**. Modifying the substitution to the CRBN ligand from the 4-to 5-position resulted in degrader **11** with slightly enhanced potency (DC_50_ = 340 nM) but decreased efficacy (D_max_ = 26%). Keeping the 4-subsituted CRBN ligand intact while altering the IDO1 ligand substitution from the 4-to the 4-chloro-3-subsituted to provide analog **12**, lead to a complete loss in potency. Interestingly, changing both the CRBN and IDO1 substitutions yielded a highly potent degrader **13**, with a DC_50_ of 7.6 nM and a D_max_ of 48%, suggesting that the rigidification of the linker may be more tolerated when connected at the CRBN 5-position and the IDO1 4-chloro-3-position. The enhanced IDO1 degradation resulting from replacement of the flexible linker in analogs **7** and **9** with a more rigid linker in degraders **11** and **13** suggests the linker composition is highly responsible for degradation activity.

To investigate the optimized linker rigidity, we replaced the central piperidine ring with a flexible PEG_1_ unit to yield analogs **14** – **17**. De-rigidifying the central unit of the PROTAC linker, while retaining rigid ring systems to both the E3 and IDO1 ligands led to a complete loss in IDO1 degradation potency, implying the entire flexible PEG linker should be replaced with a rigid linker.

To further increase the rigidity of the more potent analogs, **11** and **13**, the methylene unit was removed, providing degraders **18** and **19**, which both possessed the 5-subsitution to the CRBN ligand though different substitutions to the IDO1 ligand, 4-subsituted and 4-chloro-3-substituted, respectively. Shortening the length of the linker by only one carbon increased the potency by 10-fold for the 4-substituted degrader (**18**, DC_50_ = 34 nM), though the D_max_ remained at just 21%. The 4-chloro-3substituted analog, **19**, displayed both enhanced potency with a DC_50_ value of 6.6 nM and enhanced efficiency with a D_max_ of 51%.

Taken all together, the above data demonstrates that the combination of the linker composition and the substitution patterns to the E3 ligase and IDO1 ligands collectively play an important role in the ability of the PROTACs to reduce levels of IDO1 in U87 cells. Additionally, we were able to establish the optimal structural combination to include the 5-subsituted CRBN ligand and the 4-chloro-3-substituted IDO1 ligand with a highly rigid nitrogen-heterocycle containing linker.

### Optimization of Degraders 13 and 19

The initial linker and substitution optimizations led to the identification of degraders **13** and **19** which possessed improved potency for IDO1 degradation compared to the initial lead PROTAC **6**. Analysis of the structural modifications made to the degrader series suggests that replacement of the flexible PEG linker with a more rigid, nitrogen heterocycle-containing linker can be achieved with the optimal combination of substitution patterns to both the E3 ligase and IDO1 ligands. More specifically, a common trend was noted, in which 4-chloro-3-substitution to the IDO1 ligand was more preferential compared to the 4-substitution. Based on this observation, the previously described docking studies, and the increased potency correlating to the IDO1 substitution modification, we chose to focus our attentions on optimizing the exit vector of the IDO1 ligand. Since substitution at the 3-position provides an ideal orientation for the linker to exit (Figure 3E), we kept the substitution of the linker to the phenyl ring consistent and removed the chlorine atom (Table 2). We hypothesized that replacement of this larger substituent with a hydrogen atom would allow for more flexibility of the targeting ligand within the binding site to accommodate the attached ether-piperidine linker, while still allowing for the formation of key interactions. Therefore, we synthesized several 3-substituted IDO1 PROTACs with various linkers and analyzed them for IDO1 protein degradation using the HiBiT degradation assay (Table 2).

**Table 2.** Optimization of *meta*-substituted IDO1 PROTACs.

| Compound | Linker | IDO1 Degradation<sup>a</sup> |  |
| --- | --- | --- | --- |
|  |  | DC<sub>50</sub> (μM) | D<sub>max</sub> (%) |
| **20** |  | 0.020 | 67 |
| **21** |  | 0.005 | 63 |
| **22** |  | >10 | <10 |
| **23** |  | >10 | <10 |
| **24** |  | >10 | <10 |

<sup>a</sup>IDO1 degradation activity was tested using HiBiT assay with 24 h treatment time. DC<sub>50</sub> = dose that reduces IDO1 protein by 50%. D<sub>max</sub> = maximum reduction of IDO1 protein achieved by the compound.

We began by investigating **13** and **19** derived analogs, in which the CRBN substitution and linkers were kept consistent, and the chlorine atom on the IDO1 ligand was removed to provide the corresponding IDO1 PROTACs, **20** and **21**. Upon removal of the chlorine group, an increase in IDO1 degradation potency and efficiency was observed. Degrader **20**, the analog of PROTAC **13**, exhibited a DC_50_ of 20 nM and a D_max_ of 67% and **21**, the direct analog of **19**, possessed a DC_50_ of 4.5 nM and a D_max_ of 63%. In an effort to decrease the MW of the degraders, the linker was shorted via the removal of the central piperidine ring to yield analogs **22** and **23**. Reducing the length of the linker resulted in a complete loss of potency, suggesting that shortening the length of the linker disrupted the ability of the degraders to form the ternary complex and induce IDO1 degradation.

### *In vitro* analysis of potent IDO1 PROTACs 20 and 21

Analogs **20** and **21** were identified as novel lead IDO1 degraders possessing DC_50_ values of 20 nM and 4.5 nM, respectively, in the HiBit degradation assay (Figure 5A). Therefore PROTACs **20** and **21** underwent additional and extensive *in vitro* analysis. Both **20** and **21** were subjected to western blot analysis at an extended dose range to measure IDO1 degradation effect in cultured U87 cells (Figure 5B). The results indicated both analogs dose-dependently degrade IDO1 with **20** reaching maximum degradation at 300 nM (68% IDO1 degraded) and **21** reaching maximum degradation at 100 nM (88% IDO1 degraded). In addition to their IDO1 degradation potency, both **20** and **21** dose dependently inhibit the amount of Kyn levels present in treated U87 cells (Figure 5C). Since IDO1 PROTAC **21** displayed the highest percentage of IDO1 degradation at the lowest dose, **21** was further evaluated in human GBM43 cells. Degrader **21** was found to dose-dependently decrease protein levels in GBM43 cells as well as decrease the levels of Kyn, similarly to the response in U87 cells (Figure 5D and Figure 5E). Additionally, the DC_50_ was determined through evaluating the concentration of the drug at which 50% of the IDO1 protein is degraded. A DC_50_ of 7.1 nM and 11.8 nM in U87 and GBM43 cells, respectively, was determined (Figure 5F). It should be noted that a hook effect is observed for the potent degraders **20** and **21** in both the HiBit assay and western blot analysis resulting in increased levels of IDO1 protein at increased PROTAC concentrations. Interestingly, despite this, Kyn levels continue to decrease with increasing doses (Figure 5C and 5E). Furthermore, analysis of the parental BMS IDO1 inhibitor (**4**) and the E3 ligase dead analog of **21** (**24**, **NU227428**) shows no degradation of IDO1 in both the HiBit degradation assay and western blot analysis (Figure 5G).

**Figure 5.**
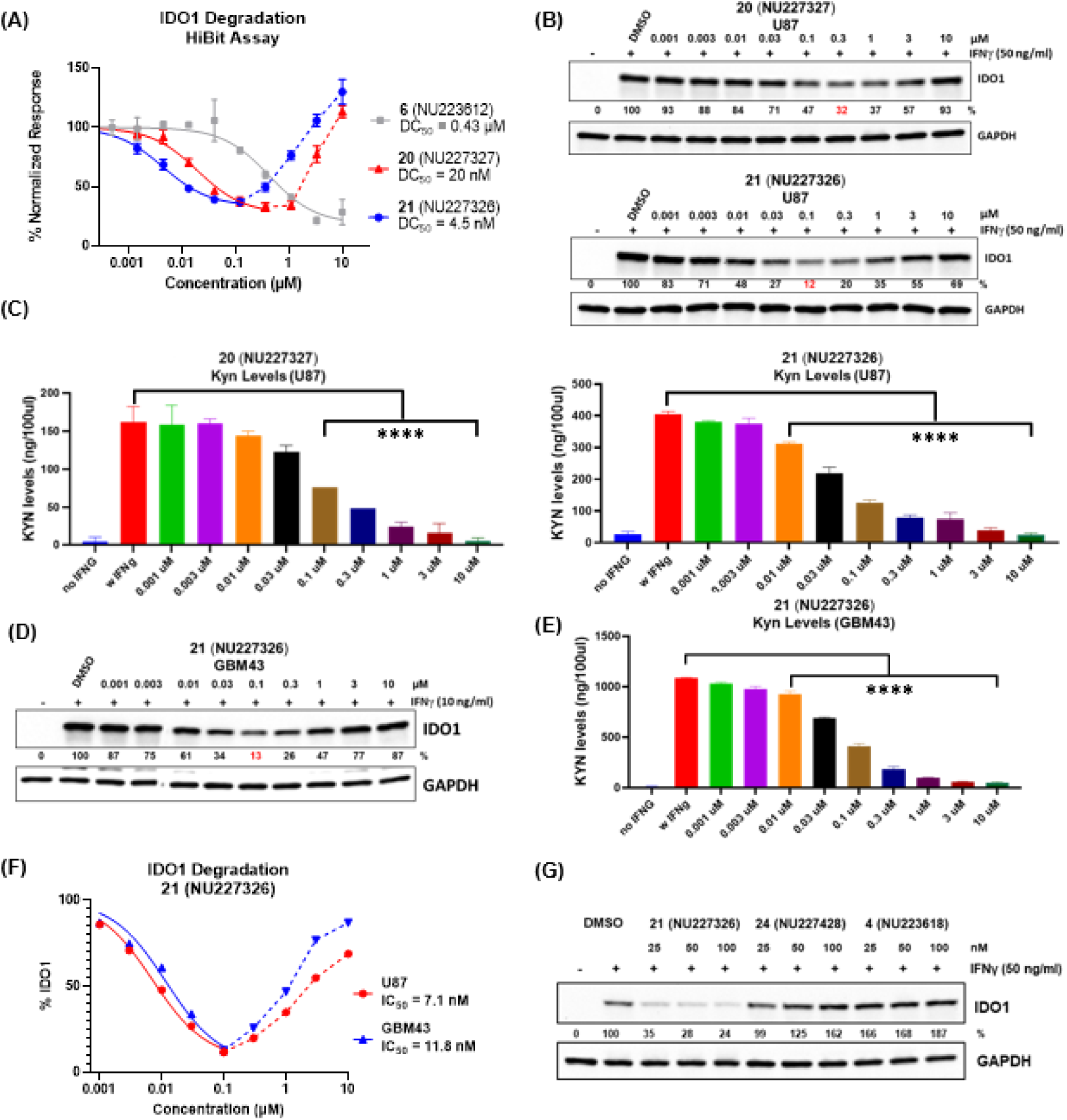
Characterization of IDO1 degradation by PROTACs **20** and **21**. (A) Degradation of IDO1 by **20**, **21**, and **NU223612** in the HiBiT assay. DC_50_ was calculated from the lowest concentration down to where maximum degradation was observed and excluded data points within the Hook effect range. R^2^ = 0.94 for **20**; R^2^ = 0.93 for **21**. (B) Dose-dependent degradation by compounds **20** and **21** on IDO1 in U87 cells treated with an extended dose range of **20** or **21** for 24 h, and protein samples were analyzed by western blotting. (C) Effects of **20** and **21** on kynurenine production in U87 human GBM cells. Cells were treated with **20** or **21** in the presence of 100 ng/mL IFNγ beginning at 24 hours after plating the cells and for a total of 24 h, followed by the collection of cell culture supernatants to measure IFNγ-induced kynurenine levels using Ehrlich’s reagent. Cells cultured in the absence of IFNγ served as the control group. (D) GBM43 cells were treated with an extended dose range of **21** for 24 h and protein samples were analyzed by western blotting. (E) Effect of **21** on kynurenine production in GBM43 cells. Cells were treated with **21** in the presence of 100 ng/mL IFNγ for 24 h and cell culture supernatants were collected to measure IFNγ-induced kynurenine levels using Ehrlich’s reagent. Cells cultured in the absence of IFNγ served as the control group. (F) Representative curves of percent IFNγ-induced IDO1 levels that were normalized to untreated samples as calculated in U87 and GBM43 cells to determine DC_50_ that produces 50% of IDO1 degradation (n = 3 independent experiments). Data are presented as mean ± SEM. Statistical significance was determined using Tukey’s multiple comparison test for comparisons between more than two groups. Significance levels are indicated as follows: *P* < 0.05 (*), *P* < 0.01 (**), *P* < 0.001 (***), and *P* < 0.0001 (****). (G) Western blotting analysis of IDO1 protein in U87 cells. IDO1 was induced in U87 cells with 50 ng/mL human IFNγ for 24 h followed by a 25, 50 or 100 nM treatment with IDO1 PROTAC (**21**, **NU227326**), inactive PROTAC (**24**, **NU227428**) or IDO1 enzyme inhibitor (**4**, **NU223618**) for 24 h before protein samples were prepared for western blotting analysis. Percent of normalized IDO1 protein levels were derived from densitometric analysis of band intensities. Western blotting results are representative of outcomes from 3 separate experimental replicates.

### Mechanism and Validation of IDO1 Degrader 21 (NU227326)

The kinetics of protein degradation by IDO1 PROTAC shows that **21** (**NU227326**) begins initial active degradation between 8 and 16 hours (Figure 6A). In evaluating the sustained degradation potential of analog **21** (**NU227326**), we subjected cells to continuous treatments of 50 and 100 nM of **21 (NU227326)** (Figure 6B). Remarkably, the impact endured for up to 72 hours. Notably, even after a 24-hour pulse treatment with subsequent withdrawal, the profound effect of **21** (**NU227326**) persisted significantly for at least 48 hours post washout (Figure 6C). This observation suggests that the pharmacological effects of degrader **21** (**NU227326)** will exhibit enduring efficacy beyond the initial administration.

**Figure 6.**
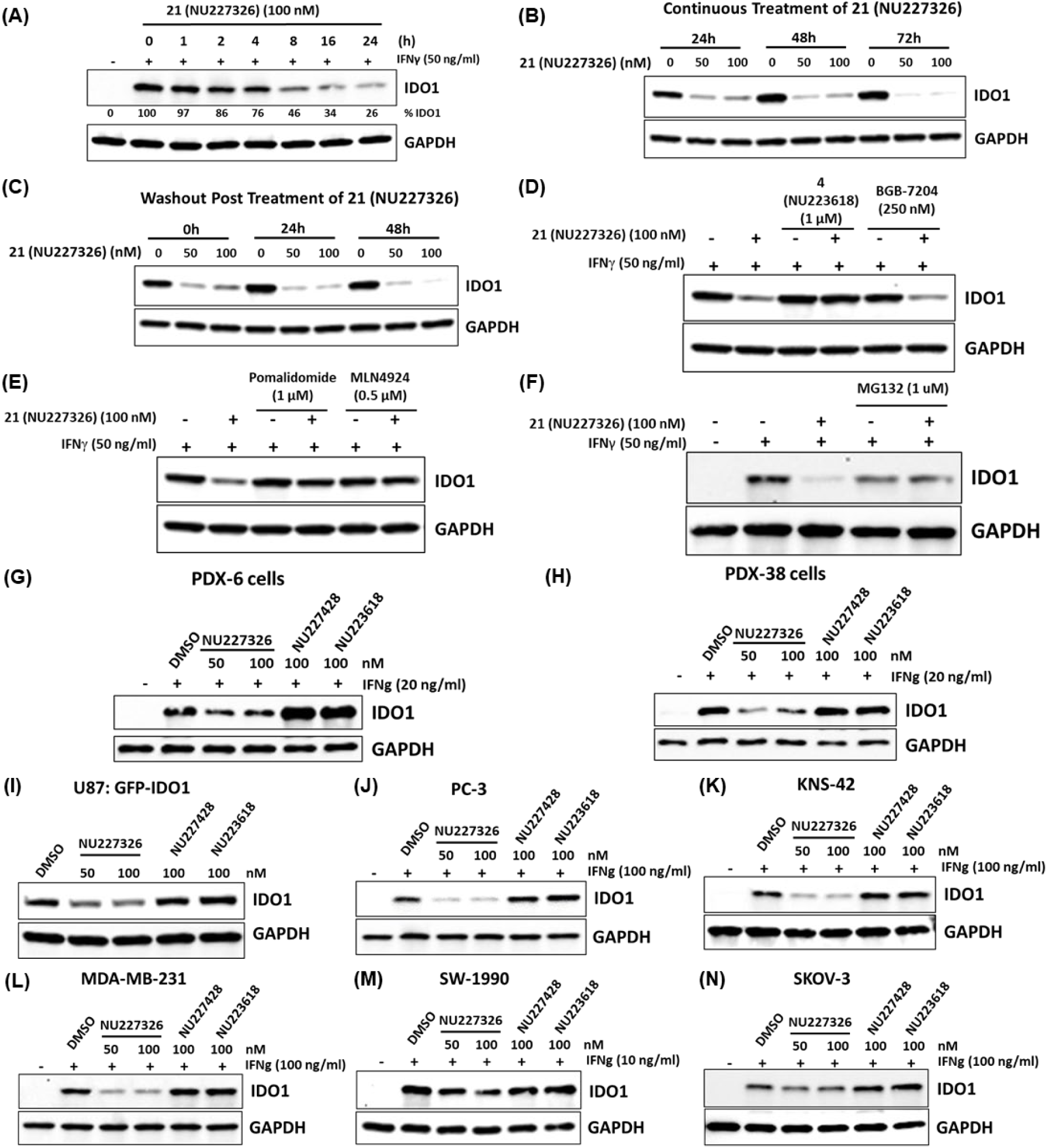
Characterization of **21** (**NU227326**) as a potent IDO1 PROTAC. (A) Western blot analysis of IDO1 and GAPDH to determine the kinetics of **21** (**NU227326**)-induced IDO1 protein degradation in U87 cells. (B) Western blot analysis of IDO1 to determine the effect of a single continuous treatment of **21** (**NU227326**) on IDO1 protein levels at multiple time points in U87 cells. (C) Similar to (B), U87 cells were treated with **21** (**NU227326**) for 24 h, and cells were cultured without **21** (**NU227326**) for up to 48 h. Protein samples were tested in western blot analysis to determine IDO1 levels upon withdrawal of the IDO1 PROTAC. (D) Western blotting analysis of IDO1 and GAPDH to determine the effect of parental competitive IDO1-inhibitor (**4**, **NU223618**) and non-competitive IDO1-inhibitor (BGB-7204); (E) E3 ligase ligand (pomalidomide) and E1 ligase neddylation inhibitor (MLN4924); and (F) proteasome inhibitor (MG132) on NU227326-induced IDO1 degradation. Western blotting analysis to determine the effect of **NU227326** on IFNγ-induced IDO1 levels in human adult PDX cells; PDX-6 (G) and PDX-38 (H), U87 cells exogenously overexpressing GFP-fused IDO1 (I), prostate cancer PC-3 cells (J), human pediatric KNS42 GBM cells (K), triple negative breast cancer MDA-MB-231 cells (L), pancreatic cancer SW-1990 cells (M), and ovarian SKOV-3 cells (N). Cells treated with either **21** (**NU227326), 24 (NU227428),** or **4 (NU223618)** at indicated concentrations for 24 hours and protein samples were analyzed for IDO1 and GAPDH using Western blotting analysis. Western blotting results are representative of outcomes from 3 separate experimental replicates.

To study the basic principle of the PROTAC’s mechanism for protein degradation, we sought to assess the protein degradation capability of compound **21** (**NU227326**). Treatment of U87 cells with a combination of **21** (**NU227326**) and the competitive IDO1 enzyme inhibitor **4** (**BMS-986205**, **NU223618**) markedly reduced IDO1 protein degradation as compared to the treatment of **21** (**NU227326**) alone (Figure 6D). This effect was conspicuously absent when U87 cells were treated with the combination of IDO1 PROTAC **21** (**NU227326**) and the non-competitive IDO1 enzyme inhibitor, BGB-7204^34^ (Figure 6D). This implies that the binding site occupied by the competitive inhibitor hinders the degradation ability of IDO1 PROTAC **21** (**NU227326**).

Analogously, the administration of an excess of the E3 ligase ligand, pomalidomide, hinders the binding of **21** (**NU227326**) to the E3-ligase CRBN and eliminates the degrader’s ability to degrade the target protein (Figure 6E). Introducing disruption to the multimeric E3 ubiquitin ligase complex through the addition of the E1 (ubiquitin-activating enzyme) ligase inhibitor, MLN4924,^46^ which renders the CRL complex (CRL4^CRBN^) inactive and disrupts CRBN-cullin-RING ligase-mediated protein turnover, resulted in the accumulation of IDO1 protein compared to treatment with **21** (**NU227326**) alone (Figure 6E). Furthermore, treatment with proteasome inhibitor MG132 in combination with degrader **21** (**NU227326**) results in no degradation of IDO1 protein (Figure 6F). Collectively, these findings support the necessity of the CRL4^CRBN^-mediated E3 ubiquitin ligase complex for the effective degradation of targeted protein substrates by IDO1 PROTAC **21** (**NU227326**). To further investigate the broad degradation potential of compound **21** (**NU227326**), we extended our testing to additional glioblastoma (GBM) models including two patient-derived xenograft (PDX) lines, PDX-6 (Figure 6G) and PDX-38 (Figure 6H), the pediatric GBM cell line, KNS-42 (Figure 6K), as well as a genetically-engineered U87 line expressing an IDO1 cDNA fused to GFP (Figure 6I). We also explored the efficacy of **21** (**NU227326**) across a wide range of human tumor types including pancreatic-, triple-negative breast-, ovarian-, and prostate-cancers (Figures 6J, 6L-6M). Compound **21** (**NU227326**) consistently demonstrated potent, dose-dependent degradation of IDO1, whereas neither the inactive PROTAC **24** (**NU227428**) nor the parental compound **4** (**NU223618**) elicited comparable effects. The selective activity of Compound **21** (**NU227326**) reinforces its specificity and generalizability for targeting IDO1 across multiple tumor types and confirms its reproducible efficacy in both GBM and other solid tumors (Figure 6G–6N). Strikingly, we observed a dose-dependent reduction in kynurenine levels across all tested cell lines after treatment with **NU227326**, indicating that Compound **21** is also able to influence IDO1 enzyme activity (**Supplementary** Figure 1). These findings highlight the broad and consistent activity of Compound **21** (**NU227326**) and supports its potential as a targeted therapeutic across a spectrum of IDO1-driven cancers.

### Proteomics of NU227236 and NU227327

To define the selectivity of our lead degraders and identify any other proteins that they degrade, they were tested in global quantitative proteomics. Each compound was tested at 1μM for 24 hrs. in Kelly neuroblastoma cells without induction by IFN-γ in order to assess unperturbed cells. As these cells do not endogenously express IDO1, mass spectrometry did not detect these proteins. Treatment with either compound only resulted in a significant decrease of 4-5 proteins. With both compounds, proteins ADO, EPHX2, CDK8, and SALL4 were found to be decreased (**Supplementary** Figure 2). Compound NU227326 also degrades ITCH.

### Characterization of IDO1 binding

Intrigued by the *in vitro* degradation profiles, **20 (NU227327)** and **21 (NU227326)** were further assessed using bio-layer interferometry (BLI) to measure their ability to form binary and ternary complexes with recombinant IDO1 and CRBN proteins in vitro. Bio-layer interferometry is a label-free optical technique used to analyze bimolecular interactions in real time.^47^ Both **20** and **21** were confirmed to directly bind to CRBN, possessing binding affinities (K_d_) of 520 nM and 380 nM, respectively (Figure 7A). In comparison to initial lead analog **6** (K_d_ = 290 nM) degraders **20** and **21** exhibited weaker binding to CRBN. As expected, no measurable binding to CRBN was observed for IDO1 inhibitor **4** (**NU223618**) lacking an E3 ligase ligand or inactive PROTAC **24** (**NU227428**) (**Supplementary** Figure 3).

**Figure 7.**
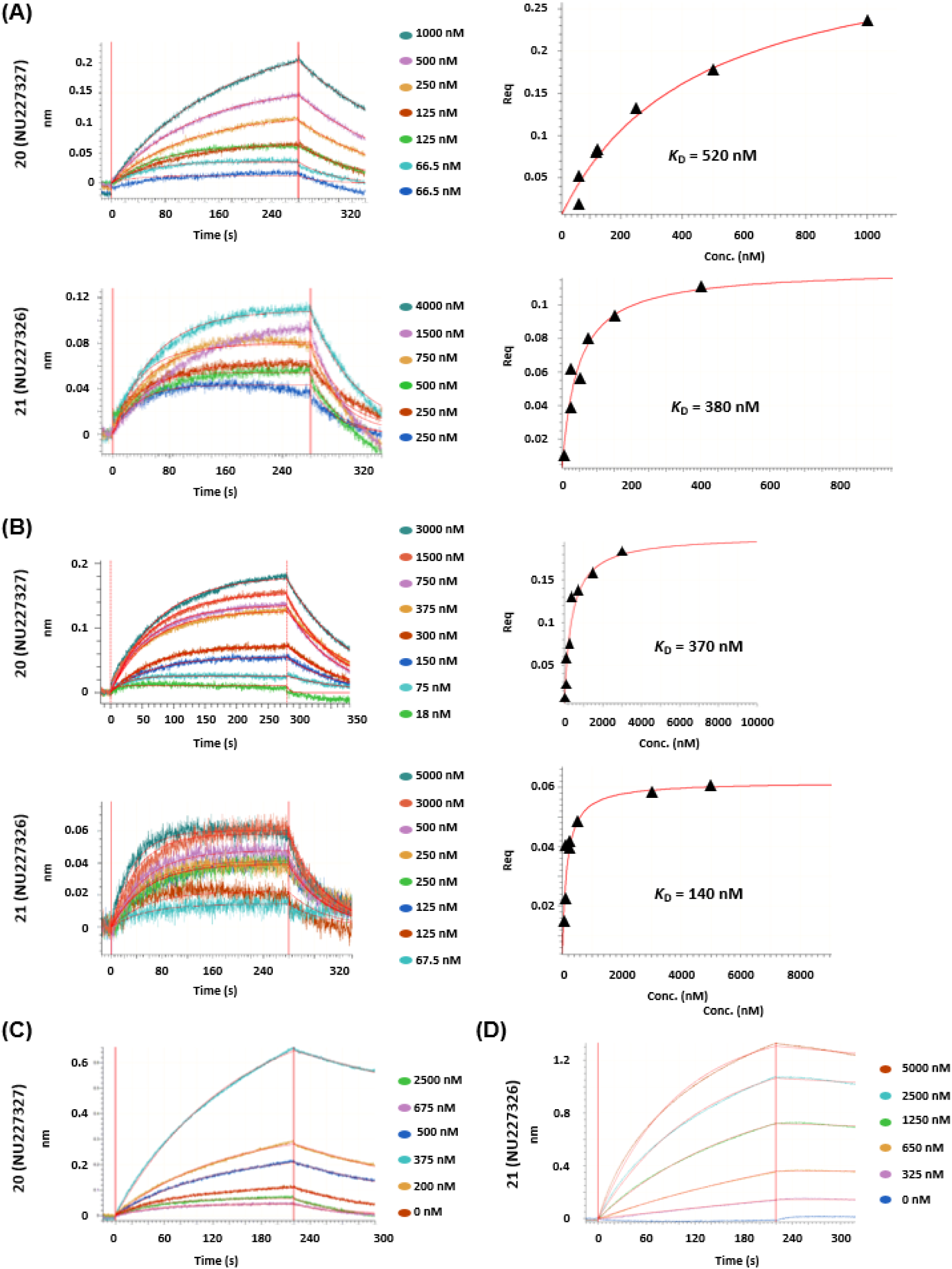
(A) Left: BLI sensorgrams showing association and dissociation of compounds **20** (**NU227327**) (upper) and **21** (**NU227326**) (lower) with CRBN (immobilized on AR2G sensors). Steady-state data fitted as described above yielded *K*_d_ values of 520 ± 106 nM and 380 ± 63 nM for the binary complexes formed by CRBN with **20** and **21**, respectively. (B) Left: BLI sensograms showing association and dissociation of compounds **20** (**NU227327**) (upper) and **21** (**NU227326**) (lower) to IDO1 protein (loaded on NI NTA sensors). Right: For each complex, the equilibrium dissociation constant (*K*_d)_ was obtained by fitting the steady state data (Req as a function of compound concentration) with a 1:1 binding model. The measured affinities for the binary complexes formed by IDO1 with compounds **20** and **21** were determined to be 370 ± 62 nM and 240 ± 49 nM, respectively. (C) BLI sensograms monitoring the association and dissociation kinetics of IDO1-**20**-CRBN (left) and IDO1-**21**-CRBN (right) ternary complexes. IDO1 protein was loaded on NiNTA biosensors. Double-referenced data sets fitted globally with a 1:1 kinetic model yielded *K*_d_ values of 321 ± 54 nM for IDO1-**21**-CRBN and 997 ± 143 nM for IDO1-**20**-CRBN. The k_off_ values obtained from the kinetic fits translate into the following lifetimes (t_1/2_): IDO1-**20** binary = 53 s, IDO1-**20**-CRBN ternary = 256 s.; IDO1-**21** binary = 38.6 s, IDO1-**21**-CRBN ternary =1039 s.

Analysis of PROTAC binding to IDO1 revealed **20** and **21** possessed stronger interactions compared to **6** (K_d_ = 640 nM), with binding affinities of 370 nM and 140 nM, respectively (Figure 7B). This enhancement in binding affinity to IDO1 could be attributable to both optimized PROTACs (**20** and **21**) possessing the 3-substituted versus the 4-substituted IDO1 ligand. The previously described docking studies provide evidence that this substitution position offers a more favorable exit vector orientation. Following similar procedures, the ternary IDO1-PROTAC-CRBN complex could be characterized. IDO1 degraders **20** and **21** were preincubated with an excess of CRBN to provide a PROTAC-CRBN binary complex. Binding of the resulting PROTAC-CRBN binary complex to IDO1 was detected and measured in real-time using BLI (Figure 7C). Interestingly, degrader **21** possessed a stronger ternary complex compared to degrader **20**, with binding affinities of 321 nM and 997 nM, respectively. Analog **21** also exhibited a more stable ternary interaction as evidenced by a longer half-life (t_1/2_) of 1039 seconds compared to a 256 second half-life for analog **20**. This enhancement in ternary complex formation and stability for degrader **21** over degrader **20** may be attributable to the loss in flexibility of the linker moiety via loss of a methylene unit between the piperazine and piperidine ring systems. Furthermore, the more desirable ternary complex interaction of **21** is consistent with the slightly increased potency of **21** compared to **20**.

### Pharmacokinetics of 21 (NU227326)

To characterize its suitability for future use in *in vivo* experiments, we measured the mouse pharmacokinetics of degrader **21** (**NU227326**) in plasma and brain tissue. After identifying 50 mg/kg as the maximum dose that could be administered without producing clinical signs of toxicity, 50 mg/kg was dosed intraperitoneally (IP) in mice and the resulting plasma and brain homogenate drug concentrations were measured (**Figure 8**). Initial studies tested **NU227326** for oral bioavailability using oral gavage administration but found very little plasma exposure (data not shown). Degrader **NU227326** was found to have a favorable half-life (5.7 hrs) in plasma and was retained in brain tissue for approximately twice as long (t_1/2_ = 11.8 hrs). Similar to virtually all other PROTACs described to date, **NU227326** has a limited ability to cross the blood-brain barrier (BBB). Total drug exposure in the brain was 1.0 μM·h, with a C_max_ of 0.1 μM, which indicated 4% of drug was able to cross the BBB. We also tested the PK of compound **NU227327** (**20**) and found it had lower plasma exposure, but a greater ability to cross the BBB, with total brain exposure being 10% of that in plasma (**Supplementary Figure 4**). Both compounds were >99% plasma protein bound as measured by a 6 hr rapid equilibrium dialysis.

**Figure 8.**
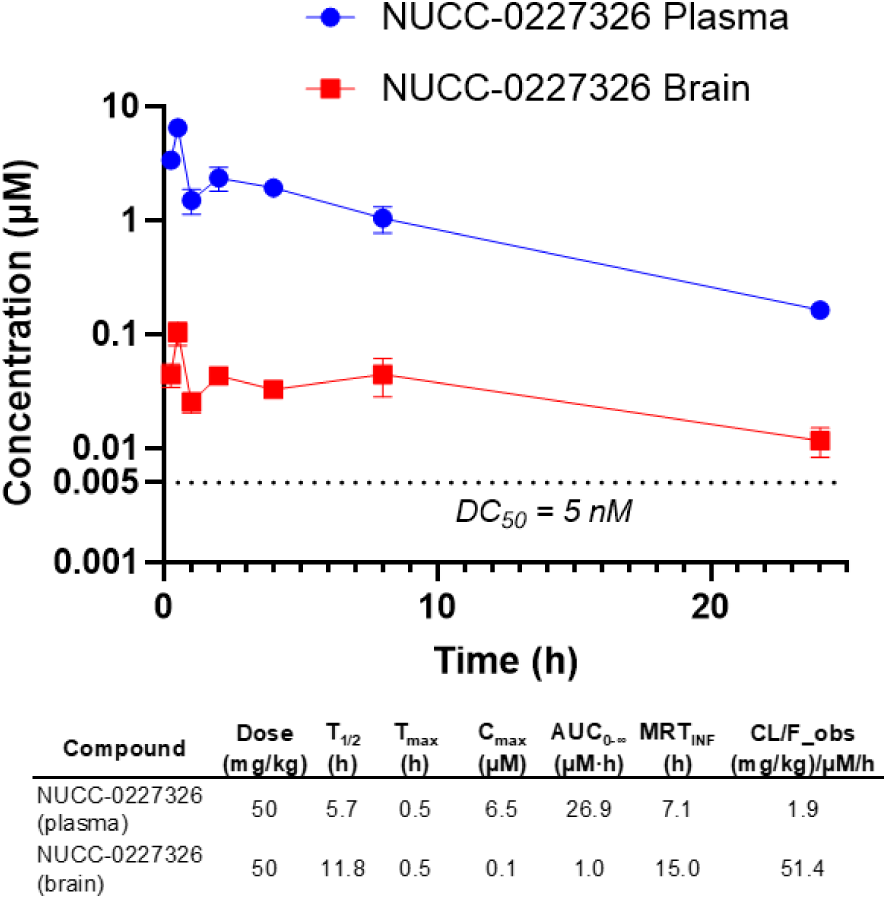
Pharmacokinetics of **NU227326** (**21**) in mice. Compound was dosed at 50 mg/kg by IP in mice.

### Chemistry

IDO1 PROTACs **6** – **23** were all designed to embody a piperidine ring unit as an attachment anchor for the linker to the IDO1 ligand. Therefore, to provide an efficient route towards diversification of the degrader structures, the synthesis of piperidine-containing IDO1 ligands to provide an amine building block for amide coupling was of interest. This strategy would allow for early-stage functionalization of the E3 ligase ligand with various linker motifs, simplifying the syntheses of the new IDO1 degraders.

To synthesize both the 4-chloro-3-substituted and 3-susbtituted type IDO1 ligand building blocks, alkylation of phenol **25** or **26** with *tert*-butyl 4-((methylsulfonyl)oxy)piperidine-1-carboxylate under basic conditions provided aryl nitro intermediates **27** and **28** in moderate to good yields, 90% and 60% respectively (Scheme 1). Selective reduction of the aryl nitro groups via catalytic hydrogenation using palladium on carbon afforded aryl amines **29** and **30**, which were taken forward without further purification. Both intermediates were reacted with previously described (*R*)-2-((1*S*,4*S*)-4-(6-fluoroquinolin-4-yl)cyclohexyl)propanoic acid^48^ under amide coupling EDCI conditions followed by acid-mediated Boc-removal to provide 4-chloro-3-substituted (**31**) and 3-susbtituted (**32**) piperidine analogs.

**Scheme 1.**
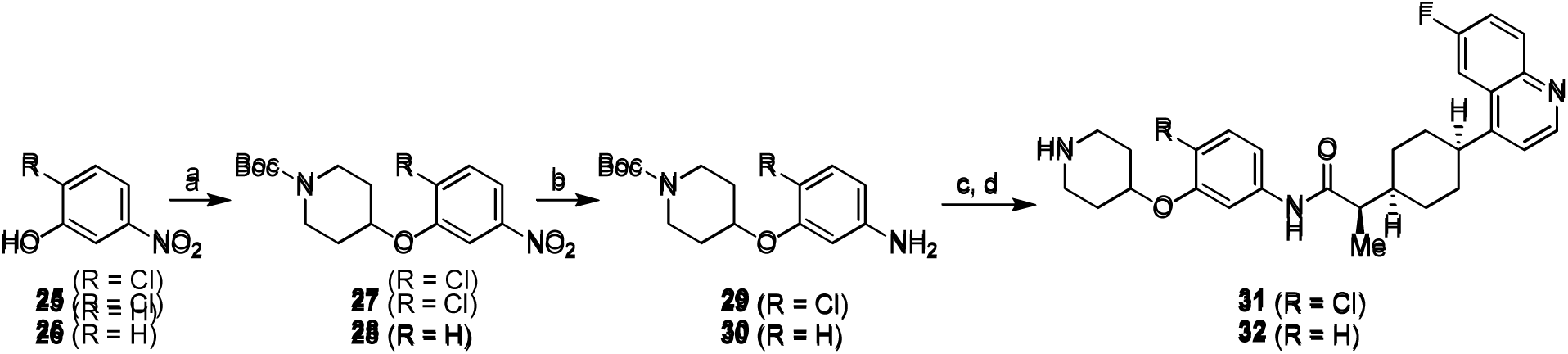
Synthesis of 4-chloro-3-substituted and 3-substitutied piperidine IDO1 binder building blocks 31 and 32^a^. *^a^*Reagents and conditions: (a) *tert*-butyl 4-((methylsulfonyl)oxy)piperidine-1-carboxylate, K_2_CO_3_, DMF, 75 °C, 12 h; (b) Pd/C, H_2_ (g), EtOH, 23 °C, 24-36 h; (c) (*R*)-2-((1s,4*S*)-4-(6-fluoroquinolin-4-yl)cyclohexyl)propanoic acid, EDCI, pyridine, 0 °C to 23 °C, 12 h; (d) 4 N HCl in dioxane, 23 °C, 12 h.

With the IDO1 binder building blocks in hand, the bifunctionalized CRBN-linker components and subsequent final degraders were synthesized. Final targets, **7** – **9**, were synthesized by first performing a nucleophilic aromatic substitution between either 4-fluoro-or 5-fluroro-thalidomide, **33** and **34**, and amino-PEG_2_-acid-*tert*-butyl ester in the presence of DIPEA (Scheme 2). Following an acid-mediated removal of the *tert*-butyl group to afford carboxylic acids **35** and **36**, the desired bifunctionalized derivatives were coupled with the previously synthesized 4-substituted amine or the 4-chloro-3-substituted amine (**31)** to provide the corresponding final PROTACs.

**Scheme 2.**
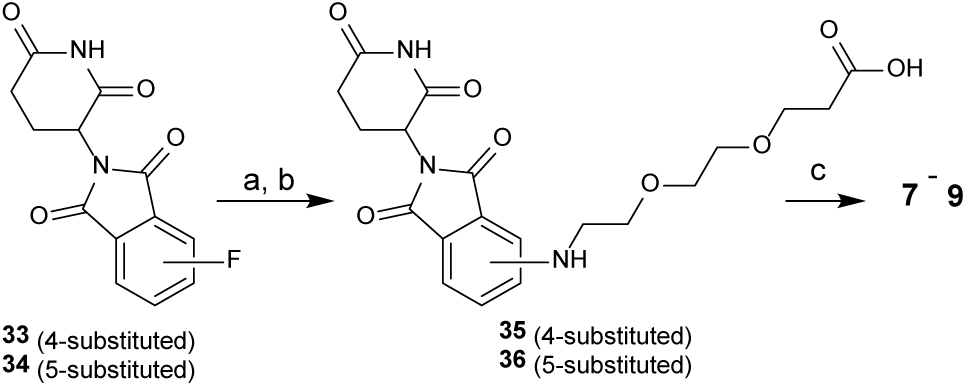
Synthesis of IDO1 degraders 7 – 9*^a^*. *^a^*Reagents and conditions: (a) *tert*-butyl 3-(2-(2-aminoethoxy)ethoxy)propanoate, DIPEA, DMF, 75 °C, 12-48 h; (b) CF_3_COOH, CH_2_Cl_2_, 0 °C to 23 °C, 1 h; (c) (R)-2-((1s,4S)-4-(6-fluoroquinolin-4-yl)cyclohexyl)-N-(4-(piperidin-4-yloxy)phenyl)propanamide or **31**, HATU, DIPEA, DMF, 0 °C to 23 °C, 6-12 h.

Introduction of the rigid linkers first involved implementing a piperazine ring connector to the E3 ligase ligand. Therefore, treatment of either **33** or **34** with *N*-Boc-piperazine under SnAr conditions followed by deprotection with TFA yielded 4-piperazine and 5-piperazine-thalidomide derivatives, **37** and **38**, in excellent yields (98% and 85%) (Scheme 3). Further functionalization of the piperazine intermediates via DIPEA promoted *N*-alkylation with *tert*-butyl 3-(2-iodoethoxy)propanoate and subsequent Boc removal afforded the desired acid intermediates, **39** and **40**. Amidation with the corresponding piperidine functionalized IDO1 ligands resulted in the formation of final degraders **10** – **13**. Incorporation of an entire rigid linker was achieved with further derivatization of thalidomide-piperazine intermediates, **37** and **38**. Therefore, through *N*-alkylation with *tert*-butyl 4-(bromomethyl)piperidine-1-carboxylate in the presence of DIPEA followed by acid mediated Boc removal, 4-and 5-thalidomide-piperazine-methylene-piperidine intermediates, **41** and **42**, respectively, were synthesized. Additionally, subjecting **38** to a reductive amination protocol with 1-Boc-4-piperidone and subsequent deprotection with TFA yielded 5-thalidomide-piperazine-piperidine intermediate, **43**. Piperidine analogs **41** – **43** were then treated with *tert*-butyl 2-bromoacetate and DIPEA to provide *N*-alkylated intermediates which upon acid-mediated removal of the *tert*-butyl group provided acids **44** – **46**. Amide coupling between the bifunctionalized acids with the previously described piperidine IDO1 intermediates provided the target IDO1 PROTACs **14** – **21**.

Truncated PROTACs, **22** and **23**, were synthesized according to Scheme 4. Utilizing the 5-piperazine-thalidomide derivative, **38**, *N*-alkylation with either *tert*-butyl 2-bromoacetate or *tert*-butyl 3-bromopropionate under basic conditions followed by deprotection to the acid yielded the corresponding intermediates, **47** and **48**. Subsequent amide coupling with the 3-substituted piperidine, **38**, resulted in the formation of target degraders **22** and **23**.

**Scheme 3.**
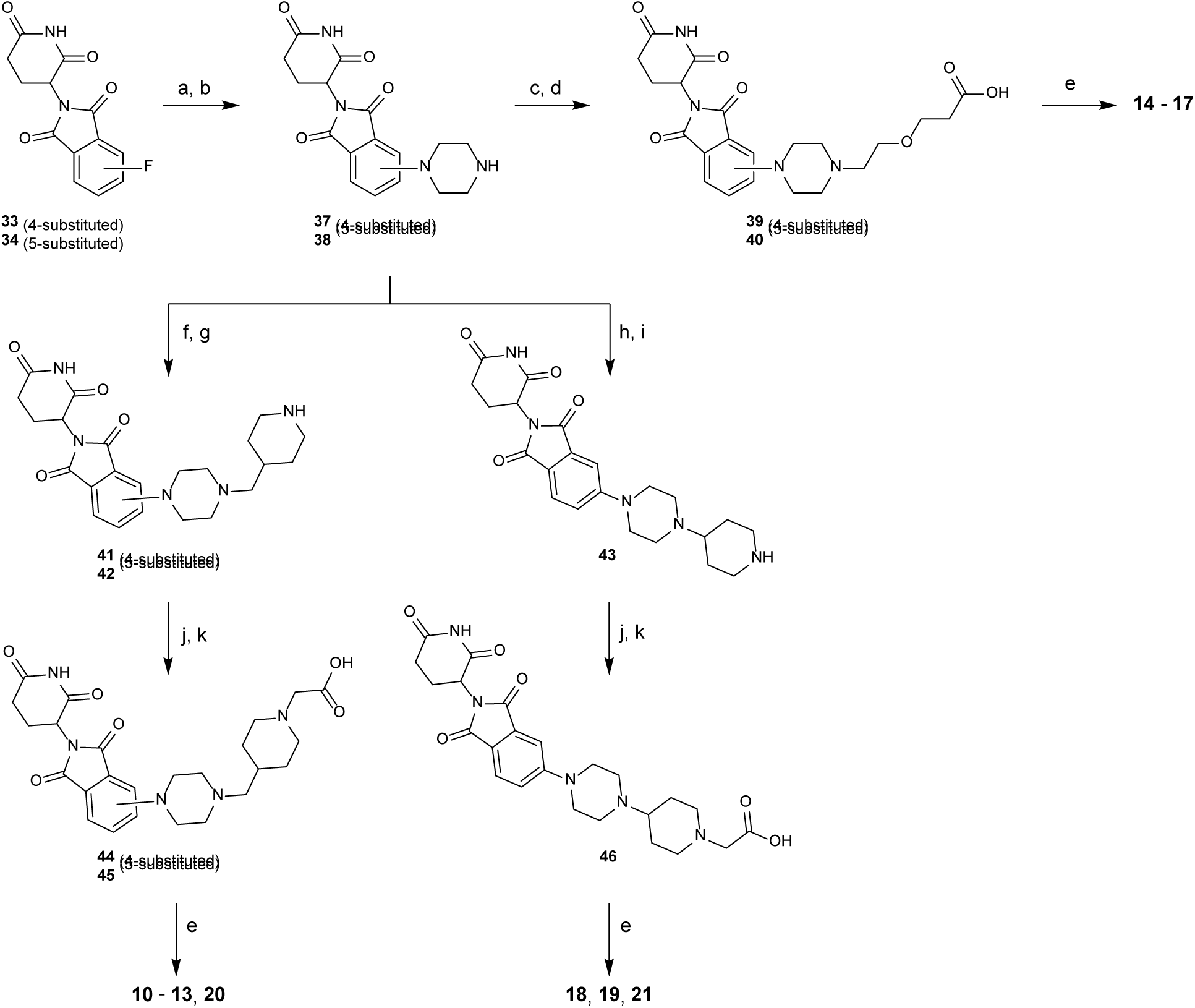
Synthesis of IDO1 degraders 10-21*^a^*. *^a^*Reagents and conditions: (a) *tert*-butyl piperazine-1-carboxylate, DIPEA, DMSO, 100 °C, 12 h; (b) CF_3_COOH, CH_2_Cl_2_, 0 °C to 23 °C, 1 h; (c) *tert*-butyl 3-(2-iodoethoxy)propanoate, DIPEA, DMF, 75 °C, 24 h; (d) CF_3_COOH, CH_2_Cl_2_, 0 °C to 23 °C, 1 h; (e) (c) (R)-2-((1s,4S)-4-(6-fluoroquinolin-4-yl)cyclohexyl)-N-(4-(piperidin-4-yloxy)phenyl)propanamide**, 31** or **32**, HATU, DIPEA, DMF, 0 °C to 23 °C, 1.5-24 h; (f) *tert*-butyl 4-(bromomethyl)piperidine-1-carboxylate, DIPEA, DMF, 75 °C, 12 h; (g) CF_3_COOH, CH_2_Cl_2_, 0 °C to 23 °C, 1-2.5 h; (h) *tert*-butyl 4-oxopiperidine-1-carboxylate, STAB, DMF, 23 °C, 2 h; (i) CF_3_COOH, CH_2_Cl_2_, 0 °C to 23 °C, 1 h; (j) tert-butyl 2-bromoacetate, DIPEA, DMF, 23 °C, 2-3 h; (k) CF_3_COOH, CH_2_Cl_2_, 0 °C to 23 °C, 2-4 h; (l) (c) (R)-2-((1s,4S)-4-(6-fluoroquinolin-4-yl)cyclohexyl)-N-(4-(piperidin-4-yloxy)phenyl)propanamide**, 31** or **32**, HATU or T3P, DIPEA, DMF, 0 °C to 23 °C, 1-48 h.

**Scheme 4.**
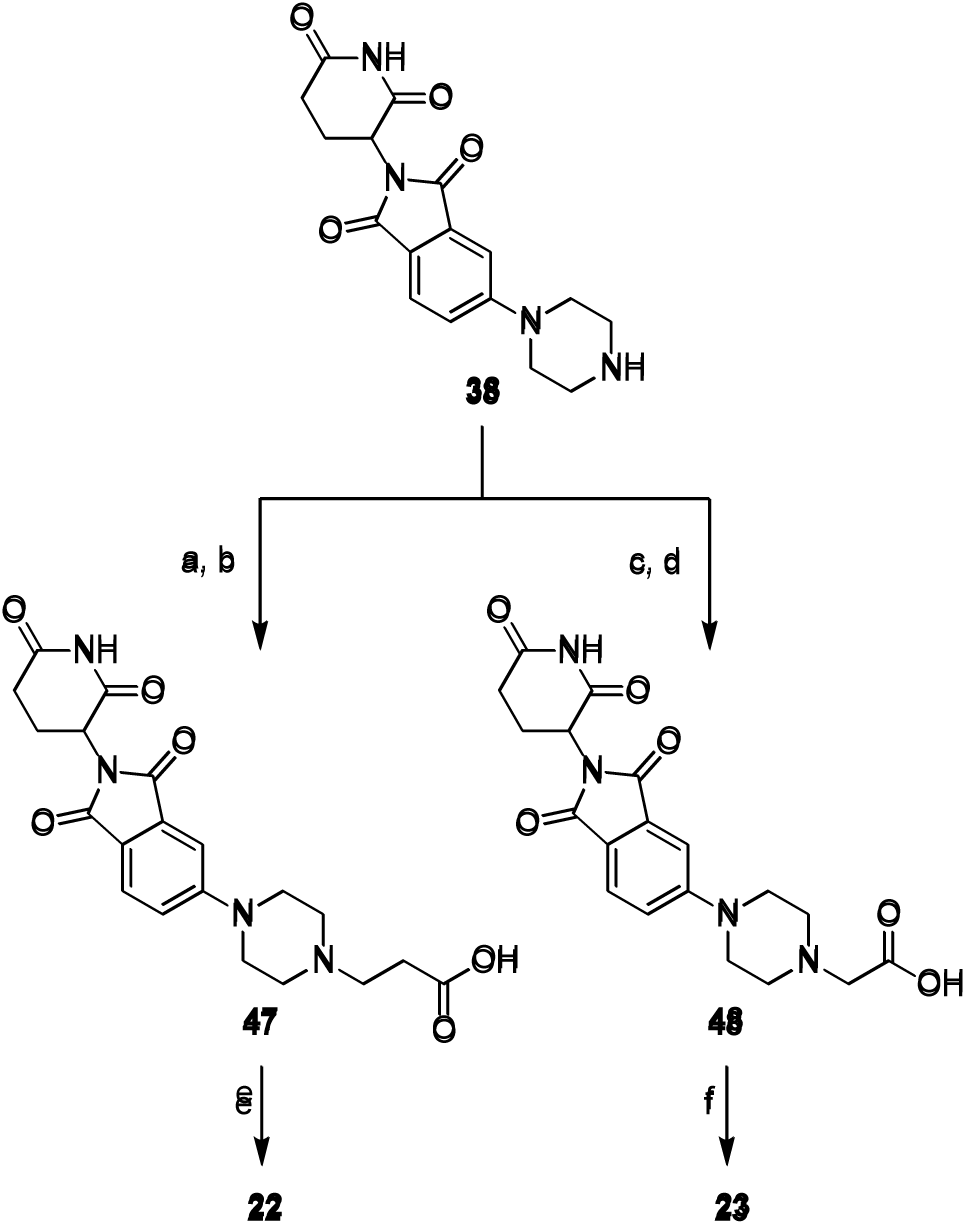
Synthesis of IDO1 degraders 22 and 23*^a^*. *^a^*Reagents and conditions: (a) *tert*-Butyl 3-bromopropionate, DIPEA, DMF, 23 °C, 12 h; (b) CF_3_COOH, CH_2_Cl_2_, 0 °C to 23 °C, 24 h; (c) *tert*-butyl 2-bromoacetate, DIPEA, DMF, 23 °C, 2 h; (d) CF_3_COOH, CH_2_Cl_2_, 0 °C to 23 °C, 24 h; (e) **37**, HATU, DIPEA, DMF, 0 °C to 23 °C, 30 min.

Synthesis of the inactive PROTAC, **24**, was achieved through methylation of the glutarimide nitrogen of the 5-fluoro-thalidomide starting material (**34**) with methyl iodide and potassium carbonate to provide **49** in 82% yield (Scheme 5). With *N*-Boc-piperazine, an SnAr reaction followed by acid-mediated deprotection yielded intermediate **50** in excellent yield. Reductive amination with 1-Boc-4-piperidone and sodium triacetoxyborohydride formed the desired piperazine-piperidine connectivity. Acid mediated Boc removal with TFA provided free amine **51**, which was alkylated with *tert*-butyl 2-bromoacetate under basic conditions. Subsequent cleavage of the *tert*-butyl group yielded carboxylic acid **52**. The final amide coupling with the previously synthesized 3-substituted piperidine intermediate (**32**) afforded the desired, inactive PROTAC **24**.

**Scheme 5.**
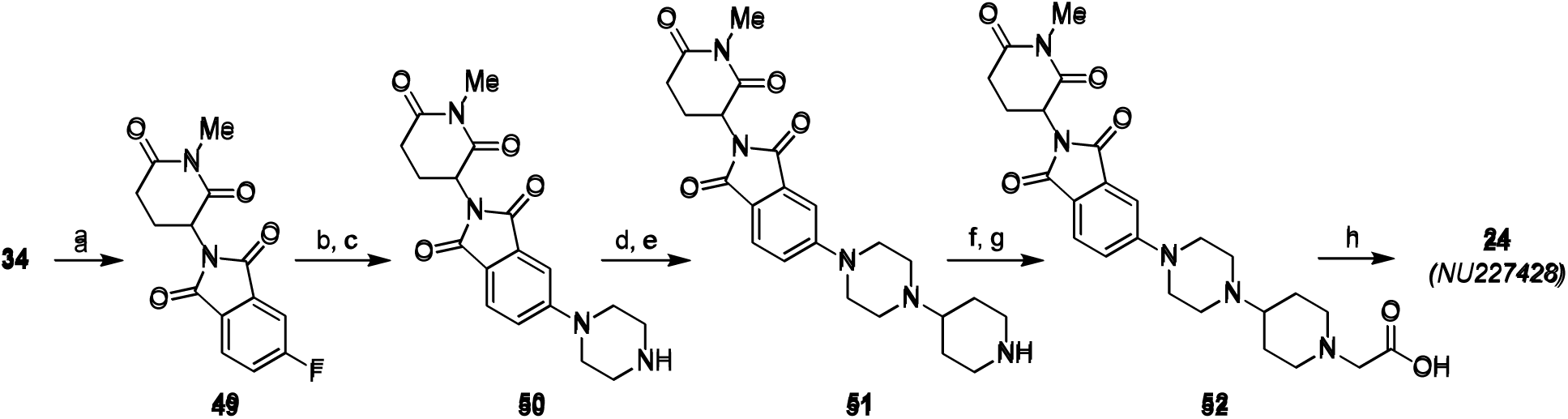
Synthesis of inactive IDO1 PROTAC 24*^a^*. *^a^*Reagents and conditions: (a) MeI, K_2_CO_3_, DMF, 0 °C to 23 °C, 12 h, 82%; (b) *tert*-butyl piperazine-1-carboxylate, DIPEA, DMSO, 100 °C, 2.5 h, 90%; (c) CF_3_COOH, CH_2_Cl_2_, 0 °C to 23 °C, 2 h, 90%; (d) *tert*-butyl 4-oxopiperidine-1-carboxylate, STAB, DMF, 23 °C, 1.5 h; (e) CF_3_COOH, CH_2_Cl_2_, 0 °C to 23 °C, 12 h, 77% over 2 steps; (f) *tert*-butyl 2-bromoacetate, DIPEA, DMF, 23 °C, 12 h; (g) CF_3_COOH, CH_2_Cl_2_, 0 °C to 23 °C, 4 h, 72% over 2 steps; (h) **32**, HATU, DIPEA, DMF, 0 °C to 23 °C, 30 min, 48%.

## DISCUSSION

Indoleamine 2,3 dioxygenase 1 (IDO1) is highly expressed in many intractable cancers including prostate, pancreatic, ovarian, colorectal, and glioblastoma (GBM).^49^ The relationship between IDO1 and its potent inhibition of the anti-cancer immune response is well-established.^12, 50^ The genetic knockdown of IDO1 in glioma cells leads to the spontaneous eradication of bulk tumor mass through a T cell-dependent mechanism.^18, 51^ ^18, 50^ Specifically, the genetic silencing of IDO1 results in a failure of immunosuppressive regulatory T cells (Tregs; CD3^+^CD4^+^CD25^+^FoxP3^+^) to accumulate in malignant glioma whereas cytolytic T cell (CD3^+^CD8^+^) infiltration is relatively unaffected by the genetic absence of tumor cell IDO1 expression. The higher CD8^+^ T:Treg ratio in the genetic absence of tumor cell IDO1 is associated with brain tumor eradication and long-term survival.^18^ While IDO1 is not normally expressed at high levels throughout a majority of tissues including the brain, human GBM-infiltrating T cells induce significant IDO1 expression in the bulk tumor mass.^52^ Furthermore, GBM cell IDO1 cDNA expression increases intratumoral Treg levels over that of GBM cells without forced IDO1 expression, and importantly, the IDO1-mediated recruitment of immunosuppressive Tregs is not reversed with the treatment of a potent IDO1 enzyme inhibitor.^34^ Similarly, glioma cell expression of both wild-type IDO1 cDNA leading to enzymatically active IDO1 protein expression, as well as, mutant IDO1 cDNA leading to enzymatically inactive IDO1 protein expression, similarly increase intratumoral Treg levels compared to the effects of IDO1-deficient glioma cells that express an empty cDNA vector.^25^ Based on the increased immunosuppression mediated by both enzymatically-active and enzymatically-inactive IDO1, we sought to develop IDO1 PROTACs to abolish all of these functions. We previously reported a first generation IDO1 PROTAC that possesses *in vivo* efficacy in a mouse model with intracranial GBM.^39^ That work motivated us to refine and optimize our IDO1 PROTAC into more potent and effective degraders.

Here, we describe our work in rationally designing and synthesizing a new series of potent IDO1 PROTACs. Using insights gained from molecular modeling studies, we designed new compounds that allowed more favorable binding to IDO1 and allowed more efficient access of the linker to the solvent. We also incorporated more rigid linkers to improve ternary complex formation and to increase pharmaceutical properties. Our improved compound described in this work, **NU227326** (**21**), has a 5 nM IDO1 DC_50_ in U87 cells and is similarly potent in PDX GBM43 cells. NU227326 further demonstrated generalizability for IDO1 protein degradation across many types of human cancers including additional human PDXs for GBM6 and GBM38, pediatric GBM KNS-42, prostate cancer line PC-3, breast cancer line MDA-MB-231, pancreatic cancer line SW-1990, as well as ovarian cancer line SKOV-3. The compound shows the expected reliance on CRBN and the ubiquitin-proteasome system for its degradation. We also found that **NU227326** achieves profound IDO1 degradation for at least 48 hours after washout from U87 cells and confirms a long-lasting effect. We also developed a HiBiT-based assay in U87 GBM cells to enable quantitative measurement of IDO1 degradation which provided orthogonal confirmation of **NU227326**’s potency. Compound **NU227326** was tested for mouse PK and we found it possessed favorable half-life and plasma exposure. It also crossed the BBB with 4% penetration. A closely related IDO1 degrader we describe here (**20**, **NU227327**) was found to achieve 10% brain penetration, suggesting this series of degraders has the ability for further structural optimization of brain exposure. Finally, to define the selectivity of our lead compounds and identify any off-targets that are degraded, we carried out global quantitative proteomics. Proteomics showed that both **NU227326** and **NU227327** significantly degraded <5 non-IDO1 proteins, indicating a high degree of selectivity. Notably, SALL4, which is a common cereblon neosubstrate, was degraded by both compounds.

Although the majority of our work focuses on the development of an IDO1 PROTAC for reversing tumor cell IDO1-mediated effects, it’s important to acknowledge that IDO1-expressing non-tumor cells also inhibit the antitumor immune response.^17^ In contrast to the tumor cell IDO1 effects that significantly increase Tregs, MDSCs, and are independent of treatment,^18^ it’s important to also recognize the role of non-tumor cell IDO1 that is inducibly relevant when immunotherapy is applied.^34^ This consideration is made further complicated by the role of non-tumor cell IDO1 effects that vary by subject age. The median age of an adult glioblastoma patient is 68-70 years old, which is roughly age-equivalent to a 22-24 month-old C57BL/6 mouse.^25, 53^ As we have previously demonstrated, old WT mice with intracranial malignant glioma lack a survival benefit after treatment with IDO1 enzyme inhibitor when coupled with radiation (RT) + PD-1 mAb. This outcome is dramatically different than the effect(s) of IDO1 enzyme inhibitor efficacy in younger WT mice with an intracranial brain tumor that experience an impressive survival benefit after combined IDO1 enzyme inhibitor + RT + PD-1 mAb treatment.^24^ Notably, although the IDO1 enzyme inhibitor fails to improve survival in older wild-type (WT) mice, similarly treated age-matched older adult IDO1KO mice demonstrate a significantly improved overall survival.^24^ This data does not suggest that the IDO1 enzyme inhibitor works better in older IDO1KO mice as compared to age-matched WT mice. Rather, it suggests that the genetic absence of IDO1 in non-tumor cells synergizes with RT + PD-1 mAb during old age and that this beneficial effect is not recapitulated with an IDO1 enzyme inhibitor – presumably due to IDO1 enzyme independent effects that are triggered during advanced age. Collectively, the data suggest that an IDO1 PROTAC targeting tumor cell-derived and non-tumor cell-derived IDO1 will possess a maximum net survival benefit for older adults with GBM. Ongoing work by our group aims to characterize the *in vivo* effects of our new IDO1 degrader **NU227326** on the tumor-and non-tumor immune response in immunocompetent GBM tumor models across the lifespan.

## EXPERIMENTAL SECTION

### General Chemical Methods

All chemical reagents were acquired from commercial suppliers and utilized without further purification. Reactions were completed without taking precautions to exclude air or moisture, unless otherwise noted. Normal-phase column chromatography was performed using silica gel columns and ACS grade solvents. ^1^H NMR and ^13^C NMR spectroscopy were recorded on Bruker 400 MHz or Bruker Avance III 500 MHz spectrometers. The chemical shifts for ^1^H NMR are reported to the second decimal place in parts per million (ppm). Proton coupling constants are expressed in hertz (Hz). Standard abbreviations were used to denote spin multiplicity for ^1^H NMR data. The chemical shifts for ^13^C NMR are reported to the first decimal place in ppm. The representative residual solvent peaks (CDCl_3,_^1^H *δ* = 7.27 ppm, ^13^C *δ* = 77.16 ppm; CD_3_OD-d_4_, ^1^H *δ* = 3.31 ppm, ^13^C *δ* = 49.00 ppm; DMSO-*d*_6_, ^1^H *δ* = 2.50 ppm, ^13^C *δ* = 39.52 ppm) were used as an internal standard. High-resolution mass spectrometry (HRMS) values were measured and calculated with an Agilent 6545 QTOF mass spectrometer coupled with an Agilent 1200 series LC, with direct loop injection (no column). All compounds presented in the manuscript are >95% pure by HPLC analysis.

### *tert*-butyl 4-(2-chloro-5-nitrophenoxy)piperidine-1-carboxylate (27)

To a solution of *tert-*butyl 4-((methylsulfonyl)oxy)piperidine-1-carboxylate (3.0 g, 0.01 mol, 2.0 equiv) and **25** (1.0 g, 0.01 mol, 2.0 equiv) in DMF (15 mL) was added K_2_CO_3_ (2.0 g, 0.01 mol, 2.0 equiv). The resulting mixture was stirred at 75 °C for 12 h. The reaction mixture was allowed to cool to 23 °C and then diluted with DI water (20 mL). The resulting mixture was extracted with EtOAc (3 × 20 mL), and the organic layers were combined, washed with brine (2 × 30 mL), dried over anhydrous MgSO_4_, filtered, and concentrated under reduced pressure. The crude product was purified by column chromatography (SiO_2_, hexanes/EtOAc, 100:0 to 50:50) to give **27** (1.9 g, 90%) as an off white solid. ^1^H NMR (500 MHz, CDCl_3_) δ 7.83 – 7.76 (m, 2H), 7.53 (d, *J* = 8.5 Hz, 1H), 4.69 (tt, *J* = 6.6, 3.4 Hz, 1H), 3.64 (ddd, *J* = 12.7, 8.5, 3.7 Hz, 2H), 3.52 (dd, *J* = 12.6, 6.9 Hz, 2H), 1.96 (td, *J* = 9.1, 4.3 Hz, 2H), 1.87 (ddt, *J* = 10.8, 7.1, 3.9 Hz, 2H), 1.48 (s, 9H). Spectral data matched previously reported characterization data.^54^

### *tert*-butyl 4-(3-nitrophenoxy)piperidine-1-carboxylate (28)

To a solution of *tert*-butyl 4-((methylsulfonyl)oxy)piperidine-1-carboxylate (4.0 g, 0.01 mol, 2.0 equiv) and **26** (1.0 g, 7.0 mmol, 1.0 equiv) in DMF (20 mL) was added K_2_CO_3_ (2.0 g, 0.01 mol, 2.0 equiv). The resulting mixture was stirred at 75 °C for 12 h. Upon completion of the reaction, the mixture was allowed to cool to 23 °C, diluted with DI water, and extracted with EtOAc (3 × 15 mL). The organic layers were combined, washed with brine (1 × 30 mL), dried over anhydrous MgSO_4_, and concentrated *in vacuo.* The crude residue was purified by column chromatography (SiO_2_, hexanes/EtOAc, 100:0 to 50:50) to yield **28** (1.5 g, 60 %) as a yellow oil. Spectral data matched previously reported characterization data.^55^

### *tert*-butyl 4-(5-amino-2-chlorophenoxy)piperidine-1-carboxylate (29)

To a solution of **27** (850 mg, 2.38 mmol, 1.0 equiv) in EtOH (30 mL) was added Pd/C (12.7 mg, 10% Pd on charcoal, wet, containing 50% H_2_O) (catalytic) at 23 °C. The reaction was fitted with a hydrogen balloon and allowed to stir at 23 °C for 36 h. The resulting mixture was filtered through a celite pad with EtOH rinses and the filtrate was concentrated to yield **29** (720 mg, 92%) as a yellow oil. Spectral data matched previously reported characterization data.^54^

### *tert*-butyl 4-(3-aminophenoxy)piperidine-1-carboxylate (30)

To a solution of **28** (1.5 g, 4.7 mmol, 1.0 equiv) in EtOH (4 mL) was added Pd/C (50 mg, 10% Pd on charcoal, wet, containing 50% H_2_O) (catalytic) at 23 °C. The reaction mixture was fitted with a hydrogen gas balloon and allowed to stir at 23 °C for 24 h. The resulting mixture was filtered through a celite pad with EtOH rinse, and the filtrate was concentrated under reduce pressure to provide **28** (1.0 g, 70 %) as a yellow oil. Spectral data matched previously reported characterization data.^56^ The product was taken forward without further purification.

### (*R*)-*N*-(4-chloro-3-(piperidin-4-yloxy)phenyl)-2-((1s,4*S*)-4-(6-fluoroquinolin-4-yl)cyclohexyl)propenamide (31)

To a solution of **29** (4.0 g, 0.01 mol, 1.0 equiv) and (*R*)-2-((1s,4*S*)-4-(6-fluoroquinolin-4-yl)cyclohexyl)propanoic acid (3.0 g, 0.01 mmol, 1.0 equiv) in pyridine (20 mL) was added EDCI (3 g, 0.01 mmol, 1.5 equiv) at 0 °C. The resulting mixture was allowed to stir at 23 °C for 12 h. The reaction was poured into 1N HCl (100 mL) and extracted with EtOAc (3 × 75 mL). The combined organic layers were washed with 1N HCl (3 × 30 mL), Sat Na_2_CO_3_ solution (3 × 50 mL), dried over anhydrous MgSO_4_, filtered and concentrated under reduced pressure to provide *tert*-butyl 4-(2-chloro-5-((*R*)-2-((1*s*,4*S*)-4-(6-fluoroquinolin-4-yl)cyclohexyl)propanamido)phenoxy)piperidine-1-carboxylate as an orange residue. The resulting product was taken forward without additional purification. To a solution of the crude *tert*-butyl 4-(2-chloro-5-((*R*)-2-((1*s*,4*S*)-4-(6-fluoroquinolin-4-yl)cyclohexyl)propanamido)phenoxy)piperidine-1-carboxylate in 1,4-dioxane (5 mL) was added a solution HCl/Dioxane (10 mL, 4 molar). The resulting mixture was allowed to stir at 23 °C overnight. Upon reaction completion, the mixture was diluted with cold Et_2_O and the solid precipitate was collected via vacuum filtration to yield **31** (4 g, 70% over two steps) as a dark yellow solid. ^1^H NMR (DMSO, 500 MHz): δ = 10.77 (s, 1H), 9.22 (d, 1H, *J*=4.4 Hz), 9.08 (s, 1H), 8.53 – 8.17 (m, 3H), 8.04 (dt, 1H, *J*=26.6, 7.8 Hz), 7.85 (t, 1H, *J*=1.4 Hz), 7.45 – 7.14 (m, 2H), 4.62 (tt, 1H, *J*=6.9, 3.2 Hz), 3.68 – 3.57 (m, 1H), 3.23 – 3.17 (m, 2H), 3.11 (d, 3H, *J*=11.8 Hz), 2.29 – 2.08 (m, 3H), 1.98 – 1.57 (m, 10H), 1.12 (d, 3H, *J*=6.6 Hz) ppm. ^13^C NMR (126 MHz, DMSO) δ 175.4, 162.0, 160.0, 151.8, 145.7, 142.4, 139.6, 130.0, 128.1, 128.0, 127.1, 122.8, 122.6, 120.0, 116.4, 113.2, 108.8, 108.6, 107.4, 70.7, 48.6, 38.6, 35.5, 28.4, 27.8, 27.1, 27.0, 26.9, 26.2, 16.3 ppm. HRMS (*m/z*): [M + H]^+^ calcd. for C_29_H_34_ClFN_3_O_2_ 510.2324; found, 510.2337.

### (*R*)-2-((1s,4*S*)-4-(6-fluoroquinolin-4-yl)cyclohexyl)-*N*-(3-(piperidin-4-yloxy)phenyl)propenamide (32)

To a solution of 30 (1.0 g, 4.0 mmol, 1.0 equiv) and (*R*)-2-((1s,4*S*)-4-(6-fluoroquinolin-4-yl)cyclohexyl)propanoic acid (1.0 g, 3.0 mmol, 1.0 equiv) in pyridine (15 mL) was EDCI (1.0 g, 5.0 mmol, 1.5 equiv) at 0 °C. The resulting mixture was allowed to stir at 23 °C overnight. The resulting mixture was poured into 1N HCl (75 mL) and extracted with EtOAc (3 × 50 mL). The organic layers were combined, washed with 1N HCl (3 × 30 mL), Sat Na_2_CO_3_ solution (3 × 50 mL), dried over anhydrous MgSO_4_, filtered, and concentrated *in vacuo* to a provide a brown residue. The crude mixture was purified by column chromatography (SiO_2_, hexanes/EtOAc, 100:0 to 50:50) to provide *tert*-butyl 4-(3-((*R*)-2-((1*s*,4*S*)-4-(6-fluoroquinolin-4-yl)cyclohexyl)propanamido)phenoxy)piperidine-1-carboxylate (700 mg, 40%) as a yellow solid. ^1^H NMR (500 MHz, CDCl_3_) δ 8.76 (d, *J* = 4.6 Hz, 1H), 8.10 (dd, *J* = 9.2, 5.6 Hz, 1H), 8.04 (s, 1H), 7.63 (dd, *J* = 10.5, 2.8 Hz, 1H), 7.51 – 7.40 (m, 2H), 7.28 (d, *J* = 4.6 Hz, 1H), 7.17 (t, *J* = 8.1 Hz, 1H), 7.01 (d, *J* = 8.0 Hz, 1H), 6.68 – 6.58 (m, 1H), 4.45 (dt, *J* = 7.0, 3.6 Hz, 1H), 3.64 (ddd, *J* = 13.4, 7.7, 3.7 Hz, 2H), 3.31 (ddt, *J* = 16.0, 8.0, 3.9 Hz, 3H), 2.63 (dd, *J* = 10.9, 6.7 Hz, 1H), 2.18 – 2.09 (m, 1H), 1.92 – 1.58 (m, 12H), 1.46 (s, 9H), 1.29 – 1.24 (m, 3H) ppm. To *tert*-butyl 4-(3-((*R*)-2-((1*s*,4*S*)-4-(6-fluoroquinolin-4-yl)cyclohexyl)propanamido)phenoxy)piperidine-1-carboxylate (700 mg, 1.22 mmol, 1.0 equiv) was added a solution of TFA:CH_2_Cl_2_ (1:1, 3 mL) at 0 °C. The resulting mixture was stirred at 23 °C for 2 h. The reaction mixture was concentrated under nitrogen atmosphere and the crude residue was triturated with cold Et_2_O (2 × 15 mL) to provide pure 32 (650 mg, 90.8%) as an orange solid. ^1^H NMR (500 MHz, DMSO-*d*_6_) δ 10.07 (s, 1H), 9.00 (d, *J* = 4.9 Hz, 1H), 8.69 (d, *J* = 36.0 Hz, 2H), 8.16 (ddd, *J* = 32.8, 10.0, 4.3 Hz, 2H), 7.79 (ddd, *J* = 22.5, 11.2, 3.8 Hz, 2H), 7.48 (s, 1H), 7.29 – 7.05 (m, 2H), 6.78 – 6.59 (m, 1H), 4.59 (tt, *J* = 7.1, 3.2 Hz, 1H), 3.59 – 3.43 (m, 1H), 3.11 (p, *J* = 6.1, 5.2 Hz, 2H), 2.90 (dq, *J* = 13.5, 7.0 Hz, 1H), 2.20 – 1.53 (m, 14H), 1.12 (d, *J* = 6.6 Hz, 3H) ppm. ^13^C NMR (126 MHz, DMSO) δ 175.0, 161.4, 159.4, 159.0, 158.7, 158.4, 158.1, 156.8, 156.4, 148.0, 141.9, 140.6, 130.2, 129.6, 127.6, 120.8, 120.6, 119.1, 117.4, 115.1, 112.2, 110.2, 108.0, 107.8, 107.4, 68.9, 40.5, 38.0, 35.6, 28.5, 27.626 27.3, 27.2, 26.3, 16.2 ppm. [M + H]^+^ calcd. for C_29_H_35_FN_3_O_2_ 476.2713; found, 476.2724. 3-(2-(2-((2-(2,6-dioxopiperidin-3-yl)-1,3-dioxoisoindolin-4-yl)amino)ethoxy)ethoxy)propanoic acid (35). To a solution of *tert*-butyl 3-(2-(2-aminoethoxy)ethoxy)propanoate (150 mg, 643 µmol, 1.0 equiv) in DMF (3 mL) was added DIPEA (83.1 mg, 112 µL, 643 µmol, 1.0 equiv) followed by the addition of 33 (213 mg, 772 µmol, 1.2 equiv). The mixture was stirred at 75 °C for 12 h. After cooling to 23 °C, the mixture was diluted with DI water (10 mL) and extracted with EtOAc (3 × 10 mL). The combined organic layers were washed with brine (2 × 20 mL), dried via filtration through an isolute phase separator, and concentrated under reduced pressure. The crude product was purified by column chromatography (SiO_2_, CH_2_Cl_2_/acetone, 100:0 to 70:30) to provide *tert*-butyl 3-(2-(2-((2-(2,6-dioxopiperidin-3-yl)-1,3-dioxoisoindolin-4-yl)amino)ethoxy)ethoxy)propanoate (80 mg, 25%). ^1^H NMR (400 MHz, CDCl_3_): δ 8.15 (br s, 1H), 7.50 (dd, *J* = 8.4, 7.2 Hz, 1H), 7.11 (d, *J* = 7.1 Hz, 1H), 6.93 (d, *J* = 8.5 Hz, 1H), 6.50 (br s, 1H), 4.92 (dd, *J* = 12.1, 5.3 Hz, 1H), 4.13 (q, *J* = 7.1 Hz, 1H), 3.75–3.70 (m, 4H), 3.68–3.62 (m, 4H), 3.47 (t, *J* = 5.4 Hz, 2H), 2.89–2.67 (m, 3H), 2.52 (t, *J* = 6.6 Hz, 2H), 2.16–2.09 (m, 1H), 1.45 (s, 9H) ppm. Spectral data matched previously reported characterization data.^57^ To *tert*-butyl 3-(2-(2-((2-(2,6-dioxopiperidin-3-yl)-1,3-dioxoisoindolin-4-yl)amino)ethoxy)ethoxy)propanoate (80 mg, 0.16 mmol, 1.0 equiv) was added a 1:1 solution of TFA/CH_2_Cl_2_ (1 mL) at 0 °C. The reaction was allowed to stir at 23 °C for 1 h. The resulting mixture was concentrated under nitrogen atmosphere and the crude reside was triturated with cold Et_2_O (3 × 5 mL). The resulting solid was dried under vacuum to provide pure 35 (80 mg, 90%) as a green solid. ^1^H NMR (400 MHz, CDCl_3_): δ 8.77 (s, 1H), 7.53–7.44 (m, 1H), 7.09 (d, J = 7.0 Hz, 1H), 6.91 (d, J = 8.5 Hz, 1H), 4.98–4.89 (m, 1H), 3.77 (t, J = 6.3 Hz, 2H), 3.74–3.71 (m, 2H), 3.66 (s, 4H), 3.46 (br t, J = 5.2 Hz, 2H), 2.91–2.70 (m, 3H), 2.64 (t, J = 6.3 Hz, 2H) 2.16–2.08 (m, 1H) ppm. Spectral data matched previously reported characterization data.^57^

### 3-(2-(2-((2-(2,6-dioxopiperidin-3-yl)-1,3-dioxoisoindolin-5-yl)amino)ethoxy)ethoxy)propanoic acid (36)

To a solution of *tert*-butyl 3-(2-(2-aminoethoxy)ethoxy)propanoate (200 mg, 857 µmol, 1.0 equiv) in DMF (2 mL) was added DIPEA (111 mg, 149 µL, 857 µmol, 1.0 equiv) followed by 34 (284 mg, 1.03 mmol, 1.2 equiv). The resulting mixture was allowed to stir at 75 °C for 24 h. After cooling to 23 °C, the mixture was diluted with DI water (5 mL) and extracted with EtOAc (3 × 10 mL). The combined organic layers were washed with brine (2 × 15 mL), dried via filtration through an isolute phase separator and concentrated under reduced pressure. The crude product was purified by column chromatography (SiO_2_, CH_2_Cl_2_/acetone, 100:0 to 70:30) to provide *tert*-butyl 3-(2-(2-((2-(2,6-dioxopiperidin-3-yl)-1,3-dioxoisoindolin-5-yl)amino)ethoxy)ethoxy)propanoate (50 mg, 13%) as a yellow solid. Spectral data matched previously reported characterization data.^58^ To *tert*-butyl 3-(2-(2-((2-(2,6-dioxopiperidin-3-yl)-1,3-dioxoisoindolin-5-yl)amino)ethoxy)ethoxy)propanoate (45 mg, 92 µmol, 1.0 equiv) was added a 1:1 solution of TFA/CH_2_Cl_2_ (1 mL) at 0 °C. The reaction was allowed to stir at 23 °C for 1 h. The resulting mixture was concentrated under nitrogen atmosphere and the crude reside was triturated with Et_2_O (3 × 5 mL). The resulting solid was dried under vacuum to provide 36 (40 mg, 80%) as a green solid, which was taken forward without further purification. Spectral data matched previously reported characterization data.^58^

### (2R)-*N*-(4-((1-(3-(2-(2-((2-(2,6-dioxopiperidin-3-yl)-1,3-dioxoisoindolin-5-yl)amino)ethoxy)ethoxy)propanoyl)piperidin-4-yl)oxy)phenyl)-2-((1s,4S)-4-(6-fluoroquinolin-4-yl)cyclohexyl)propenamide (7)

To a solution of **36** (80 mg, 0.15 mmol, 1.0 equiv) in DMF (2 mL) was added HATU (67 mg, 0.18 mmol, 1.2 equiv) and DIPEA (57 mg, 76 μL, 0.44 mmol, 3.0 equiv) at 0 °C. The resulting mixture was allowed to stir at 23 °C for 30 min at which time (R)-2-((1s,4S)-4-(6-fluoroquinolin-4-yl)cyclohexyl)-N-(4-(piperidin-4-yloxy)phenyl)propanamide (75 mg, 0.15 mmol, 1.0 equiv) was added and the reaction was allowed to continue at 23 °C overnight. The crude product was directly purified by reverse phase HPLC eluting from 10% to 90 % MeCN in water (0.1 % TFA) to provide pure **7** (39 mg, 26%) as an orange solid. ^1^H NMR (500 MHz, CD_3_CN) δ 8.98 (d, *J* = 6.0 Hz, 2H), 8.41 – 8.37 (m, 2H), 8.19 (dd, *J* = 10.1, 2.7 Hz, 1H), 7.99 (d, *J* = 5.8 Hz, 1H), 7.91 (ddd, *J* = 9.4, 7.9, 2.7 Hz, 1H), 7.52 – 7.42 (m, 4H), 6.96 (dd, *J* = 8.0, 1.8 Hz, 1H), 6.84 (ddd, *J* = 17.4, 6.7, 2.6 Hz, 3H), 4.90 (ddd, *J* = 12.3, 5.4, 1.7 Hz, 1H), 4.48 (tt, *J* = 7.5, 3.6 Hz, 1H), 3.74 – 3.55 (m, 10H), 3.35 (dq, *J* = 20.9, 6.2 Hz, 4H), 2.78 – 2.56 (m, 6H), 2.08 – 1.97 (m, 3H), 1.91 – 1.51 (m, 12H), 1.20 (d, *J* = 6.7 Hz, 3H) ppm. ^13^C NMR (126 MHz, CD_3_CN) δ 175.2, 172.6, 170.3, 169.8, 168.4, 168.0, 163.0, 161.0, 160.8, 160.5, 155.0, 154.0, 145.7, 138.7, 135.1, 132.9, 129.2, 129.1, 127.6, 125.5, 123.4, 123.1, 122.0, 119.7, 117.0, 116.4, 115.8, 109.2, 109.0, 106.3, 73.1, 70.6, 70.5, 69.3, 67.6, 49.5, 43.4, 43.0, 41.5, 39.8, 38.9, 36.5, 33.6, 31.6, 31.5, 30.8, 29.0, 28.3, 27.9, 26.9, 22.9, 16.2. [M + H]^+^ calcd. for C_49_H_56_FN_6_O_9_ 891.4093; found, 891.4098.

### (2R)-*N*-(4-chloro-3-((1-(3-(2-(2-((2-(2,6-dioxopiperidin-3-yl)-1,3-dioxoisoindolin-4-yl)amino)ethoxy)ethoxy)propanoyl)piperidin-4-yl)oxy)phenyl)-2-((1s,4*S*)-4-(6-fluoroquinolin-4-yl)cyclohexyl)propenamide (8)

To a solution of **35** (80 mg, 0.15 mmol, 1.0 equiv) in DMF (2 mL) was added HATU (67 mg, 0.18 mmol, 1.2 equiv) and DIPEA (57 mg, 76 μL, 0.44 mmol, 3.0 equiv) at 0 °C. The resulting mixture was allowed to stir at 23 °C for 30 min. **31** (80 mg, 0.15 mmol, 1.0 equiv) was then added and the resulting mixture was allowed to stir at 23 °C for 6 h. The resulting mixture was quenched with DI water (1 mL) and the resulting mixture was directly purified by reverse phase HPLC eluting from 10% to 90 % MeCN in water (0.1 % TFA) to provide pure **8** (31 mg, 22%) was collected as a yellow solid. ^1^H NMR (500 MHz, CD_3_CN) δ 9.08 (s, 1H), 8.99 (h, *J* = 4.0 Hz, 1H), 8.58 (s, 1H), 8.41 (dd, *J* = 9.3, 5.2 Hz, 1H), 8.08 (dd, *J* = 10.4, 2.8 Hz, 1H), 7.90 – 7.83 (m, 1H), 7.80 (ddd, *J* = 9.3, 7.9, 2.7 Hz, 1H), 7.58 (ddt, *J* = 7.4, 5.2, 2.7 Hz, 1H), 7.53 – 7.45 (m, 1H), 7.27 (dd, *J* = 8.6, 1.0 Hz, 1H), 7.10 (tt, *J* = 8.8, 2.1 Hz, 1H), 7.06 – 6.96 (m, 2H), 4.92 (dd, *J* = 12.5, 5.4 Hz, 1H), 4.57 (dt, *J* = 7.0, 3.5 Hz, 1H), 3.74 – 3.34 (m, 15H), 2.80 – 2.62 (m, 4H), 2.55 (tt, *J* = 6.3, 3.1 Hz, 2H), 2.13 – 1.97 (m, 3H), 1.93 – 1.60 (m, 12H), 1.20 (d, *J* = 6.8 Hz, 3H) ppm. ^13^C NMR (126 MHz, CD_3_CN) δ 176.3, 173.0, 170.6, 170.4, 170.2, 168.6, 163.4, 162.8, 161.5, 161.2, 160.9, 153.5, 147.8, 146.0, 139.9, 139.1, 137.0, 133.4, 131.0, 129.6, 129.5, 128.0, 127.9, 123.8, 123.5, 120.2, 116.2, 113.8, 111.7, 110.8, 109.6, 109.4, 108.5, 108.4, 79.2, 78.9, 78.7, 74.5, 71.0, 70.0, 68.2, 68.2, 49.8, 43.1, 42.9, 42.2, 40.2, 38.9, 36.8, 34.0, 32.0, 31.6, 31.0, 29.4, 28.8, 28.2, 27.3, 23.3, 16.6 ppm. [M + H]^+^ calcd. for C_49_H_55_ClFN_6_O_9_ 925.3703; found, 925.3694.

### (2R)-*N*-(4-chloro-3-((1-(3-(2-(2-((2-(2,6-dioxopiperidin-3-yl)-1,3-dioxoisoindolin-5-yl)amino)ethoxy)ethoxy)propanoyl)piperidin-4-yl)oxy)phenyl)-2-((1s,4S)-4-(6-fluoroquinolin-4-yl)cyclohexyl)propenamide (9)

To a solution of **36** (40 mg, 73 μmol, 1.0 equiv) in DMF (2 mL) was added DIPEA (28 mg, 38 μL, 0.22 mmol, 3.0 equiv) and HATU (33 mg, 88 μmol, 1.2 equiv) at 0 °C. The resulting mixture was stirred at 23 °C for 15 min at which time **31** (40 mg, 73 μmol, 1.0 equiv) was added and the reaction mixture was allowed to stir at 23 °C overnight. The crude product was directly purified by reverse phase HPLC eluting from 10% to 90 % MeCN in water (0.1 % FA) to provide pure **9** (24 mg, 34%) as a yellow solid. ^1^H NMR (500 MHz, CD_3_CN) δ 9.03 (t, *J* = 11.4 Hz, 1H), 8.83 (t, *J* = 4.9 Hz, 1H), 8.51 (d, *J* = 5.5 Hz, 1H), 8.14 (dd, *J* = 9.3, 5.6 Hz, 1H), 7.90 (dd, *J* = 10.8, 2.8 Hz, 1H), 7.64 – 7.54 (m, 3H), 7.54 – 7.49 (m, 1H), 7.31 – 7.25 (m, 1H), 7.10 (td, *J* = 8.8, 2.3 Hz, 1H), 7.01 (dd, *J* = 4.5, 2.2 Hz, 1H), 6.86 (ddd, *J* = 7.4, 4.9, 2.1 Hz, 1H), 4.94 – 4.88 (m, 1H), 4.61 – 4.55 (m, 1H), 3.79 – 3.31 (m, 15H), 2.78 – 2.62 (m, 4H), 2.59 – 2.53 (m, 2H), 2.09 – 1.97 (m, 3H), 1.93 – 1.60 (m, 12H), 1.19 (d, *J* = 6.7 Hz, 3H) ppm. ^13^C NMR (126 MHz, CD_3_CN) δ 176.0, 172.6, 170.3, 169.8, 168.5, 168.0, 162.2, 160.2, 155.0, 153.1, 149.3, 144.3, 139.5, 135.1, 132.0, 131.9, 130.6, 128.4, 128.3, 125.5, 120.4, 120.2, 119.2, 116.4, 113.4, 108.1, 108.0, 108.0, 107.9, 106.3, 74.1, 70.6, 70.5, 69.3, 67.6, 49.5, 43.4, 42.6, 41.9, 38.9, 38.5, 36.6, 33.6, 31.6, 31.2, 30.6, 29.2, 28.5, 27.9, 27.9, 27.0, 22.9, 16.1 ppm. [M + H]^+^ calcd. for C_49_H_56_ClFN_6_O_9_ 926.3781; found, 926.3715.

### 2-(2,6-dioxopiperidin-3-yl)-4-(piperazin-1-yl)isoindoline-1,3-dione (37)

To a solution of **33** (200 mg, 724 µmol, 1.0 equiv) and *tert*-butyl piperazine-1-carboxylate (216 mg, 1.16 mmol, 1.6 equiv) in DMSO (2 mL) was added DIPEA (281 mg, 378 µL, 2.17 mmol, 3.0 equiv). The resulting mixture was stirred at 100 °C for 12 h. After cooling to 23 °C the reaction was quenched with DI water and the resulting suspended solid was filtered and washed with cold DI water to afford the intermediate product as a yellow solid. To the intermediate product was added a solution of CH_2_Cl_2_:TFA (1:1, 3 mL) at 0 °C and the resulting reaction mixture was allowed to stir for 1h at 23 °C. The mixture was concentrated under nitrogen atmosphere and the crude residue was triturated with cold Et_2_O (2 × 10 mL) to yield pure **37** (325 mg, 98%) as a yellow solid. Spectral data matches previously reported characterization data.^59^

### 2-(2,6-dioxopiperidin-3-yl)-5-(piperazin-1-yl)isoindoline-1,3-dione (38)

To a solution of **34** (500 mg, 1.81 mmol, 1.0 equiv) and *tert*-butyl piperazine-1-carboxylate (539 mg, 2.90 mmol, 1.6 equiv) in DMSO (5 mL) was added DIPEA (702 mg, 946 μL, 5.43 mmol, 3.0 equiv). The resulting mixture was warmed with stirring to 100 °C and allowed to stir for 12 h. After cooling to 23 °C the reaction was quenched with DI water and the resulting suspended solid was filtered and washed with cold DI water to afford the intermediate product as a yellow solid. To the intermediate product was added a solution of CH_2_Cl_2_:TFA (1:1, 4 mL) at 0 °C and the resulting reaction mixture was allowed to stir for 1h at 23 °C. The mixture was concentrated under nitrogen atmosphere and the crude residue was triturated with cold Et_2_O (2 × 10 mL) to yield pure **38** (700 mg, 85%) as a yellow solid. Spectral data matches previously reported characterization data.^59^

### 3-(2-(4-(2-(2,6-dioxopiperidin-3-yl)-1,3-dioxoisoindolin-4-yl)piperazin-1-yl)ethoxy)propanoic acid (39)

To a solution of **37** (70 mg, 0.20 mmol, 1.0 equiv) and *tert*-butyl 3-(2-iodoethoxy)propanoate (98 mg, 1.6 Eq, 0.33 mmol) in DMF (3 mL) was added DIPEA (79 mg, 0.11 mL, 0.61 mmol, 3.0 equiv). The resulting mixture was allowed to stir at 75 °C for 24 h. The reaction was quenched with DI water (5 mL) and the resulting mixture was extracted with CH_2_Cl_2_ (3 × 5 mL). The organic layers were combined, washed with water (2 × 10 mL), brine (1 × 15 mL), dried via filtration through an isolute phase separator and concentrated *in vacuo*. The crude product was purified by column chromatography (SiO_2_, CH_2_Cl_2_/MeOH, 100:0 to 80:20). The fractions with product were concentrated to yield pure *tert*-butyl 3-(2-(4-(2-(2,6-dioxopiperidin-3-yl)-1,3-dioxoisoindolin-4-yl)piperazin-1-yl)ethoxy)propanoate (100 mg, 95%) as a yellow solid. ^1^H NMR (500 MHz, DMSO) δ 11.11 (s, 1H), 7.69 (dd, *J* = 8.4, 7.2 Hz, 1H), 7.34 (dd, *J* = 15.5, 7.8 Hz, 2H), 5.09 (dd, *J* = 12.8, 5.5 Hz, 1H), 3.58 (t, *J* = 6.1 Hz, 2H), 3.52 (t, *J* = 5.7 Hz, 2H), 3.27 (t, *J* = 4.9 Hz, 3H), 2.87 (ddd, *J* = 16.9, 13.9, 5.5 Hz, 1H), 2.63 – 2.55 (m, 5H), 2.54 – 2.50 (m, 4H), 2.42 (t, *J* = 6.1 Hz, 2H), 2.02 (dtd, *J* = 12.8, 5.1, 2.1 Hz, 1H), 1.39 (s, 9H). ^13^C NMR (126 MHz, DMSO) δ 172.9, 170.6, 170.1, 167.1, 166.3, 149.8, 136.0, 133.7, 123.8, 116.5, 114.9, 79.8, 68.5, 66.2, 57.2, 53.1, 50.5, 48.8, 35.9, 31.0, 27.8, 22.1 ppm. [M + H]^+^ calcd. for C_26_H_35_N_4_O_7_ 515.2506; found, 515.2508. To *tert*-butyl 3-(2-(4-(2-(2,6-dioxopiperidin-3-yl)-1,3-dioxoisoindolin-4-yl)piperazin-1-yl)ethoxy)propanoate (100 mg, 194 μmol, 1.0 equiv) was added a 1:1 solution of TFA:CH_2_Cl_2_ (1 mL) at 0 °C. The resulting mixture was allowed to stir at 23 °C for 1h. The mixture was concentrated under nitrogen atmosphere and the crude residue was triturated with cold Et_2_O (2 × 5 mL). The resulting solid was dried under vacuum to yield **39** (97 mg, 87%) as a yellow solid which was taken forward without additional purification.

### 3-(2-(4-(2-(2,6-dioxopiperidin-3-yl)-1,3-dioxoisoindolin-5-yl)piperazin-1-yl)ethoxy)propanoic acid (40)

To a solution of **38** (118 mg, 345 μmol, 1.0 equiv) and *tert*-butyl 3-(2-iodoethoxy)propanoate (166 mg, 551 μmol, 1.6 equiv) in DMF (3 mL) was added DIPEA (134 mg, 180 μL, 1.03 mmol, 3.0 equiv). The resulting mixture was stirred at 75 °C for 24 h. The mixture was allowed to cool to 23 °C, quenched with cold DI water (5 mL) and extracted with CH_2_Cl_2_ (3 × 10 mL). The organic layers were combined, washed with water (2 × 20 mL), brine (1 × 30 mL), dried via filtration through an isolute phase separator, and concentrated *in vacuo*. To the crude residue was added cold Et_2_O and the resulting mixture was sonicated. Yellow solid precipitated out and was collected through vacuum filtration. The solid was rinsed with cold Et_2_O to provide pure *tert*-butyl 3-(2-(4-(2-(2,6-dioxopiperidin-3-yl)-1,3-dioxoisoindolin-5-yl)piperazin-1-yl)ethoxy)propanoate (150 mg, 85%) as a yellow solid. To *tert*-butyl 3-(2-(4-(2-(2,6-dioxopiperidin-3-yl)-1,3-dioxoisoindolin-5-yl)piperazin-1-yl)ethoxy)propanoate (280 mg, 544 µmol, 1.0 equiv) was added a 1:1 solution of TFA:CH_2_Cl_2_ (2 mL) at 0 °C. The resulting mixture was allowed to stir at 23 °C for 1h. The mixture was concentrated under nitrogen atmosphere and the crude residue was triturated with cold Et_2_O (2 × 5 mL). The resulting solid was dried under vacuum to yield **40** (246 mg, 98%) as a yellow solid which was taken forward without additional purification. ^1^H NMR (500 MHz, DMSO-*d*_6_) δ 12.20 (s, 1H), 10.98 (s, 1H), 7.66 (d, *J* = 8.4 Hz, 1H), 7.37 (s, 1H), 7.24 (dd, *J* = 8.6, 2.3 Hz, 1H), 4.99 (dd, *J* = 12.8, 5.4 Hz, 1H), 4.07 (t, *J* = 4.2 Hz, 2H), 3.65 (t, *J* = 5.0 Hz, 2H), 3.62 – 3.39 (m, 5H), 2.78 (ddd, *J* = 16.6, 13.6, 5.3 Hz, 1H), 2.52 – 2.30 (m, 10H), 1.96 – 1.88 (m, 1H) ppm. ^13^C NMR (126 MHz, DMSO) δ 172.8, 172.7, 170.0, 167.4, 166.9, 154.1, 133.8, 125.0, 119.9, 118.8, 108.9, 66.2, 64.0, 54.7, 50.7, 48.9, 44.2, 34.4, 34.3, 31.0, 28.5, 22.1 ppm. [M + H]^+^ calcd. for C_22_H_27_N_4_O_7_ 459.1880; found, 459.1888.

### 2-(2,6-dioxopiperidin-3-yl)-4-(4-(piperidin-4-ylmethyl)piperazin-1-yl)isoindoline-1,3-dione, Trifluoroacetic acid (41)

To a solution of **37** (200 mg, 439 μmol, 1.0 equiv) and *tert*-butyl 4-(bromomethyl)piperidine-1-carboxylate (147 mg, 527 μmol, 1.2 equiv) in DMF (3 mL) was added DIPEA (170 mg, 230 μL, 1.32 mmol, 3.0 equiv). The resulting mixture was stirred at 75 °C overnight. The reaction mixture was cooled to 23 °C, quenched with cold DI water (5 mL) and extracted with CH_2_Cl_2_ (3 × 10 mL). The organic layers were combined, washed with DI water (2 × 20 mL), brine (1 × 30 mL), dried via filtration through an isolute phase separator, and concentrated *in vacuo*. The crude product was purified was purified by column chromatography (SiO_2_, CH_2_Cl_2_/Acetone, 100:0 to 50:50). The desired fractions were combined and concentrated to yield pure tert-butyl 4-((4-(2-(2,6-dioxopiperidin-3-yl)-1,3-dioxoisoindolin-4-yl)piperazin-1-yl)methyl)piperidine-1-carboxylate (245 mg, 78%) as a bright yellow solid. Spectral data matches previously reported characterization data.^59^ To *tert*-butyl 4-((4-(2-(2,6-dioxopiperidin-3-yl)-1,3-dioxoisoindolin-4-yl)piperazin-1-yl)methyl)piperidine-1-carboxylate (245 mg, 454 μmol, 1.0 equiv) was added a 1:1 mixture of TFA:CH_2_Cl_2_ (2 mL) at 0 °C. The resulting mixture was stirred at 23 °C for 2.5 h. The reaction mixture was concentrated under nitrogen atmosphere and the crude product was triturated with diethyl ether (2 × 10 mL) to yield **41** (235 mg, 93%) as a yellow solid which was taken forward without additional purification.

### 2-(2,6-dioxopiperidin-3-yl)-5-(4-(piperidin-4-ylmethyl)piperazin-1-yl)isoindoline-1,3-dione (42)

To a solution of **38** (200 mg, 584 µmol, 1.0 equiv) and *tert*-butyl 4-(bromomethyl)piperidine-1-carboxylate (195 mg, 701 µmol 1.2 equiv) in DMF (2.5 mL) was added DIPEA (227 mg, 305 µL, 1.75 mmol, 3.0 equiv). The resulting mixture was stirred at 75 °C for 12 h. The reaction was allowed to cool to 23 °C and then quenched with DI water and extracted with CH_2_Cl_2_ (3 × 5 mL). The organic layers were combined and washed with water (2 × 10 mL) and brine (1 × 20 mL). The organic phase was dried using a phase separator and the solvent was removed under vacuum. The crude residue was purified by column chromatography (SiO_2,_ CH_2_Cl_2_/acetone, 100:0 to 0:100) to afford *tert*-butyl 4-((4-(2-(2,6-dioxopiperidin-3-yl)-1,3-dioxoisoindolin-5-yl)piperazin-1-yl)methyl)piperidine-1-carboxylate (300 mg, 556 µmol, 95.2 %) as a yellow solid. To *tert*-butyl 4-((4-(2-(2,6-dioxopiperidin-3-yl)-1,3-dioxoisoindolin-5-yl)piperazin-1-yl)methyl)piperidine-1-carboxylate (300 mg, 556 µmol, 1.0 equiv,) was added a TFA:CH_2_Cl_2_ (1:1, 2.0 mL) at 0 °C. The reaction was stirred at 23 °C for 1 h. The mixture was concentrated to dryness under nitrogen atmosphere. The resulting residue was triturated with Et_2_O (2 × 5 mL) to yield **42** (300 mg, 97%) as a yellow solid. ^1^H NMR (500 MHz, DMSO) δ 11.12 (s, 1H), 7.76 (d, *J* = 8.4 Hz, 1H), 7.49 (s, 1H), 7.36 (d, *J* = 8.5 Hz, 1H), 5.10 (dd, *J* = 12.8, 5.4 Hz, 1H), 4.23 (s, 1H), 3.63 (s, 2H), 3.47 – 3.26 (m, 7H), 3.14 (d, *J* = 22.1 Hz, 3H), 2.88 (ddt, *J* = 20.6, 9.1, 4.3 Hz, 3H), 2.65 – 2.51 (m, 2H), 2.23 – 1.82 (m, 4H), 1.33 (p, *J* = 9.5 Hz, 2H) ppm. Spectral data matches previously reported characterization data.^60^

### 2-(2,6-dioxopiperidin-3-yl)-5-(4-(piperidin-4-yl)piperazin-1-yl)isoindoline-1,3-dione (43)

To a solution of **38** (150 mg, 329 μmol, 1.0 equiv) and *tert*-butyl 4-oxopiperidine-1-carboxylate (78.8 mg, 395 μmol, 1.2 equiv) in DMF (2 mL) was added sodium triacetoxyhydroborate (209 mg, 988 μmol, 3.0 equiv). The reaction mixture was allowed to stir at 23 °C. After 2 h the reaction was quenched with DI water and extracted with EtOAc (3 × 5mL). The organic layers were combined and washed with brine (1 × 15 mL). The organic layer was dried by filtering through phase separator and concentrated to a yellow oil to provide pure tert-butyl 4-(4-(2-(2,6-dioxopiperidin-3-yl)-1,3-dioxoisoindolin-5-yl)piperazin-1-yl)piperidine-1-carboxylate (160 mg, 92%). ^1^H NMR (500 MHz, CDCl_3_) δ 8.10 (s, 1H), 7.69 (d, *J* = 8.5 Hz, 1H), 7.28 (d, *J* = 2.3 Hz, 1H), 7.06 (dd, *J* = 8.6, 2.3 Hz, 1H), 4.94 (dd, *J* = 12.3, 5.4 Hz, 1H), 4.17 (s, 2H), 3.43 (s, 4H), 2.93 – 2.62 (m, 9H), 2.45 (s, 1H), 2.21 – 2.06 (m, 2H), 1.82 (s, 2H), 1.62 (s, 2H), 1.46 (s, 9H) ppm. Spectral data matches previously reported characterization data.^61^ To *tert*-butyl 4-(4-(2-(2,6-dioxopiperidin-3-yl)-1,3-dioxoisoindolin-5-yl)piperazin-1-yl)piperidine-1-carboxylate (250 mg, 476 μmol, 1.0 equiv) was added CH_2_Cl_2_:TFA (1:1, 3.0 mL) at 0 °C. The resulting mixture was allowed to stir at 23 °C. After 1 h the reaction was concentrated to dryness under nitrogen atmosphere. The crude residue was triturated with Et_2_O (2 ×10 mL) and then concentrated to yield **43** (245 mg, 96%) as a dark yellow solid. The product was taken forward without further purification.

### 2-(4-((4-(2-(2,6-dioxopiperidin-3-yl)-1,3-dioxoisoindolin-4-yl)piperazin-1-yl)methyl)piperidin-1-yl)acetic acid (44)

To a solution of **41** (150 mg, 271 μmol, 1.0 equiv) and *tert*-butyl 2-bromoacetate (58.2 mg, 38.6 μL, 299 μmol, 1.1 equiv) in DMF (2 mL) was added DIPEA (42.1 mg, 56.7 μL, 326 μmol, 1.2 equiv). The resulting mixture was stirred at 23 °C for 1 h. The mixture was quenched with DI water (2 mL) and extracted with CH_2_Cl_2_ (3 × 5 mL). The organic layers were combined, washed with water (2 × 10 mL), brine (1 × 15 mL), filtered through isolute phase separator and concentrated to dryness. To the resulting crude residue was added cold Et_2_O. After sonicating for 10 min, a bright yellow solid precipitated out. The Et_2_O was removed with trituration and the resulting solid was concentrated *in vacuo* to provide pure *tert*-butyl 2-(4-((4-(2-(2,6-dioxopiperidin-3-yl)-1,3-dioxoisoindolin-4-yl)piperazin-1-yl)methyl)piperidin-1-yl)acetate (80 mg, 53%) as a yellow solid. ^1^H NMR (500 MHz, CDCl_3_) δ 8.18 (s, 1H), 7.59 (dd, *J* = 8.4, 7.2 Hz, 1H), 7.41 (d, *J* = 7.1 Hz, 1H), 7.21 – 7.14 (m, 1H), 4.95 (dd, *J* = 12.3, 5.4 Hz, 1H), 3.37 (d, *J* = 25.9 Hz, 4H), 3.15 (s, 2H), 2.99 (d, *J* = 11.0 Hz, 2H), 2.95 – 2.53 (m, 6H), 2.47 – 2.16 (m, 4H), 2.15 – 2.07 (m, 1H), 1.79 (d, *J* = 13.0 Hz, 2H), 1.46 (s, 12H), 1.25 (d, *J* = 2.6 Hz, 1H) ppm. ^13^C NMR (126 MHz, CDCl_3_) δ 171.1, 168.3, 167.4, 166.8, 135.9, 134.2, 127.9, 123.6, 116.1, 114.0, 81.3, 64.5, 60.2, 53.8, 53.6, 53.3, 49.2, 32.6, 31.5, 30.8, 29.4, 28.3, 22.8. [M + H]^+^ calcd. for C_29_H_40_N_5_O_6_ 554.2979; found, 554.2990. To *tert*-butyl 2-(4-((4-(2-(2,6-dioxopiperidin-3-yl)-1,3-dioxoisoindolin-4-yl)piperazin-1-yl)methyl)piperidin-1-yl)acetate (50 mg, 90 µmol, 1.0 equiv) was added a 1:1 mixture of CH_2_Cl_2_:TFA (1.5 mL) at 0 °C. The resulting reaction was allowed to stir at 23 °C for 4 h at which time the mixture was concentrated under nitrogen atmosphere. The crude residue was taken up in Et_2_O and triturated (3 × 5 mL). The resulting solid was dried under vacuum to provide **44** (55 mg, 99 %) as a yellow solid which was taken forward without additional purification.

### 2-(4-((4-(2-(2,6-dioxopiperidin-3-yl)-1,3-dioxoisoindolin-5-yl)piperazin-1-yl)methyl)piperidin-1-yl)acetic acid (45)

To a solution of **42** (300 mg, 543 μmol, 1.0 equiv) and *tert*-butyl 2-bromoacetate (127 mg, 84.2 μL, 652 μmol, 1.2 equiv) in DMF (3.0 mL) was added DIPEA (84.2 mg, 113 μL, 652 μmol, 1.2 equiv). The resulting mixture was stirred at 23 °C for 3 h. The reaction mixture was diluted with DI water (5 mL) and extracted with CH_2_Cl_2_ (3 × 10 mL). The organic layers were combined, washed with brine (1 × 20 mL), filtered through a phase separator, and concentrated to dryness. The crude mixture was purified by column chromatography (SiO_2_, CH_2_Cl_2_/acetone, 100:0 to 0:100) to afford *tert*-butyl 2-(4-((4-(2-(2,6-dioxopiperidin-3-yl)-1,3-dioxoisoindolin-5-yl)piperazin-1-yl)methyl)piperidin-1-yl)acetate (250 mg, 83%) as a yellow solid. LCMS (*m/z*): 555.3 [M+2]^+^. ^1^H NMR (500 MHz, CDCl_3_) δ 8.07 (s, 1H), 7.68 (d, *J* = 8.5 Hz, 1H), 7.27 (d, *J* = 2.3 Hz, 1H), 7.04 (dd, *J* = 8.6, 2.4 Hz, 1H), 4.93 (dd, *J* = 12.3, 5.4 Hz, 1H), 3.41 (t, *J* = 5.1 Hz, 4H), 3.15 (s, 2H), 2.98 (d, *J* = 11.0 Hz, 2H), 2.93 – 2.67 (m, 4H), 2.55 (t, *J* = 5.1 Hz, 4H), 2.24 (d, *J* = 7.1 Hz, 3H), 2.21 – 2.07 (m, 2H), 1.76 (d, *J* = 13.0 Hz, 2H), 1.48 (s, 9H), 1.36 (q, *J* = 8.3 Hz, 2H). ppm. ^13^C NMR (126 MHz, CDCl_3_) δ 171.1, 168.4, 168.1, 167.4, 155.7, 134.4, 125.5, 119.5, 117.9, 108.7, 81.2, 64.6, 60.3, 53.4, 53.2, 49.3, 47.6, 33.0, 31.6, 30.8, 28.3, 22.9 ppm. [M + H]^+^ calcd. For C_29_H_40_N_5_O_6_ 554.2979; found, 554.2990. To *tert*-butyl 2-(4-((4-(2-(2,6-dioxopiperidin-3-yl)-1,3-dioxoisoindolin-5-yl)piperazin-1-yl)methyl)piperidin-1-yl)acetate (300 mg, 542 μmol, 1.0 equiv) was added a solution of TFA:CH_2_Cl_2_ (1:1, 2.0 mL) at 0 °C. The resulting mixture was stirred at 23 °C for 2 h. The reaction mixture was concentrated to dryness under nitrogen atmosphere to provide crude product, The crude oil was triturated with Et_2_O (2 × 10 mL) to yield **45** (293 mg, 88%) as a yellow solid. The product was taken forward without further purification.

### 2-(4-(4-(2-(2,6-dioxopiperidin-3-yl)-1,3-dioxoisoindolin-5-yl)piperazin-1-yl)piperidin-1-yl)acetic acid (46)

To a solution of **43** (200 mg, 371 μmol, 1.0 equiv) in DMF (2.5 mL) was added DIPEA (144 mg, 194 μL, 1.11 mmol, 3.0 equiv) followed by *tert*-butyl 2-bromoacetate (86.9 mg, 57.6 μL, 446 μmol, 1.2 equiv). The resulting mixture was stirred for 2 h at 23 °C. The resulting mixture was diluted with water (5 mL) and extracted with CH_2_Cl_2_ (3 x 5mL). The organic layers were combined and washed with brine (1 x 15 mL), dried via filtration through a phase separator, and the filtrate was concentrated to a yellow oil. Cold Et_2_O was added to initiate precipitation of product and the solid was filtered and washed with cold Et_2_O to yield *tert*-butyl 2-(4-(4-(2-(2,6-dioxopiperidin-3-yl)-1,3-dioxoisoindolin-5-yl)piperazin-1-yl)piperidin-1-yl)acetate (190 mg, 95%) as a yellow solid. LCMS (*m/z*): 540.3 [M+1]^+^. ^1^H NMR (500 MHz, CDCl_3_) δ 8.30 (s, 1H), 7.69 (d, *J* = 8.5 Hz, 1H), 7.28 (d, *J* = 2.3 Hz, 1H), 7.05 (dd, *J* = 8.6, 2.4 Hz, 1H), 4.93 (dd, *J* = 12.3, 5.4 Hz, 1H), 3.46 (s, 4H), 3.13 (s, 2H), 3.05 (d, *J* = 10.9 Hz, 2H), 2.94 – 2.67 (m, 7H), 2.22 (t, *J* = 11.4 Hz, 2H), 2.15 – 2.03 (m, 1H), 1.73 (d, *J* = 12.3 Hz, 5H), 1.46 (s, 9H) ppm. ^13^C NMR (126 MHz, CDCl_3_) δ 171.2, 169.8, 168.4, 168.0, 167.4, 155.5, 134.4, 125.5, 118.1, 108.8, 81.3, 61.8, 59.9, 53.9, 52.9, 49.3, 48.5, 47.7, 31.6, 29.4, 28.3, 22.9 ppm. To *tert*-butyl 2-(4-(4-(2-(2,6-dioxopiperidin-3-yl)-1,3-dioxoisoindolin-5-yl)piperazin-1-yl)piperidin-1-yl)acetate (100 mg, 185 µmol, 1.0 equiv) was added CH_2_Cl_2_:TFA (1:1, 2.0 mL) at 0 °C. The resulting mixture was stirred at 23 °C for 2 h. The reaction mixture was concentrated to dryness under nitrogen atmosphere and the crude residue was triturated with Et_2_O (2 × 5 mL) to yield pure **46** (100 mg, 90%) as a yellow solid. The product was taken forward without further purification.

### (2R)-*N*-(4-((1-(2-(4-((4-(2-(2,6-dioxopiperidin-3-yl)-1,3-dioxoisoindolin-4-yl)piperazin-1-yl)methyl)piperidin-1-yl)acetyl)piperidin-4-yl)oxy)phenyl)-2-((1s,4S)-4-(6-fluoroquinolin-4-yl)cyclohexyl)propenamide (10)

To a solution of **44** (25 mg, 41 μmol, 1.0 equiv) in DMF (0.5 mL) was added HATU (19 mg, 49 μmol, 1.2 equiv) and DIPEA (16 mg, 0.12 mmol, 3.0 equiv) at 0 °C under nitrogen atmosphere. The resulting mixture was stirred at 23 °C for 15 min at which time (R)-2-((1s,4S)-4-(6-fluoroquinolin-4-yl)cyclohexyl)-N-(4-(piperidin-4-yloxy)phenyl)propanamide (21 mg, 41 μmol, 1.0 equiv) was added and the reaction was allowed to stir at 23 °C overnight. The reaction mixture was concentrated under nitrogen atmosphere and the crude product was directly purified by reverse phase HPLC eluting from 5% to 75% MeCN in water (0.1 % TFA) to provide pure **10** (9 mg, 20%) was collected as a yellow solid. ^1^H NMR (500 MHz, CD_3_CN) δ 8.98 (d, *J* = 4.1 Hz, 2H), 8.42 (s, 1H), 8.35 (dd, *J* = 9.3, 5.4 Hz, 1H), 8.03 (dd, *J* = 10.6, 2.7 Hz, 1H), 7.80 (d, *J* = 5.2 Hz, 1H), 7.78 – 7.67 (m, 2H), 7.53 – 7.47 (m, 2H), 7.44 (dd, *J* = 7.3, 0.7 Hz, 1H), 7.34 – 7.28 (m, 1H), 6.98 – 6.86 (m, 2H), 4.98 (dd, *J* = 12.8, 5.3 Hz, 1H), 4.55 (s, 1H), 3.99 (d, *J* = 8.9 Hz, 2H), 3.86 – 2.98 (m, 23H), 2.85 – 2.57 (m, 4H), 2.18 – 1.97 (m, 5H), 1.92 – 1.56 (m, 11H), 1.19 (d, *J* = 6.7 Hz, 3H) ppm. ^13^C NMR (126 MHz, CD_3_CN) δ 175.3, 172.5, 170.0, 167.7, 167.4, 162.8, 162.3, 160.8, 153.8, 149.1, 146.8, 140.6, 136.6, 134.5, 133.2, 129.0, 128.9, 124.3, 122.4, 122.2, 122.0, 119.6, 118.8, 117.1, 116.9, 108.8, 108.6, 72.3, 61.1, 57.5, 54.9, 54.0, 52.8, 49.7, 48.1, 41.9, 41.5, 39.6, 39.4, 36.5, 31.6, 30.8, 30.3, 29.1, 29.0, 28.3, 28.0, 27.5, 27.0, 26.5, 22.7, 16.2 ppm. [M + H]^+^ calcd. for C_54_H_64_FN_8_O_7_ 955.4882; found, 955.4885.

### (2R)-*N*-(4-((1-(2-(4-((4-(2-(2,6-dioxopiperidin-3-yl)-1,3-dioxoisoindolin-5-yl)piperazin-1-yl)methyl)piperidin-1-yl)acetyl)piperidin-4-yl)oxy)phenyl)-2-((1s,4S)-4-(6-fluoroquinolin-4-yl)cyclohexyl)propanamide (11)

To a solution of **45** (50 mg, 82 μmol, 1.0 equiv) in DMF (1 mL) was added HATU (37 mg, 98 μmol, 1.2 equiv) and DIPEA (32 mg, 43 μL, 0.25 mmol, 3.0 equiv) at 0 °C under nitrogen atmosphere. The resulting mixture was stirred for 15 min at 23 °C at which time (R)-2-((1s,4S)-4-(6-fluoroquinolin-4-yl)cyclohexyl)-N-(4-(piperidin-4-yloxy)phenyl)propanamide (42 mg, 82 μmol, 1.0 equiv) was added. The reaction was stirred at 23 °C overnight at which time it was dried with nitrogen atmosphere. The crude product was directly purified by reverse phase HPLC eluting from 5% to 50% MeCN in water (0.1 % FA) to provide pure **11** (10 mg, 13%) as a yellow solid. ^1^H NMR (500 MHz, DMSO) δ 11.06 (s, 1H), 10.13 (s, 1H), 8.92 (d, *J* = 4.7 Hz, 1H), 8.12 (dd, *J* = 9.2, 5.8 Hz, 1H), 8.04 (dd, *J* = 10.9, 2.8 Hz, 1H), 7.77 – 7.65 (m, 2H), 7.65 – 7.56 (m, 2H), 7.34 (d, *J* = 8.7 Hz, 1H), 7.15 (dd, *J* = 18.4, 8.7 Hz, 2H), 7.06 (dd, *J* = 7.1, 2.5 Hz, 1H), 6.63 (s, 1H), 5.10 – 5.00 (m, 1H), 4.61 (s, 1H), 3.73 (dt, *J* = 50.7, 5.0 Hz, 8H), 3.57 – 3.39 (m, 7H), 3.35 – 3.23 (m, 3H), 3.12 – 2.80 (m, 5H), 2.64 – 2.51 (m, 2H), 2.05 – 1.56 (m, 19H), 1.13 (d, *J* = 6.6 Hz, 3H) ppm. ^13^C NMR (126 MHz, DMSO) δ 174.4, 172.8, 170.9, 167.4, 166.9, 162.5, 161.0, 159.1, 158.4, 158.2, 157.9, 157.6 154.1, 152.5, 149.3, 144.4, 133.8, 132.8, 132.1, 127.3, 125.0, 120.9, 119.9 119.6, 119.4, 118.8, 117.6, 116.3, 115.3, 112.9, 109.0, 107.4, 71.5, 56.1, 52.6, 50.8, 48.9, 44.1, 41.5, 37.6, 35.6, 31.0, 30.5, 30.1, 28.5, 27.7, 27.4, 26.7, 26.4, 22.1, 16.2 ppm. [M + H]^+^ calcd. for C_54_H_64_FN_8_O_7_ 955.4882; found, 955.4890.

### (2R)-*N*-(4-chloro-3-((1-(2-(4-((4-(2-(2,6-dioxopiperidin-3-yl)-1,3-dioxoisoindolin-4-yl)piperazin-1-yl)methyl)piperidin-1-yl)acetyl)piperidin-4-yl)oxy)phenyl)-2-((1s,4S)-4-(6-fluoroquinolin-4-yl)cyclohexyl)propenamide (12)

To a solution of **44** (55 mg, 90 μmol, 1.0 equiv) in DMF (2 mL) was added HATU (41 mg, 0.11 mmol, 1.2 equiv) and DIPEA (35 mg, 47 μL, 0.27 mmol, 3.0 equiv) at 0 °C under nitrogen atmosphere. The resulting mixture was allowed to stir at 23 °C under nitrogen atmosphere for 15 minutes. **31** (39 mg, 72 μmol, 0.8 equiv) was then added and the reaction was allowed to stir at 23 °C for 48 h. The crude product was directly purified by reverse phase HPLC eluting from 5% to 70% MeCN in water (0.1 % TFA) to provide pure **12** (27 mg, TFA salt, 30%) was collected as a yellow solid. ^1^H NMR (500 MHz, CD_3_CN) δ 9.10 (s, 1H), 9.06 – 8.93 (m, 2H), 8.43 (dd, *J* = 9.3, 5.2 Hz, 1H), 8.10 (dd, *J* = 10.3, 2.7 Hz, 1H), 7.95 (d, *J* = 5.4 Hz, 1H), 7.87 – 7.77 (m, 1H), 7.74 – 7.61 (m, 2H), 7.43 (d, *J* = 7.2 Hz, 1H), 7.28 (dd, *J* = 13.1, 8.5 Hz, 2H), 7.15 (dd, *J* = 8.6, 2.2 Hz, 1H), 4.99 (dd, *J* = 12.8, 5.4 Hz, 1H), 4.65 (s, 1H), 4.02 (d, *J* = 4.6 Hz, 2H), 3.89 – 3.00 (m, 18H), 2.88 – 2.57 (m, 4H), 2.26 (s, 1H), 2.18 – 1.68 (m, 19H), 1.18 (d, *J* = 6.6 Hz, 3H) ppm. ^13^C NMR (126 MHz, CD_3_CN) δ 176.5, 173.0, 170.5, 168.2, 167.9, 163.5, 161.5, 153.4, 149.4, 145.6, 140.1, 138.5, 137.0, 134.9, 131.0, 129.6, 127.4, 124.7, 124.1, 123.8, 120.4, 119.2, 117.3, 114.0, 109.6, 108.6, 73.6, 61.6, 58.0, 54.4, 53.2, 50.1, 48.5, 42.0, 40.3, 39.5, 36.8, 32.0, 30.9, 30.4, 29.5, 28.6, 28.3, 27.9, 27.3, 23.1, 16.6 ppm. [M + H]^+^ calcd. for C_54_H_63_ClFN_8_O_7_ 989.4492; found, 989.4486.

### 2R)-*N*-(4-chloro-3-((1-(2-(4-((4-(2-(2,6-dioxopiperidin-3-yl)-1,3-dioxoisoindolin-5-yl)piperazin-1-yl)methyl)piperidin-1-yl)acetyl)piperidin-4-yl)oxy)phenyl)-2-((1*s*,4*S*)-4-(6-fluoroquinolin-4-yl)cyclohexyl) propenamide (13)

To a solution of **45** (60 mg, 98 μmol, 1.0 equiv) and **31** (54 mg, 98 μmol, 1.0 equiv) in DMF (1 mL) was added DIPEA (42 mg, 56 μL, 0.32 mmol, 3.3 equiv) and HATU (56 mg, 0.15 mmol, 1.5 equiv) at 0 °C under nitrogen atmosphere. The reaction mixture was allowed to stir at 0 °C for 30 min and allowed to warm to 23 °C. After 1h the reaction was complete, and the resulting mixture was concentrated under nitrogen atmosphere and the crude product was directly purified by reverse phase HPLC eluting from 10% to 90 % MeCN in water (0.1 % TFA) to provide pure **13** (63 mg, 53%) as a light green solid. ^1^H NMR (500 MHz, DMSO) δ 11.13 (s, 1H), 10.22 (s, 1H), 8.92 (d, *J* = 4.6 Hz, 1H), 8.13 (dd, *J* = 9.2, 5.8 Hz, 1H), 8.05 (dd, *J* = 10.9, 2.8 Hz, 1H), 7.83 – 7.67 (m, 3H), 7.63 (d, *J* = 4.7 Hz, 1H), 7.56 – 7.47 (m, 1H), 7.37 (dd, *J* = 11.2, 8.7 Hz, 2H), 7.14 (ddd, *J* = 8.8, 5.1, 2.2 Hz, 1H), 5.10 (dd, *J* = 12.8, 5.5 Hz, 1H), 4.67 (s, 1H), 4.29 (dd, *J* = 43.3, 8.9 Hz, 4H), 3.76 – 2.83 (m, 20H), 2.66 – 2.55 (m, 1H), 2.07 – 1.53 (m, 19H), 1.13 (d, *J* = 6.6 Hz, 3H) ppm. ^13^C NMR (126 MHz, DMSO) δ 175.3, 172.9, 170.1, 167.5, 167.0, 162.6, 161.2, 159.2, 158.7, 158.4, 158.1, 157.8, 154.2, 152.1, 149.1, 143.8, 139.4, 133.9, 130.1, 127.4, 125.1, 119.9, 119.8, 118.9, 118.8, 117.6, 116.5, 115.2, 112.9, 109.1, 107.7, 107.5, 107.2, 72.8, 59.9, 56.1, 52.7, 50.8, 48.9, 44.1, 41.2, 38.3, 37.7, 35.6, 31.0, 30.4, 29.8, 28.6, 28.1, 27.6, 27.5, 26.7, 26.3, 22.2, 16.2 ppm. [M + H]^+^ calcd. for C_54_H_63_ClFN_8_O_7_ 989.4492; found, 989.4493.

### (2R)-*N*-(4-((1-(3-(2-(4-(2-(2,6-dioxopiperidin-3-yl)-1,3-dioxoisoindolin-4-yl)piperazin-1-yl)ethoxy)propanoyl)piperidin-4-yl)oxy)phenyl)-2-((1s,4S)-4-(6-fluoroquinolin-4-yl)cyclohexyl)propenamide (14)

To a solution of **39** (40 mg, 70 μmol, 1.0 equiv) in DMF (2 mL) was added HATU (32 mg, 84 μmol, 1.2 equiv) and DIPEA (27 mg, 37 μL, 0.21 mmol, 3.0 equiv) at 0 °C under nitrogen atmosphere. The mixture was stirred for 30 min and then (R)-2-((1s,4S)-4-(6-fluoroquinolin-4-yl)cyclohexyl)-N-(4-(piperidin-4-yloxy)phenyl)propanamide (36 mg, 70 μmol, 1.0 equiv) was added. The mixture was allowed to warm to 23 °C and allowed to stir overnight. The reaction mixture was concentrated under nitrogen atmosphere and the crude product was directly purified by reverse phase HPLC eluting from 10% to 90 % MeCN in water (0.1 % FA) to provide pure **14** (28 mg, 42%) as a yellow solid. ^1^H NMR (500 MHz, CD3CN) δ 8.83 (s, 1H), 8.66 (d, *J* = 4.6 Hz, 1H), 8.18 (s, 1H), 7.95 – 7.91 (m, 1H), 7.69 (dd, *J* = 11.0, 2.8 Hz, 1H), 7.49 (dd, *J* = 8.4, 7.2 Hz, 1H), 7.38 (ddd, *J* = 9.3, 8.2, 2.8 Hz, 1H), 7.34 (d, *J* = 4.6 Hz, 1H), 7.33 – 7.29 (m, 2H), 7.19 (d, *J* = 7.2 Hz, 1H), 7.11 (d, *J* = 8.4 Hz, 1H), 6.76 – 6.68 (m, 2H), 4.81 (dd, *J* = 12.5, 5.4 Hz, 1H), 4.34 (dq, *J* = 7.6, 3.8 Hz, 1H), 3.71 (ddd, *J* = 12.2, 6.8, 3.8 Hz, 1H), 3.50 (dt, *J* = 24.5, 5.8 Hz, 5H), 3.21 (dt, *J* = 14.4, 5.1 Hz, 6H), 2.70 (t, *J* = 4.9 Hz, 3H), 2.67 – 2.39 (m, 9H), 1.92 (dtd, *J* = 12.8, 5.1, 2.5 Hz, 1H), 1.84 (dt, *J* = 14.7, 3.3 Hz, 2H), 1.75 – 1.47 (m, 9H), 1.42 (dp, *J* = 12.7, 3.9 Hz, 1H), 1.02 (d, *J* = 6.7 Hz, 3H) ppm. ^13^C NMR (126 MHz, CD_3_CN) δ 175.4, 172.5, 170.2, 170.0, 167.9, 167.3, 163.0, 161.9, 160.0, 153.9, 153.3, 150.4, 150.3, 146.2, 136.3, 134.6, 133.4, 133.3, 133.0, 128.2, 128.1, 124.2, 121.9, 119.4, 119.2, 119.0, 117.0, 115.9, 107.7, 107.5, 72.9, 67.1, 66.6, 57.0, 53.3, 50.5, 49.7, 42.9, 41.5, 39.0, 38.7, 36.7, 33.3, 31.6, 31.4, 30.8, 29.3, 28.5, 28.1, 27.2, 22.8, 16.2 ppm. [M + H]^+^ calcd. for C_51_H_59_FN_7_O_8_ 916.4409; found, 917.4488.

### (2R)-*N*-(4-((1-(3-(2-(4-(2-(2,6-dioxopiperidin-3-yl)-1,3-dioxoisoindolin-5-yl)piperazin-1-yl)ethoxy)propanoyl)piperidin-4-yl)oxy)phenyl)-2-((1s,4S)-4-(6-fluoroquinolin-4-yl)cyclohexyl)propenamide (15)

To a solution of **40** (100 mg, 175 μmol, 1.0 equiv) in DMF (3 mL) was added HATU (79.7 mg, 210 μmol, 1.2 equiv) and DIPEA (67.7 mg, 91.3 μL, 524 μmol, 3.0 equiv) at 0 °C under nitrogen atmosphere. The resulting mixture was allowed to stir at 23 °C for 30 min at which time (R)-2-((1s,4S)-4-(6-fluoroquinolin-4-yl)cyclohexyl)-N-(4-(piperidin-4-yloxy)phenyl)propanamide (89.4 mg, 175 μmol, 1.0 equiv) was added and the resulting mixture was stirred overnight. The reaction mixture was concentrated under nitrogen atmosphere and the crude product was directly purified by reverse phase HPLC eluting from 10% to 90 % MeCN in water (0.1 % FA) to provide pure **15** (63 mg, 37%) as a yellow solid. ^1^H NMR (500 MHz, CD_3_CN) δ 9.01 (d, *J* = 15.2 Hz, 1H), 8.82 (d, *J* = 4.5 Hz, 1H), 8.35 (s, 1H), 8.17 – 8.00 (m, 2H), 7.85 (dd, *J* = 11.0, 2.8 Hz, 1H), 7.67 (d, *J* = 8.5 Hz, 1H), 7.54 (ddd, *J* = 9.2, 8.1, 2.8 Hz, 1H), 7.51 – 7.44 (m, 3H), 7.32 (d, *J* = 2.3 Hz, 1H), 7.18 (dd, *J* = 8.6, 2.4 Hz, 1H), 6.97 – 6.77 (m, 2H), 4.94 (ddd, *J* = 12.4, 5.3, 1.5 Hz, 1H), 4.50 (tt, *J* = 7.4, 3.6 Hz, 1H), 3.85 (ddd, *J* = 12.3, 7.2, 3.9 Hz, 1H), 3.68 (dt, *J* = 11.3, 5.7 Hz, 5H), 3.53 (t, *J* = 5.1 Hz, 4H), 3.36 (ddd, *J* = 13.7, 8.4, 3.6 Hz, 4H), 2.88 (dt, *J* = 11.1, 5.3 Hz, 6H), 2.83 – 2.58 (m, 7H), 2.12 – 1.98 (m, 2H), 1.92 – 1.49 (m, 9H), 1.17 (d, *J* = 6.7 Hz, 3H) ppm. ^13^C NMR (126 MHz, CD_3_CN) δ 175.4, 172.6, 172.6, 170.4, 170.2, 170.1, 168.3, 167.7, 162.8, 161.9, 160.0, 155.7, 153.9, 153.4, 153.3, 150.4, 150.4, 146.1, 134.8, 133.3, 133.1, 128.2, 128.1, 125.4, 121.9, 120.4, 119.5, 119.2, 119.1, 118.9, 117.0, 109.0, 107.7, 107.5, 72.9, 66.2, 66.1, 56.5, 52.6, 49.6, 46.8, 42.9, 41.5, 39.1, 38.7, 36.7, 33.2, 31.6, 31.4, 30.7, 29.3, 28.5, 28.1, 27.2, 22.8, 16.2 ppm. [M + H]^+^ calcd. for C_51_H_59_FN_7_O_8_ 916.4409; found, 916.4414.

### 2R)-*N*-(4-chloro-3-((1-(3-(2-(4-(2-(2,6-dioxopiperidin-3-yl)-1,3-dioxoisoindolin-4-yl)piperazin-1-yl)ethoxy)propanoyl)piperidin-4-yl)oxy)phenyl)-2-((1s,4S)-4-(6-fluoroquinolin-4-yl)cyclohexyl)propenamide (16)

To a solution of **39** (40 mg, 70 μmol, 1.0 equiv) in DMF (2 mL) was added HATU (32 mg, 84 μmol, 1.2 equiv) and DIPEA (27 mg, 37 μL, 0.21 mmol, 3.0 equiv) at 0 °C under nitrogen atmosphere. The resulting mixture was allowed to stir for 30 min at 23 °C. **31** (38 mg, 70 μmol, 1.0 equiv) was added and the reaction was stirred at 23 °C overnight. The reaction mixture was concentrated under nitrogen atmosphere and the crude product was directly purified by reverse phase HPLC eluting from 10% to 90 % MeCN in water (0.1 % FA) to provide pure **16** (15 mg, 22%) as a yellow solid. ^1^H NMR (500 MHz, CD_3_CN) δ 9.02 (s, 1H), 8.82 (d, *J* = 4.5 Hz, 1H), 8.53 (s, 1H), 8.09 (dd, *J* = 9.2, 5.8 Hz, 1H), 7.84 (dd, *J* = 11.0, 2.8 Hz, 1H), 7.61 (ddt, *J* = 13.8, 8.0, 2.4 Hz, 2H), 7.57 – 7.47 (m, 2H), 7.32 (dd, *J* = 7.2, 3.8 Hz, 1H), 7.26 (t, *J* = 8.1 Hz, 2H), 7.10 (tt, *J* = 7.1, 2.2 Hz, 1H), 4.96 (dd, *J* = 12.5, 5.4 Hz, 1H), 4.66 – 4.56 (m, 1H), 3.78 – 3.63 (m, 4H), 3.62 – 3.21 (m, 9H), 2.81 – 2.51 (m, 11H), 2.11 – 1.95 (m, 5H), 1.91 – 1.64 (m, 10H), 1.18 (d, *J* = 6.7 Hz, 3H) ppm. ^13^C NMR (126 MHz, CD_3_CN) δ 176.0, 172.6, 170.1, 169.9, 167.9, 167.3, 161.9, 160.0, 153.2, 153.1, 150.7, 150.4, 146.2, 139.6, 136.2, 134.7, 133.4, 133.303 130.6, 128.2, 128.1, 124.2, 119.5, 119.2, 119.0, 115.5, 113.3, 108.0, 107.7, 107.5, 74.1, 68.7, 67.5, 57.9, 53.7, 51.2, 49.7, 42.7, 41.8, 38.7, 38.5, 36.6, 33.6, 31.6, 31.3, 30.6, 29.3, 28.5, 28.0, 27.1, 22.7, 16.2 ppm. [M + H]^+^ calcd. for C_51_H_58_ClFN_7_O_8_ 950.4019; found, 950.4019.

### (2R)-*N*-(4-chloro-3-((1-(3-(2-(4-(2-(2,6-dioxopiperidin-3-yl)-1,3-dioxoisoindolin-5-yl)piperazin-1-yl)ethoxy)propanoyl)piperidin-4-yl)oxy)phenyl)-2-((1s,4S)-4-(6-fluoroquinolin-4-yl)cyclohexyl)propenamide (17)

To a solution of **40** (100 mg, 175 μmol, 1.0 equiv) in DMF (3 mL) was added HATU (79.7 mg, 210 μmol, 1.2 equiv) and DIPEA (67.7 mg, 91.3 μL, 524 μmol, 3.0 equiv) at 0 °C under nitrogen atmosphere. The resulting mixture was allowed to stir at 23 °C for 30 min. After 30 min **31** (95.5 mg, 175 μmol, 1.0 equiv) was added and the reaction was stirred at 23 °C overnight. The reaction mixture was concentrated under nitrogen atmosphere and the crude product was directly purified by reverse phase HPLC eluting from 10% to 90 % MeCN in water (0.1 % FA) to provide pure **17** (57 mg, 33%) as a yellow solid. ^1^H NMR (500 MHz, CD_3_CN) δ 9.04 – 8.91 (m, 1H), 8.81 (d, *J* = 4.5 Hz, 1H), 8.52 (s, 1H), 8.12 – 8.04 (m, 2H), 7.84 (dd, *J* = 11.0, 2.9 Hz, 1H), 7.68 – 7.58 (m, 2H), 7.58 – 7.47 (m, 2H), 7.31 – 7.22 (m, 2H), 7.16 (dd, *J* = 8.6, 2.4 Hz, 1H), 7.08 (dt, *J* = 8.7, 2.2 Hz, 1H), 4.93 (dd, *J* = 12.4, 5.4 Hz, 1H), 4.61 (dt, *J* = 7.1, 3.5 Hz, 1H), 3.70 (q, *J* = 6.8 Hz, 4H), 3.63 (td, *J* = 5.6, 1.6 Hz, 2H), 3.58 – 3.30 (m, 7H), 2.83 – 2.64 (m, 13H), 2.11 – 1.98 (m, 3H), 1.89 – 1.68 (m, 9H), 1.18 (d, *J* = 6.7 Hz, 3H) ppm. ^13^C NMR (126 MHz, DMSO) δ 175.1, 172.8, 170.1, 168.8, 167.5, 167.0, 163.0, 160.9, 159.0, 152.4, 152.2, 149.7, 145.2, 139.3, 133.8, 132.7, 129.9, 127.2, 124.9, 119.2, 119.0, 118.6, 116.5, 112.7, 107.3, 107.1, 73.3, 66.7, 48.8, 41.8, 37.8, 37.4, 35.6, 32.7, 31.0, 30.7, 30.0, 28.5, 27.6, 27.5, 26.3, 22.2, 16.1. [M + H]^+^ calcd. for C_51_H_58_ClFN_7_O_8_ 950.4019; found, 950.4017.

### (2R)-*N*-(4-((1-(2-(4-(4-(2-(2,6-dioxopiperidin-3-yl)-1,3-dioxoisoindolin-5-yl)piperazin-1-yl)piperidin-1-yl)acetyl)piperidin-4-yl)oxy)phenyl)-2-((1s,4S)-4-(6-fluoroquinolin-4-yl)cyclohexyl)propenamide (18)

To a solution of **46** (44 mg, 74 µmol, 1.0 equiv) in DMF (2 mL) was added HATU (34 mg, 88 µmol, 1.2 equiv) and DIPEA (29 mg, 38 µL, 0.22 mmol, 3.0 equiv) at 0 °C under nitrogen atmosphere. The resulting mixture was allowed to stir at 23 °C under nitrogen atmosphere for 15 minutes. (R)-2-((1s,4S)-4-(6-fluoroquinolin-4-yl)cyclohexyl)-N-(4-(piperidin-4-yloxy)phenyl)propanamide (30 mg, 59 µmol, 0.8 equiv) was then added and the reaction was stirred at 23 °C for 4 h. The resulting mixture was concentrated under nitrogen atmosphere and the crude product was directly purified by reverse phase HPLC eluting from 5% to 70% MeCN in water (0.1 % TFA) to provide pure **18** (14 mg, 18%) was collected as a yellow solid. ^1^H NMR (500 MHz, CD_3_CN) δ 9.00 (d, *J* = 6.2 Hz, 2H), 8.47 (s, 1H), 8.40 (dd, *J* = 9.3, 5.3 Hz, 1H), 8.07 (dd, *J* = 10.5, 2.7 Hz, 1H), 7.86 (d, *J* = 5.3 Hz, 1H), 7.84 – 7.75 (m, 1H), 7.71 (d, *J* = 8.4 Hz, 1H), 7.53 – 7.46 (m, 2H), 7.37 (d, *J* = 2.4 Hz, 1H), 7.23 (dd, *J* = 8.5, 2.4 Hz, 1H), 6.96 – 6.84 (m, 2H), 4.95 (dd, *J* = 12.5, 5.4 Hz, 1H), 4.55 (tt, *J* = 7.2, 3.5 Hz, 1H), 4.07 (s, 2H), 3.85 – 3.13 (m, 17H), 2.85 – 2.61 (m, 5H), 2.40 – 2.28 (m, 4H), 2.15 – 1.97 (m, 4H), 1.91 – 1.62 (m, 10H), 1.19 (d, *J* = 6.6 Hz, 3H) ppm. ^13^C NMR (126 MHz, CD_3_CN) δ 175.8, 173.0, 170.5, 168.5, 168.1, 163.4, 161.4, 155.2, 154.2, 146.4, 139.6, 135.2, 133.6, 129.5, 128.3, 125.9, 123.4, 122.4, 121.9, 120.2, 119.9, 117.5, 110.0, 109.4, 72.7, 68.3, 60.2, 50.1, 49.2, 45.7, 42.3, 41.9, 40.2, 39.8, 36.9, 32.0, 31.2, 30.8, 29.5, 28.7, 28.4, 27.4, 26.2, 23.2, 16.6 ppm. [M + H]^+^ calcd. for C_53_H_62_FN_8_O_7_ 941.4725; found, 941.4723.

### (2R)-*N*-(4-chloro-3-((1-(2-(4-(4-(2-(2,6-dioxopiperidin-3-yl)-1,3-dioxoisoindolin-5-yl)piperazin-1-yl)piperidin-1-yl)acetyl)piperidin-4-yl)oxy)phenyl)-2-((1*s*,4*S*)-4-(6-fluoroquinolin-4-yl)cyclohexyl)propenamide (19)

To a solution of the **43** (150 mg, 251 μmol, 1.0 equiv) in DMF (2.5 mL) was added DIPEA (162 mg, 219 μL, 1.26 mmol, 5.0 equiv) and **31** (137 mg, 251 μmol, 1.0 equiv). The reaction mixture was cooled to 0 °C and 2,4,6-tripropyl-1,3,5,2,4,6-trioxatriphosphinane 2,4,6-trioxide (480 mg, 449 μL, 50% Wt in EtOAc, 754 μmol, 3.0 equiv) was added slowly. The reaction mixture was allowed to warm to 23 °C and stirred for 1 h. The reaction was quenched with cold DI water (2 mL), extracted with EtOAc (3 x 5 mL), and dried via filtration through a phase separator. The filtrate was concentrated under reduced pressure the crude product was directly purified by reverse phase HPLC eluting from 10% to 90 % MeCN in water (0.1 % TFA) to provide pure **19** (150 mg, 49%) as a yellow solid. ^1^H NMR (500 MHz, DMSO) δ 11.13 (s, 1H), 10.22 (d, *J* = 2.0 Hz, 1H), 8.92 (d, *J* = 4.7 Hz, 1H), 8.13 (dd, *J* = 9.3, 5.7 Hz, 1H), 8.05 (dd, *J* = 10.9, 2.8 Hz, 1H), 7.84 – 7.70 (m, 3H), 7.64 (d, *J* = 4.8 Hz, 1H), 7.51 (d, *J* = 2.3 Hz, 1H), 7.37 (dd, *J* = 11.9, 8.7 Hz, 2H), 7.15 (ddd, *J* = 8.0, 5.4, 2.1 Hz, 1H), 5.10 (dd, *J* = 12.8, 5.4 Hz, 1H), 4.68 (s, 1H), 4.35 (s, 4H), 3.75 – 3.10 (m, 15H), 3.02 (s, 2H), 2.97 – 2.82 (m, 2H), 2.70 – 2.53 (m, 1H), 2.32 (d, *J* = 12.5 Hz, 2H), 2.08 – 1.56 (m, 16H), 1.14 (s, 3H) ppm. ^13^C NMR (126 MHz, DMSO) δ 175.3, 172.9, 170.1, 167.5, 167.0, 162.6, 161.2, 159.2, 158.7, 158.4, 158.1, 157.9, 154.2, 152.1, 149.1, 143.9, 139.4, 133.8, 131.8, 130.1, 127.4, 125.1, 120.0, 119.7, 118.9, 118.8, 117.6, 116.5, 115.2, 112.9, 109.0, 107.7, 107.5, 107.2, 72.7, 59.2, 55.9, 51.6, 48.9, 48.0, 44.5, 41.2, 38.4, 37.7, 35.6, 31.0, 30.3, 29.8, 28.6, 27.6, 27.5, 26.3, 23.7, 22.2, 16.2 ppm. [M + H]^+^ calcd. for C_53_H_61_ClFN_8_O_7_ 975.4336; found, 975.4339.

### (2R)-N-(3-((1-(2-(4-((4-(2-(2,6-dioxopiperidin-3-yl)-1,3-dioxoisoindolin-5-yl)piperazin-1-yl)methyl)piperidin-1-yl)acetyl)piperidin-4-yl)oxy)phenyl)-2-((1s,4S)-4-(6-fluoroquinolin-4-yl)cyclohexyl)propenamide (20)

To a solution of **45** (50 mg, 82 μmol, 1.0 equiv) in DMF (1 mL) was added DIPEA (32 mg, 43 μL, 0.25 mmol, 3.0 equiv) and HATU (31 mg, 82 μmol, 1.0 equiv) at 0 °C under nitrogen atmosphere. The resulting reaction mixture was stirred at 23 °C for 30 min. **32** (42 mg, 82 μmol, 1.0 equiv) was then added and the reaction was stirred at 23 °C for 1 h. The resulting mixture was concentrated under nitrogen atmosphere and the crude product was directly purified by reverse phase HPLC eluting from 10% to 90 % MeCN in water (0.1 % TFA) to provide pure **20** (32 mg, 2 x TFA salt, 33%) as a yellow solid. ^1^H NMR (500 MHz, DMSO) δ 11.13 (s, 1H), 10.03 (s, 1H), 8.90 (d, *J* = 4.5 Hz, 1H), 8.11 (dd, *J* = 9.2, 5.8 Hz, 1H), 8.02 (dd, *J* = 11.0, 2.8 Hz, 1H), 7.78 (d, *J* = 8.4 Hz, 1H), 7.70 (td, *J* = 8.6, 2.8 Hz, 1H), 7.60 (d, *J* = 4.7 Hz, 1H), 7.55 – 7.42 (m, 2H), 7.38 (d, *J* = 8.7 Hz, 1H), 7.20 (td, *J* = 8.1, 2.5 Hz, 1H), 7.07 (d, *J* = 7.9 Hz, 1H), 6.69 (dd, *J* = 8.2, 2.5 Hz, 1H), 5.10 (dd, *J* = 12.8, 5.5 Hz, 1H), 4.62 (s, 1H), 4.34 – 4.21 (m, 3H), 3.90 – 2.79 (m, 22H), 2.69 – 2.51 (m, 2H), 2.17 – 1.44 (m, 17H), 1.12 (d, *J* = 6.6 Hz, 3H) ppm. ^13^C NMR (126 MHz, DMSO) δ 175.1, 172.9, 170.1, 167.5, 167.0, 162.6, 161.1, 159.1, 158.4, 158.2, 157.9, 157.7, 157.1, 149.5, 144.7, 140.6, 133.9, 132.3, 129.7, 127.4, 127.3, 125.1, 119.6, 119.3, 118.8, 118.0, 115.6, 111.9, 110.2, 109.1, 107.5, 107.3, 71.1, 56.1, 52.7, 50.8, 48.9, 44.1, 41.5, 37.6, 35.6, 31.0, 30.6, 30.1, 28.6, 27.7, 27.4, 26.7, 26.4, 22.2, 16.2 ppm. [M + H]^+^ calcd. for C_54_H_64_FN_8_O_7_ 955.4882; found, 955.4873.

### (2R)-*N*-(3-((1-(2-(4-(4-(2-(2,6-dioxopiperidin-3-yl)-1,3-dioxoisoindolin-5-yl)piperazin-1-yl)piperidin-1-yl)acetyl)piperidin-4-yl)oxy)phenyl)-2-((1s,4S)-4-(6-fluoroquinolin-4-yl)cyclohexyl)propenamide (21)

To a solution of **46** (125 mg, 210 μmol, 1.0 equiv) in DMF (3 mL) was added DIPEA (81.2 mg, 109 μL, 629 μmol, 3.0 equiv) and HATU (79.7 mg, 210 μmol, 1.0 equiv) at 0 °C under nitrogen atmosphere. The resulting reaction mixture was stirred at 23 °C for 30 min at which time **32** (85.8 mg, 168 μmol, 0.8 equiv) was added. The reaction mixture was stirred at 23 °C for 1 h and then concentrated to dryness under nitrogen atmosphere. The crude product was directly purified by reverse phase HPLC eluting from 10% to 90 % MeCN in water (0.1 % TFA) to provide pure **21** (87 mg, 36%) as a yellow solid. ^1^H NMR (500 MHz, DMSO) δ 11.13 (s, 1H), 10.01 (s, 1H), 8.89 (d, *J* = 4.6 Hz, 1H), 8.10 (dd, *J* = 9.2, 5.8 Hz, 1H), 8.01 (dd, *J* = 10.9, 2.8 Hz, 1H), 7.78 (d, *J* = 8.3 Hz, 1H), 7.69 (td, *J* = 8.7, 2.8 Hz, 1H), 7.59 (d, *J* = 4.6 Hz, 1H), 7.56 – 7.31 (m, 3H), 7.20 (t, *J* = 8.1 Hz, 1H), 7.06 (d, *J* = 8.0 Hz, 1H), 6.69 (dd, *J* = 8.4, 2.4 Hz, 1H), 5.10 (dd, *J* = 12.8, 5.5 Hz, 1H), 4.62 (s, 1H), 4.33 (s, 2H), 3.94 – 2.82 (m, 25H), 2.67 – 2.54 (m, 2H), 2.42 – 2.19 (m, 2H), 2.17 – 1.51 (m, 13H), 1.12 (d, *J* = 6.5 Hz, 3H) ppm. ^13^C NMR (126 MHz, DMSO) δ 175.0, 172.8, 170.0, 167.4, 166.9, 162.5, 161.1, 159.2, 158.6, 158.3, 158.1, 157.8, 157.1, 154.2, 153.9, 149.2, 144.1, 140.6, 133.8, 131.9, 129.6, 127.4, 127.3, 125.0, 120.0, 119.8, 119.6, 118.8, 117.6, 115.3, 112.0, 110.3, 109.0, 107.6, 107.4, 71.2, 59.2, 55.9, 51.6, 48.9, 48.0, 44.6, 41.5, 40.2, 38.7, 37.7, 35.6, 31.0, 30.5, 30.0, 28.5, 27.6, 27.4, 26.4, 23.7, 22.2, 16.2 ppm. [M + H]^+^ calcd. for C_53_H_62_FN_8_O_7_ 941.4725; found, 941.4713.

### 3-(4-(2-(2,6-dioxopiperidin-3-yl)-1,3-dioxoisoindolin-5-yl)piperazin-1-yl)propanoic acid (47)

To a solution of **38** (100 mg, 292 μmol, 1.0 equiv) in DMF (1.5 mL) was added DIPEA (113 mg, 876 μmol, 3.0 equiv) and *tert*-butyl 3-bromopropanoate (73.3 mg, 58.6 μL, 351 μmol, 1.2 equiv). The resulting reaction mixture was allowed to stir at 23 °C for 12 h. The reaction mixture was quenched with DI water (3 mL) and extracted with CH_2_Cl_2_ (3 x 5mL). The organic layers were combined, washed with brine (1 x), filtered through a phase separator and concentrated *in vacuo* to provide an oil residue. The resulting residue was triturated with Et_2_O (2 × 5 mL) and concentrated *in vacuo* to provide pure intermediate as a yellow solid. The intermediate was then taken up in a 1:1 mixture of TFA:CH_2_Cl_2_ (1 mL) at 0 °C and stirred at 23 °C for 24 h. The resulting reaction mixture was concentrated under nitrogen atmosphere. The resulting residue was triturated with Et_2_O (2 × 10 mL) and the resulting yellow solid was dried *in vacuo* to provide **47** (70 mg, 45%) as a yellow solid. Spectral data matches previously reported characterization data.^62^

### 2-(4-(2-(2,6-dioxopiperidin-3-yl)-1,3-dioxoisoindolin-5-yl)piperazin-1-yl)acetic acid (48)

To a solution of **38** (100 mg, 292 μmol, 1.0 equiv) in DMF (1.5 mL) was added DIPEA (113 mg, 876 μmol, 3.0 equiv) and *tert*-butyl 2-bromoacetate (68.4 mg, 351 μmol, 1.2 equiv). The resulting reaction mixture was stirred at 23 °C for 2 h at which time the reaction mixture was quenched with DI water and extracted with CH_2_Cl_2_ (3 × 5 mL). The organic layers were combined, washed with brine (1 × 15 mL), filtered through a phase separator and concentrated *in vacuo* to provide an oil residue. The resulting residue was triturated with Et_2_O (2 × 5mL) and concentrated *in vacuo* to provide pure intermediate as a yellow solid. The intermediate was then taken up in a 1:1 mixture of TFA:CH_2_Cl_2_ (1 mL) at 0 °C and stirred at 23 °C for 24 h. The resulting reaction mixture was concentrated under nitrogen atmosphere. The resulting residue was triturated with Et_2_O (2 × 10 mL) and the resulting yellow solid was dried *in vacuo* to provide **48** (100 mg, 67%) as a yellow solid. Spectral data matches previously reported characterization data.^62^

### 2R)-*N*-(3-((1-(2-(4-(2-(2,6-dioxopiperidin-3-yl)-1,3-dioxoisoindolin-5-yl)piperazin-1-yl)acetyl)piperidin-4-yl)oxy)phenyl)-2-((1s,4S)-4-(6-fluoroquinolin-4-yl)cyclohexyl)propenamide (22)

To a solution of **47** (20 mg, 39 μmol, 1.0 equiv) in DMF (1 mL) was added DIPEA (15 mg, 20 μL, 0.12 mmol, 3.0 equiv) and HATU (18 mg, 147 μmol, 1.2 equiv) at 0 °C under nitrogen atmosphere3. The resulting mixture was stirred at 23 °C for 15 min at which time **32** (16 mg, 31 μmol, 0.8 equiv) was added and the resulting mixture was stirred at 23 °C. After 30 min the reaction was complete, and the resulting mixture was concentrated under nitrogen atmosphere. The crude product was directly purified by reverse phase HPLC eluting from 10% to 90 % MeCN in water (0.1 % TFA) to provide pure **22** (10 mg, 26%) as a yellow solid. ^1^H NMR (500 MHz, DMSO) δ 11.09 (s, 1H), 9.97 (s, 1H), 8.89 (d, *J* = 4.6 Hz, 1H), 8.11 (dd, *J* = 9.2, 5.8 Hz, 1H), 8.03 – 7.97 (m, 1H), 7.78 (d, *J* = 8.5 Hz, 1H), 7.73 – 7.66 (m, 1H), 7.59 (d, *J* = 4.7 Hz, 1H), 7.48 (s, 2H), 7.38 – 7.31 (m, 1H), 7.22 – 7.18 (m, 1H), 7.08 (d, *J* = 7.8 Hz, 1H), 6.72 – 6.66 (m, 1H), 5.10 (dd, *J* = 12.8, 5.5 Hz, 1H), 4.61 (s, 1H), 4.43 – 3.17 (m, 7H, hidden under water peak), 2.88 (q, *J* = 11.7 Hz, 3H), 2.65 – 2.53 (m, 3H), 2.36 (s, 1H), 2.06 – 1.55 (m, 19H), 1.13 (d, *J* = 6.6 Hz, 3H) ppm. [M + H]^+^ calcd. for C_48_H_53_FN_7_O_7_ 858.3991; found, 858.3986.

### (2R)-*N*-(3-((1-(3-(4-(2-(2,6-dioxopiperidin-3-yl)-1,3-dioxoisoindolin-5-yl)piperazin-1-yl)propanoyl)piperidin-4-yl)oxy)phenyl)-2-((1s,4S)-4-(6-fluoroquinolin-4-yl)cyclohexyl)propenamide (23)

To a solution of **48** (20 mg, 48 μmol, 1.0 equiv) in DMF (1 mL) was added DIPEA (19 mg, 25 μL, 0.14 mmol, 3.0 equiv) and HATU (15 mg, 39 μmol, 0.8 equiv) at 0 °C under nitrogen atmosphere. The resulting reaction mixture was allowed to stir at 23 °C for 15 min at which time **32** (20 mg, 39 μmol, 0.8 equiv) was added at 0 °C. The resulting mixture was stirred at 23 °C for 30 min. The mixture was quenched with DI water (1 mL) and concentrated under nitrogen atmosphere. The resulting crude product was directly purified by reverse phase HPLC eluting from 10% to 90 % MeCN in water (0.1 % TFA) to provide pure **23** (10 mg, 21%) as a yellow solid. ^1^H NMR (500 MHz, DMSO) δ 11.09 (s, 1H), 9.96 (s, 1H), 8.89 (d, *J* = 4.6 Hz, 1H), 8.10 (dd, *J* = 9.2, 5.8 Hz, 1H), 8.00 (dd, *J* = 10.9, 3.1 Hz, 1H), 7.77 (d, *J* = 8.5 Hz, 1H), 7.69 (td, *J* = 8.6, 2.8 Hz, 1H), 7.59 (d, *J* = 4.7 Hz, 1H), 7.52 – 7.46 (m, 2H), 7.37 (dd, *J* = 8.6, 2.3 Hz, 1H), 7.22 – 7.17 (m, 1H), 7.07 (d, *J* = 8.7 Hz, 1H), 6.68 (dt, *J* = 7.8, 3.8 Hz, 1H), 5.09 (dd, *J* = 12.8, 5.4 Hz, 1H), 4.58 (s, 1H), 4.23 (d, *J* = 13.8 Hz, 1H), 3.83 (s, 1H), 3.76 – 3.14 (m, 7H, hidden under water peak), 2.94 – 2.81 (m, 4H), 2.63 – 2.53 (m, 2H), 2.15 – 1.44 (m, 19H), 1.13 (d, *J* = 6.6 Hz, 3H) ppm. [M + H]^+^ calcd. for C_49_H_55_FN_7_O_7_ 872.4147; found, 872.4148.

### 5-fluoro-2-(1-methyl-2,6-dioxopiperidin-3-yl)isoindoline-1,3-dione (49)

To a solution of **34** (1.5 g, 5.4 mmol, 1.0 equiv) in DMF (12 mL) at 12 was added iodomethane (2.3 g, 1.0 mL, 16 mmol, 3.0 equiv) followed by K_2_CO_3_ (1.1 g, 8.1 mmol, 1.5 equiv). The resulting mixture was allowed to stir at 23 °C for 12 h. The reaction mixture was poured into cold DI water and the resulting solid precipitate was collected via vacuum filtration with copious cold DI water washes to provide pure **49** (1.3 g, 82%) as a light purple/grey solid. Spectral data matches previously reported characterization data.^63^

### 2-(1-methyl-2,6-dioxopiperidin-3-yl)-5-(piperazin-1-yl)isoindoline-1,3-dione (50)

To a solution of **49** (700 mg, 2.41 mmol, 1.0 equiv) and DIPEA (719 mg, 3.86 mmol, 1.6 equiv) in DMSO (7 mL) was added DIPEA (935 mg, 1.26 mL, 7.24 mmol, 3.0 equiv). The resulting mixture was stirred at 100 °C for 2.5 h. The reaction mixture was allowed to cool to 23 °C and then quenched with DI water. The resulting solid precipitate was collected via vacuum filtration with copious water rinses to provide pure *tert*-butyl 4-(2-(1-methyl-2,6-dioxopiperidin-3-yl)-1,3-dioxoisoindolin-5-yl)piperazine-1-carboxylate (1.0 g, 90 %) as a yellow solid. To *tert*-butyl 4-(2-(1-methyl-2,6-dioxopiperidin-3-yl)-1,3-dioxoisoindolin-5-yl)piperazine-1-carboxylate (1.0 g, 2.0 mmol, 1.0 equiv) was added a solution of TFA:CH_2_Cl_2_ (5 mL, 1:1) at 0 °C. The reaction mixture was allowed to stir at 23 °C for 2 h. The resulting mixture was concentrated under nitrogen atmosphere to provide a crude residue. The crude residue was triturated with Et_2_O (3 × 10mL) and the resulting solid was dried under reduced pressure to provide **50** (900 mg, 90%) as a yellow solid. Spectral data matches previously reported characterization data.^64^

### 2-(1-methyl-2,6-dioxopiperidin-3-yl)-5-(4-(piperidin-4-yl)piperazin-1-yl)isoindoline-1,3-dione (51)

To a solution of **50** (250 mg, 701 μmol, 1.0 equiv) and *tert*-butyl 4-oxopiperidine-1-carboxylate (168 mg, 842 μmol, 1.2 equiv) in DMF (4 mL) was added sodium triacetoxyhydroborate (446 mg, 2.10 mmol, 3.0 equiv). The resulting mixture was stirred at 23 °C for 1.5 h. The reaction was quenched with DI water (10 mL) and then extracted with CH_2_Cl_2_ (3 × 10 mL). The organic layers were combined, washed with water (2 × 15 mL), brine (1 × 20 mL), filtered through phase separator, and concentrated *in vacuo* to provide crude product. The crude residue was triturated with Et_2_O (3 × 5 mL) and concentrated *in vacuo* to provide pure intermediate as a yellow solid. The intermediate was then taken up in a 1:1 mixture of TFA:CH_2_Cl_2_ (1 mL) at 0 °C and stirred at 23 °C overnight. The resulting reaction mixture was concentrated under nitrogen atmosphere. The resulting residue was triturated with Et_2_O (2 × 10 mL) and the resulting yellow oil (could not get solid) was dried *in vacuo* to provide **51** (300 mg, 77%) as a yellow oil.*

### 2-(4-(4-(2-(1-methyl-2,6-dioxopiperidin-3-yl)-1,3-dioxoisoindolin-5-yl)piperazin-1-yl)piperidin-1-yl)acetic acid (52)

To a solution of **51** (250 mg, 569 μmol, 1.0 equiv) in DMF (3 mL) was added *tert*-butyl 2-bromoacetate (133 mg, 88.2 μL, 683 μmol, 1.2 equiv) and DIPEA (221 mg, 297 μL, 1.71 mmol, 3.0 equiv). The resulting mixture was stirred at 23 °C overnight. The reaction mixture was quenched with DI water and extracted with CH_2_Cl_2_ (3 × 5 mL). The organic layers were combined, washed with brine (1 × 15 mL), filtered through a phase separator and concentrated *in vacuo* to provide an oil residue. The resulting residue was triturated with Et_2_O (2 × 5mL) and concentrated *in vacuo* to provide intermediate as a yellow solid. The intermediate product was dissolved in a 1:1 mixture of CH_2_Cl_2_:TFA (2 mL) at 0 °C and stirred at 23 °C for 4 h. The resulting mixture was concentrated under nitrogen atmosphere. The crude residue was triturated in Et_2_O (3 × 5 mL) and the solid product was further dried under vacuum to provide **52** (250 mg, 72%) as a yellow solid (over two steps). ^1^H NMR (500 MHz, DMSO-*d*_6_) δ 7.76 (d, *J* = 8.4 Hz, 1H), 7.47 (d, *J* = 2.4 Hz, 1H), 7.36 (dd, *J* = 8.6, 2.3 Hz, 1H), 5.16 (dd, *J* = 13.1, 5.3 Hz, 1H), 3.99 (s, 1H), 3.65 – 3.40 (m, 12H covered by the water peak), 3.01 (s, 3H), 2.95 (ddd, *J* = 17.3, 13.8, 5.4 Hz, 3H), 2.76 (ddd, *J* = 17.2, 4.6, 2.5 Hz, 1H), 2.57 (td, *J* = 13.3, 4.6 Hz, 1H), 2.22 (d, *J* = 13.1 Hz, 2H), 2.04 (dtd, *J* = 11.2, 5.5, 2.5 Hz, 1H), 1.93 (q, *J* = 13.1 Hz, 2H) ppm. ^13^C NMR (126 MHz, DMSO) δ 171.8, 169.8, 167.4, 166.9, 154.3, 133.8, 125.0, 119.7, 118.6, 108.8, 50.9, 49.4, 47.9, 44.9, 31.1, 26.6, 21.3 ppm. [M + H]^+^ calcd. for C_25_H_32_N_5_O_6_ 498.2353; found, 498.2355.

### (2R)-2-((1s,4S)-4-(6-fluoroquinolin-4-yl)cyclohexyl)-N-(3-((1-(2-(4-(4-(2-(1-methyl-2,6-dioxopiperidin-3-yl)-1,3-dioxoisoindolin-5-yl)piperazin-1-yl)piperidin-1-yl)acetyl)piperidin-4-yl)oxy)phenyl)propenamide (24)

To a solution of **52** (150 mg, 246 μmol, 1.0 equiv) and DIPEA (95.3 mg, 128 μL, 737 μmol, 3.0 equiv) in DMF (5 mL) was added HATU (74.7 mg, 197 μmol, 0.8 equiv) at 0 °C under nitrogen atmosphere. The resulting mixture was allowed to stir at 23 °C for 30 min. To the resulting mixture was added **32** (101 mg, 197 μmol, 0.8 equiv) at 0 °C. The reaction mixture was allowed to stir at 23 °C for 30 min at which time the reaction mixture was quenched with DI water (1.5 mL) and concentrated under nitrogen atmosphere. The crude product was directly purified by reverse phase HPLC eluting from 10% to 90 % MeCN in water (0.1 % TFA) to provide pure **24** (139 mg, 48%) as a yellow solid. ^1^H NMR (500 MHz, DMSO) δ 9.98 (s, 1H), 8.89 (d, *J* = 4.6 Hz, 1H), 8.11 (dd, *J* = 9.2, 5.8 Hz, 1H), 8.00 (dd, *J* = 11.0, 2.9 Hz, 1H), 7.78 (d, *J* = 8.4 Hz, 1H), 7.69 (td, *J* = 8.7, 2.8 Hz, 1H), 7.59 (d, *J* = 4.6 Hz, 1H), 7.50 (t, *J* = 2.9 Hz, 2H), 7.37 (d, *J* = 8.3 Hz, 1H), 7.20 (t, *J* = 8.1 Hz, 1H), 7.06 (d, *J* = 8.1 Hz, 1H), 6.69 (dd, *J* = 8.3, 2.5 Hz, 1H), 5.17 (dd, *J* = 13.1, 5.4 Hz, 1H), 4.62 (s, 2H), 4.32 (s, 4H), 3.88 – 3.15 (m, 8H covered by the water peak), 3.01 (s, 7H), 2.87 (dd, *J* = 11.0, 6.6 Hz, 1H), 2.81 – 2.74 (m, 1H), 2.64 – 2.53 (m, 2H), 2.38 – 2.23 (m, 2H), 2.10 – 1.53 (m, 19H), 1.13 (d, *J* = 6.6 Hz, 3H) ppm. ^13^C NMR (126 MHz, DMSO) δ 175.0, 171.8, 169.8, 167.4, 166.9, 161.1, 159.1, 158.5, 158.3, 158.0, 157.7, 157.1, 154.2, 149.3, 144.3, 140.6, 133.8, 132.1, 129.6, 127.3, 125.0, 120.0, 119.9, 119.6, 119.4, 118.8, 117.8, 115.3, 111.9, 110.3, 108.9, 107.5, 107.3, 71.2, 59.2, 55.9, 51.6, 49.5, 48.0, 44.6, 41.5, 37.6, 35.6, 31.2, 30.5, 30.0, 28.5, 27.6, 27.5, 26.6, 26.4, 23.7, 21.4, 16.2 ppm. [M + H]^+^ calcd. for C_54_H_64_FN_8_O_7_ 955.4882; found, 955.4887.

### Cell lines, cell culture methods and PROTAC treatment

The human U87 GBM cell line and human GBM43 patient-derived xenograft (PDX) line from human adults were obtained from American Type Culture Collection (ATCC). GBM6 and GBM38 were obtained from the Mayo Clinic brain tumor PDX national resource that were developed and maintained by the Sarkaria lab at Mayo Clinic. The human pediatric glioma cell line, KNS42 GBM, was obtained from Dr. Rintaro Hashizume. The human prostate cancer cell line, PC-3, and the human pancreatic cancer cell line, SW-1990, were provided by Drs. Steve Kregel and Ali Vaziri-Gohar, respectively. The human triple negative breast cancer cell line, MDA-MB-231, and the human ovarian cancer cell line, SKOV-3, were provided by Drs. Marcelo Bonini and Dr. Daniela Matei at Northwestern University, respectively. U87, GBM6, GBM38, and KNS-42 cells were cultured in complete DMEM/F12, GBM43, SW-1990, and MDA-MB-231 were cultured in DMEM, and SKOV-3 and PC-3 were cultured in RPMI. All lines were supplemented with 10% fetal bovine serum (FBS) (Gibco) and 100 U/mL penicillin and 100 μg/mL streptomycin (Thermo Fisher Scientific) at 37 °C in a humidified chamber containing 5% CO_2_ as described.^39^

### U87-HiBit-IDO1 Degradation Assay

U87 cells were CRISPR/Cas9 genome edited to introduce the HiBiT tag fused to the N-terminus of IDO1. The IDO1-HiBiT protein level was quantified using a Nano-Glo HiBiT lytic detection system following the manufacturer’s instructions (Promega). U87 CRISPR-edited IDO1-HiBit cells were seeded in 384-well plates at 20 µl and 2,000 cells per well (Day 0). Twenty to twenty-four hours after plating, cells were treated with 20 µl of human IFN-γ (PeproTech) at a final concentration of 12.5 ng/ml (Day 1). 40 nL of compounds or DMSO control were dispensed to corresponding test wells using an Echo 555 (Labcyte) 24 hours after IFN-γ treatment (Day 2). Another 24 hours later (Day 3), cells were taken out to equilibrate at room temperature for 30 minutes. 20 µl of the culture media was removed from the 40 µl total media in each well, and an equal volume of room temperature Nano-Glo HiBiT lytic mix (Promega) were added to each well. Luminescence signal was measured using Synergy Neo2 (BioTeck) plate reader and analyzed with GraphPad Prism version 10.

### Screening Compounds by western blotting

Protein samples were prepared by lysing the cells in RIPA buffer (Sigma-Aldrich, Cat# R0278) supplemented with protease and phosphatase inhibitors on ice for 30 min. Cell lysates were clarified by centrifugation at 16,000 x *g* for 15 min at 4 °C. Equal amounts of proteins as quantified by bicinchoninic acid assay (Pierce) were separated on an SDS-polyacrylamide gel electrophoresis and proteins were electrophoretically transferred onto a polyvinylidene fluoride (PVDF) membrane using a Trans-Blot Turbo Transfer System (Bio-Rad, Hercules, CA, USA). The blotted membranes were blocked for 1 h at room temperature in blocking buffer containing 5% (w/v) nonfat dry milk in Tris-buffered saline and 0.1% Tween 20 (TBS-T) followed by incubating the membranes overnight at 4 °C with primary antibodies against a target protein diluted at a standardized concentration in blocking buffer. The blots were washed three times with TBS-T and incubated for 1 h at room temperature with horseradish peroxidase-conjugated secondary antibody generated against the host antigen in which the primary antibody was generated. The protein bands were detected using an enhanced chemiluminescence reagent (SuperSignal West Femto Maximum Sensitivity substrate) and blots were visualized with Bio-Rad ImageLab software on a Bio-Rad ChemiDoc MP imaging system. All blots were stripped and re-probed with glyceraldehyde phosphate dehydrogenase (GAPDH) to ensure the proteins were loaded equally across all the samples in a particular blot. Western blotting analysis for proteins of interest used antibodies at optimized concentrations.

### Bio-layer Interferometry (BLI) assays

A FortéBio^®^ Octet K2 BLI instrument (Sartorius) available in the Keck Biophysics Facility at Northwestern University was used for monitoring the interactions of the compounds with the IDO and CRBN proteins. 6xHis tagged recombinant IDO1 protein (Active Motif, Carlsbad, CA, cat**#** 81031) was reconstituted in PBS at 80 mg/ml, activated by pre-treatment with apo-myoglobin as described,^32^ then captured on pre-hydrated Ni-NTA biosensors (Sartorius, FortéBio^®^, cat# 18-5103). The sensors loaded with IDO1 were then equilibrated in the reaction buffer (PBS plus 0.5% DMSO, pH 7.4) for 200 seconds at 30 °C. For studies of the binary complexes between the NU227326, NU227327, and IDO1, compounds were diluted in reaction buffer to obtain stock concentrations of 60 μM. For each complex, after baseline equilibration in reaction buffer, the kinetics of association were monitored by moving individual sensors into wells containing 200 μL of analyte solutions at the concentrations shown. After the association step, the sensors were moved to reaction buffer to monitor dissociation. During the entire kinetic assay, the 96-well sample plate was kept at 30 °C with shaking at 1000 rpm. Biosensors without ligand were titrated with analyte and used as a parallel reference control. Ligand-loaded biosensors without analyte were used as baseline. Double-referenced data were fitted globally with a steady state 1:1 model using the Data Analysis HT 11.0.0.50 (FortéBio) software suite.

For examining the interaction of CRBN with NU227326, NU227327, and NU227248, BLI experiments and data analysis were performed as described above, except sensors with immobilized CRBN were used. To prepare these sensors, CRBN protein (Sino Biological, Wayne, PA) was reconstituted in Acetate Buffer, pH 6, and amine-coupled to AR2G sensors (Sartorius, FortéBio^®^, cat# 18-5092) following the manufacturer recommendations.

For experiments examining the formation of ternary complex, IDO1 was immobilized on NiNTA sensors as described above from 30 μg/mL solutions. Compounds NU227326 and NU227327 at concentrations of 1.5 μM were pre-incubated in sperate reactions with 20 molar excess of CRBN.^65, 66^ After 15 minutes of equilibration at 30 °C, a dilution series was prepared. These dilutions were allowed to equilibrate for an additional 20 minutes at 30 °C before binding reactions were monitored with BLI. Separately, experiments with identical sequences and preparation were performed using compounds NU227326 and NU227327 without CRBN and with CRBN but without NU227326 and NU227327. For the ternary and binary complexes shown, each dataset was fitted globally with a 1:1 kinetic model, using the Data Analysis HT 11.0.0.50 (FortéBio) software.

### Kynurenine assay

Kynurenine levels in cell culture supernatants were measured using a modified Erhlich method.^67^ Briefly, cell culture supernatants were incubated with a 10% final concentration of trichloroacetic acid in Eppendorf tubes for 20 min at 60 °C to release kynurenine from cells and to precipitate proteins. After 20 minutes, samples were centrifuged for 20 min at 2500 × *g* and supernatants were mixed with 20 mg/mL 4-dimethylamino benzaldehyde in acetic acid (Sigma Aldrich) at a 1:1 ratio and absorbance was measured at 490 nm using a plate reader.

### RNA isolation and Gene expression analysis using Real-Time PCR

Total RNA was extracted with lysis reagent (Ambion -Invitrogen) according to the manufacturer’s instructions. First-strand cDNA synthesis was performed with 1 μg of purified RNA using a iScript cDNA systhesi kit (Bio-Rad). Then, gene expression was performed using SYBR Green chemistry (ThermoFisher) on the ABI7300 Real-Time PCR system using QuantStudio 6 and 7 Flex software (Applied Biosystems) and analyzed by ΔΔCt method. The levels of GAPDH were used to normalize mRNA levels of target genes. The sequences of the primers used in this study were previously reported.^39^

### Statistical analysis

Comparisons between two experimental groups were performed with GraphPad Prism using Student unpaired t-test, and among more than two experimental groups were analyzed using one-way ANOVA with Tukey post hoc analysis for multiple comparisons. The data are expressed as means ±SEM from three technical replicates and three independent experiments. A value of *P* < 0.05 was considered to be significant between the two groups.

### Pharmacokinetics

All of the procedures involving animals (pharmacokinetics studies and MTD toxicity studies) were conducted in accordance with the Institutional Animal Care and Use Committee (IACUC) of HD Biosciences Co., Ltd., WuXi Apptec Co., Ltd.

Compounds **NU227326** (**21**) and **NU227327** (**20**) were formulated for IP administration in mouse *in vivo* PK studies as 5.0 mg/mL solutions. Solutions were made by dissolving compound in DMSO (5%), then adding Solutol HS-15 (10%), followed by normal saline (85%) to give a yellow, clear, solution. Sample was given by IP administration at a volume of 10 mL/kg.

Female 9–10-week-old C57BL/6J mice weighing 18-23 g and three mice per time point were used. An UPLC (Waters) chromatographic system equipped with an API 6500+ mass spectrometer (Applied Biosystems, Concord, Ontario, Canada) was used for bioanalytical analyses. Chromatographic separation was achieved on a Waters Acquity UPLC BEH C18 1.7 μm, 2.1*50 mm column maintained at RT. The flow rate was maintained at 0.7 mL/min and a mobile phase gradient of H_2_O:Acetonitrile (0.1% Formic Acid) was used. Analyst 1.7 software packages (Applied Biosystems) were used to control the LC-MS/MS system, as well as for data acquisition and processing. All data were acquired using Analyst 1.7 software (Applied Biosystems). Plasma concentration versus time data was analyzed by non-compartmental approaches using PKSolver 2.0. Data were plotted using GraphPad Prism 10.

At each time point, around 100 μL of blood was collected and quickly put on ice. Around 50 μl plasma sample was collected from blood after 4 °C, 4000 rpm centrifugation, and stored at -80 °C until LC/MS analysis. All plasma and brain samples were stored at -80 °C before PK analysis. An aliquot of 20 μL plasma was spiked into a 96-well plate, and 120 μL of acetonitrile containing internal standard were added for protein precipitation. The mixture was vortexed, centrifuged at 4000 rpm for 10 min. 100 μL of supernatant was transferred into a clean 96-well plate, and 300 μL of water were added, the mixture was vortexed and 5.0 µL of the final solution was injected for LC-MS/MS analysis. For concentrated samples, 2 μL of plasma was diluted 10x then analyzed as above. Brain samples were homogenized with ice-cold phosphate buffer saline (pH 7.4) at a ratio of 2 (buffer) : 1 (tissue) (v/w). An aliquot of 50 μL homogenate was spiked into a 1.5 mL tube, and 300 μL of acetonitrile containing internal standard was added for protein precipitation. The mixture was vortexed, centrifuged at 14000 rpm for 10 min. 100 μL of supernatant was transferred into a clean 96-well plate, and 300 μL of water were added, the mixture was vortexed and 5.0 µL of the final solution was injected for LC-MS/MS analysis.

### Proteomics

#### Sample preparation LFQ quantitative mass spectrometry

Cells were lysed by addition of lysis buffer (8 M Urea, 50 mM NaCl, 50 mM 4-(2-hydroxyethyl)-1-piperazineethanesulfonic acid (EPPS) pH 8.5, Protease and Phosphatase inhibitors) and homogenization by bead beating (BioSpec) for three repeats of 30 seconds at 2400 strokes/min. Bradford assay was used to determine the final protein concentration in the clarified cell lysate. Fifty micrograms of protein for each sample was reduced, alkylated and precipitated using methanol/chloroform as previously described ^68^ and the resulting washed precipitated protein was allowed to air dry. Precipitated protein was resuspended in 4 M urea, 50 mM HEPES pH 7.4, followed by dilution to 1 M urea with the addition of 200 mM EPPS, pH 8. Proteins were digested with the addition of LysC (1:50; enzyme:protein) and trypsin (1:50; enzyme:protein) for 12 h at 37 °C. Sample digests were acidified with formic acid to a pH of 2-3 before desalting using C18 solid phase extraction plates (SOLA, Thermo Fisher Scientific). Desalted peptides were dried in a vacuum-centrifuged and reconstituted in 0.1% formic acid for liquid chromatography-mass spectrometry analysis.

Data were collected using a TimsTOF HT (Bruker Daltonics, Bremen, Germany) coupled to a nanoElute LC pump (Bruker Daltonics, Bremen, Germany) via a CaptiveSpray nano-electrospray source. Peptides were separated on a reversed-phase C_18_ column (25 cm x 75 µm ID, 1.6 µM, IonOpticks, Australia) containing an integrated captive spray emitter. Peptides were separated using a 50 min gradient of 2 -30% buffer B (acetonitrile in 0.1% formic acid) with a flow rate of 250 nL/min and column temperature maintained at 50 °C.

The TIMS elution voltages were calibrated linearly with three points (Agilent ESI-L Tuning Mix Ions; 622, 922, 1,222 *m/z*) to determine the reduced ion mobility coefficients (1/K_0_). To perform diaPASEF, we used py_diAID ^69^, a python package, to assess the precursor distribution in the *m/z*-ion mobility plane to generate a diaPASEF acquisition scheme with variable window isolation widths that are aligned to the precursor density in m/z. Data was acquired using twenty cycles with three mobility window scans each (creating 60 windows) covering the diagonal scan line for doubly and triply charged precursors, with singly charged precursors able to be excluded by their position in the m/z-ion mobility plane. These precursor isolation windows were defined between 350 -1250 *m/z* and 1/k0 of 0.6 -1.45 V.s/cm^2^.

#### LC-MS data analysis

The diaPASEF raw file processing and controlling peptide and protein level false discovery rates, assembling proteins from peptides, and protein quantification from peptides were performed using library free analysis in DIA-NN 1.8 ^70^. Library free mode performs an in silico digestion of a given protein sequence database alongside deep learning-based predictions to extract the DIA precursor data into a collection of MS2 spectra. The search results are then used to generate a spectral library which is then employed for the targeted analysis of the DIA data searched against a Swissprot human database (January 2021). Database search criteria largely followed the default settings for directDIA including: tryptic with two missed cleavages, carbamidomethylation of cysteine, and oxidation of methionine and precursor Q-value (FDR) cut-off of 0.01. Precursor quantification strategy was set to Robust LC (high accuracy) with RT-dependent cross run normalization. Proteins with low sum of abundance (<2,000 x no. of treatments) were excluded from further analysis and resulting data was filtered to only include proteins that had a minimum of 3 counts in at least 4 replicates of each independent comparison of treatment sample to the DMSO control. Protein with missing values were imputed by random selection from a Gaussian distribution either with a mean of the non-missing values for that treatment group or with a mean equal to the median of the background (in cases when all values for a treatment group are missing) using in-house scripts in the R framework (R Development Core Team, 2014). Significant changes comparing the relative protein abundance of these treatment to DMSO control comparisons were assessed by moderated t test as implemented in the limma package within the R framework.^71^

## ASSOCIATED CONTENT

### Supporting Information

This material is available free of charge via the Internet at http://pubs.acs.org. SMILES strings table of the compounds described here and their IDO1 DC_50_ (CSV).

Kynurenine inhibition data for compound **21** (**NU227326**), global quantitative proteomics data for **NU227326** and **NU227327**, BLI data for inactive control compound **24** (**NU227428**), pharmacokinetics data for **20** (**NU227327**), ^1^H-NMR, ^13^C-NMR, and HPLC of key final compounds, raw western blot images.

## Author Contributions

The manuscript was written through contributions of all authors. All authors read and intellectually contributed to the manuscript. *P.J.M. and *P.V.B. contributed equally.

## ACKNOWLEDGEMENT

This work was supported in part by National Institutes of Health (NIH) grants R01NS097851 (D.A.W. and G.E.S.), K02AG068617 (D.A.W.), R01NS129835 (D.A.W.), American Cancer Scholar Research Scholar Award RSG-21-058-01 -CCE (D.A.W.), and the GBM Foundation (D.A.W.). This work was supported by the Northwestern University High Throughput Analysis Core, which received funding from the Lurie Cancer Center (NCI grant CA060553) and the Acoustic liquid handler SIG (NIH S10OD023681). This work made use of the IMSERC NMR facility at Northwestern University, which has received support from the Soft and Hybrid Nanotechnology Experimental (SHyNE) Resource (NSF ECCS-2025633), the State of Illinois, the International Institute for Nanotechnology (IIN), and Northwestern University. Some of the work was completed in the Keck Biophysics Facility, a shared resource of the Robert H. Lurie Comprehensive Cancer Center of Northwestern University supported in part by the NCI Cancer Center Support Grant P30CA060553. We thank Katherine A. Donovan, Eric S. Fischer and the Fischer Lab Degradation Proteomics Initiative for collection of the global proteomics data supported by NIH CA214608 and CA218278. Global proteomics data will be publicly available at the Fischer Lab’s Proteomics database: https://proteomics.fischerlab.org.

## Abbreviations Used

AR2G: Amine reactive 2^nd^ generation
BLI: Biolayer Interferometry
CRBN: Cereblon
CH_2_Cl_2_: dichloromethane
DIPEA: diisopropylethylamine
DMF: *N*,*N*-Dimethylformamide
DMSO: dimethylsulfoxide
EDCI: 1-Ethyl-3-(3-dimethylaminopropyl)carbodiimide
GAPDH: Glyceraldehyde phosphate dehydrogenase
GBM: Glioblastoma
HATU: Hexafluorophosphate Azabenzotriazole Tetramethyl Uronium
ICB: Immune checkpoint blockade
IDO1: Indoleamine 2,3-dioxygenase 1
IFNγ: Interferon gamma
ITC: Isothermal titration calorimetry
KYN: Kynurenine
MeOH: methanol
PROTAC: Proteolysis Targeting Chimera
SAR: Structure-Activity relationship
STAB: sodium triacetoxyborohydride
*t*-BuOK: potassium tert-butoxide
THF: tetrahydrofuran
Treg: Regulatory T-cell
Trp: Tryptophan
T3P: Propanephosphonic acid anhydride
VHL: Von Hippel-Lindau

---

**Table of Contents Graphic.**
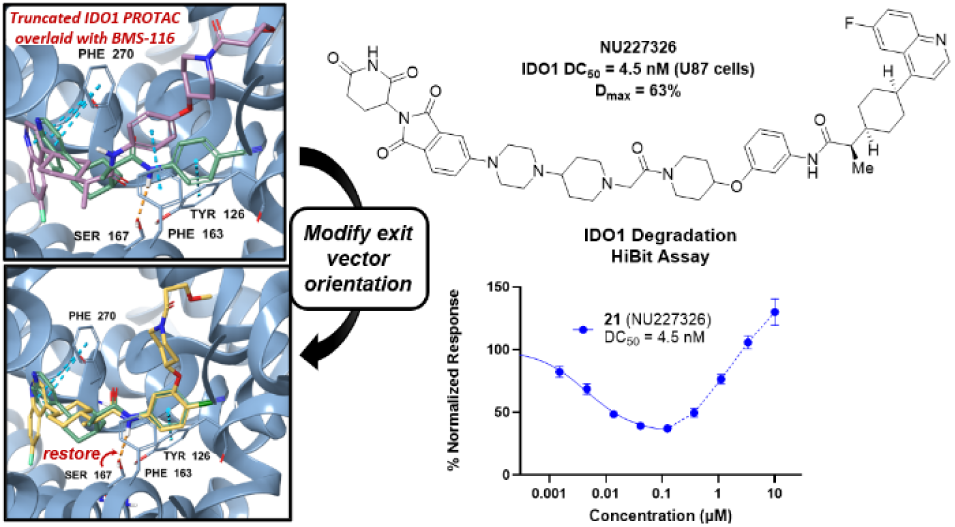

