## Supplementary Information for "Rational Design and Optimization of a Potent IDO1 Proteolysis Targeting Chimera (PROTAC)"

<u>**Supporting Information**</u>

**Affiliations:** [1]Department of Chemistry, Northwestern University, Evanston, IL, USA. [2]Department of Cancer Biology, Loyola University Chicago Stritch School of Medicine, Maywood, IL, USA. [3]Department of Molecular Biosciences, Northwestern University Weinberg College of Arts and Sciences, Evanston, IL, USA. [4]High-Throughput Analysis Laboratory, Chemistry of Life Processes Institute, Northwestern University, Evanston, IL, USA. [5]HD Biosciences (China) Co. Ltd, a WuXi AppTec company, Shanghai 201201, China. [6]Cardinal Bernardin Cancer Center, Maywood, IL, USA. [7]Department of Surgery, Loyola University Chicago Stritch School of Medicine, Maywood, IL, USA. [8]Neuro-oncology Unit, Unidad Funcional de Investigación en Enfermedades Crónicas (UFIEC), Instituto de Salud Carlos III (ISCIII), Madrid, Spain. [9]Department of Health and Kinesiology, University of Illinois at Urbana-Champaign, Urbana, IL, USA. [10]Department of Neurological Surgery, University of Chicago Medicine, Chicago, IL, USA. [11]Department of Neurology, Northwestern University Feinberg

School of Medicine, Chicago, IL, USA. [12]Department of Neurological Surgery, Loyola University Medical Center, Maywood, IL, USA. [13]Robert H. Lurie Comprehensive Cancer Center, Chicago, IL, USA. [14]Department of Pharmacology, Northwestern University, Feinberg School of Medicine, Chicago, IL, USA.

**Table of Contents**

Supplementary Figure 1. Kynurenine inhibition data of **21** (**NU227236**)………………..……...4

Supplementary Figure 2. Global quantitative proteomics of NU227326 and NU227327…........5-6

Supplementary Figure 3. BLI data for Inactive IDO1 PROTAC **24** (**NU227428**).........................7

Supplementary Figure 4. Pharmacokinetics of **NU227327** (**20**)....…………………………..……8

$^{1}$H-NMR, $^{13}$C-NMR, and HPLC of key IDO1 Degraders…………………...……………...…9-28

Raw western blot images…………………………...…………………………………...…29-30

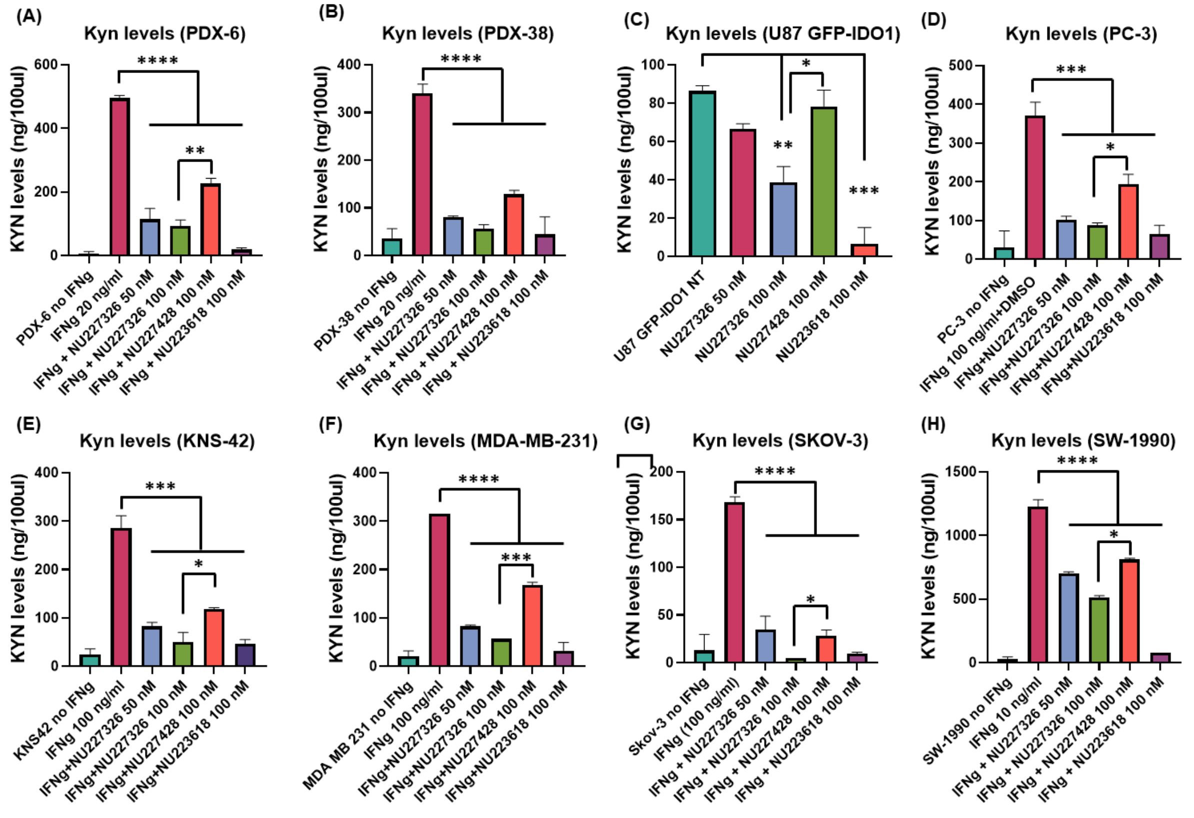


**Supplementary Figure 1**. **Inhibition of IDO1 enzyme activity by 21 (NU227326) in human cancer cell lines.** Supernatants from various human cancer cell lines, including adult GBM patient-derived xenograft cells PDX-6 (A) and PDX-38 (B), U87 cells overexpressing GFP-fused IDO1 (C), prostate cancer PC-3 cells (D), pediatric GBM KNS42 cells (E), triple-negative breast cancer MDA-MB-231 cells (F), pancreatic cancer SW-1990 cells (G), and ovarian cancer SKOV-3 cells (H), were collected after 24-hour treatment with either compound 21 (NU227326), mutant PROTAC 28 (NU227428), or parental compound 4 (NU223618) at indicated concentrations. Kynurenine levels in the supernatants were measured using a modified Ehrlich method. Where indicated, cells were also treated with IFNγ at specified concentrations. Data are presented as mean ± SEM. Statistical significance was determined using Tukey's multiple comparison test for comparisons between more than two groups and an unpaired Student's t-test for comparisons between two groups. Significance levels are indicated as follows: $P < 0.05$ (*), $P < 0.01$ (**), $P < 0.001$ (***), and $P < 0.0001$ (****).

**Quantitative proteomics of NU227326**

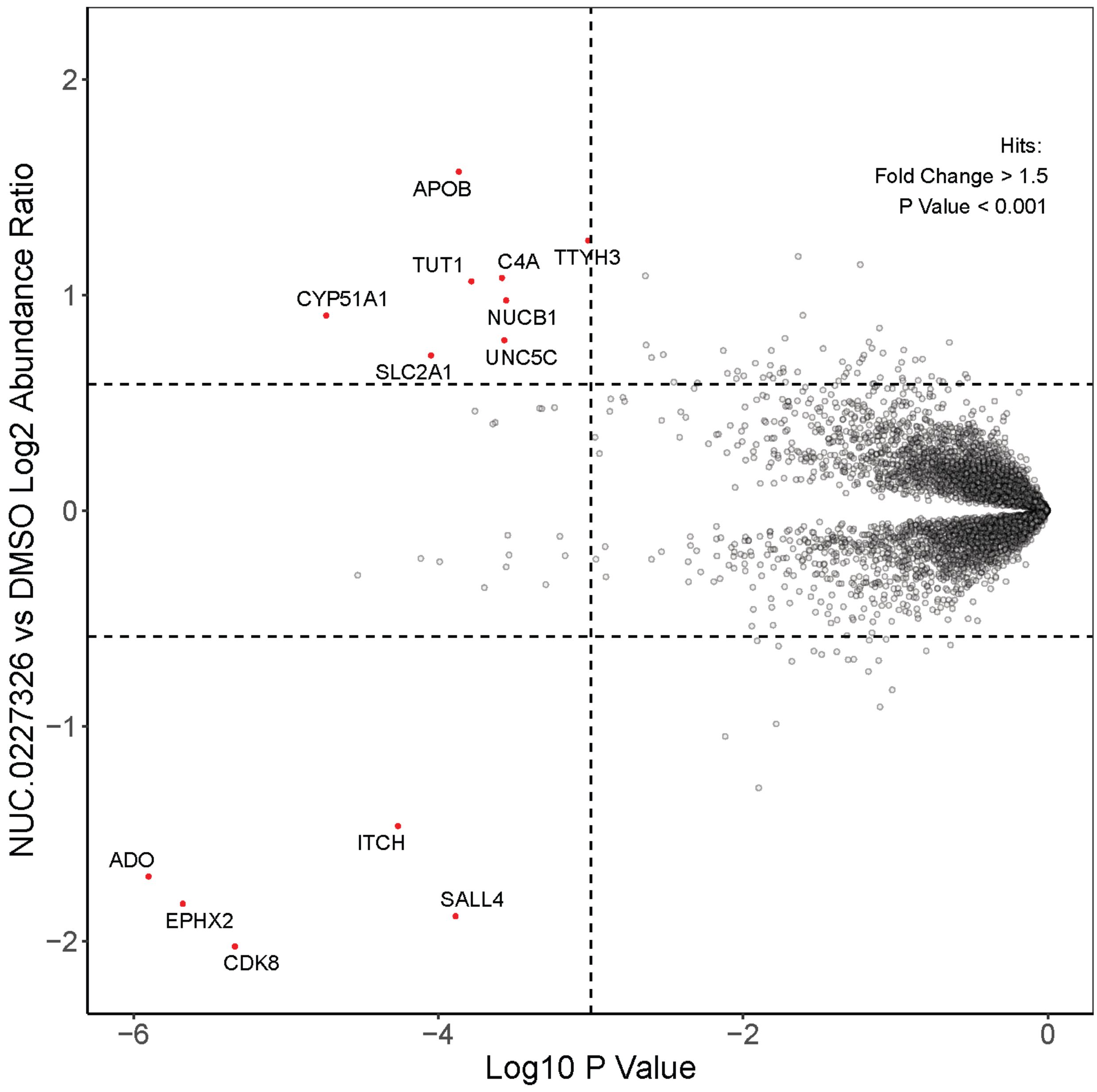

**Quantitative proteomics of NU227327**

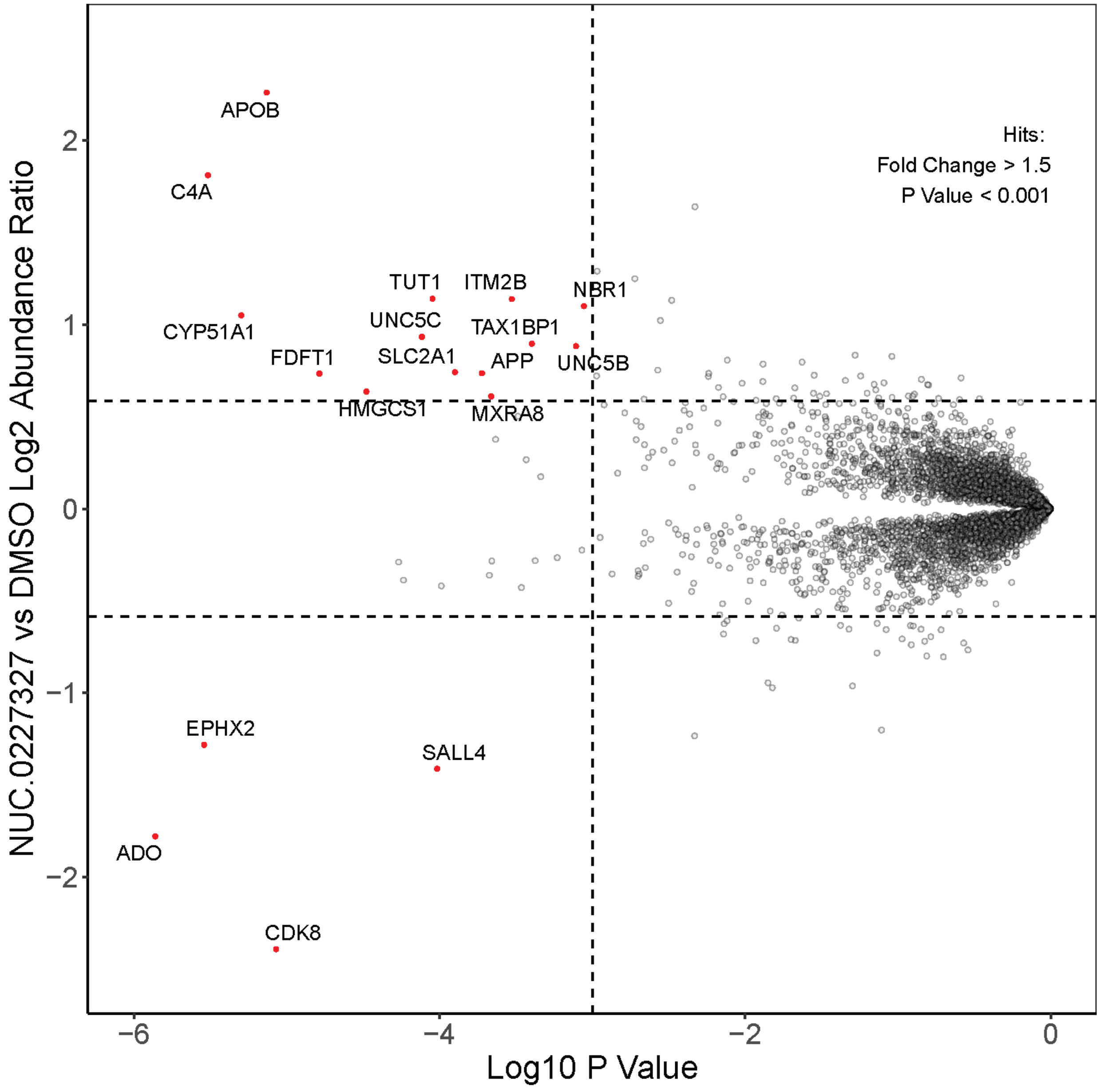


**Supplementary Figure 2**. Global quantitative proteomics of **NU227326** and **NU222737**. Kelly cells were treated with 1μM compound for 24 hrs and analyzed as described in the Experimental section.

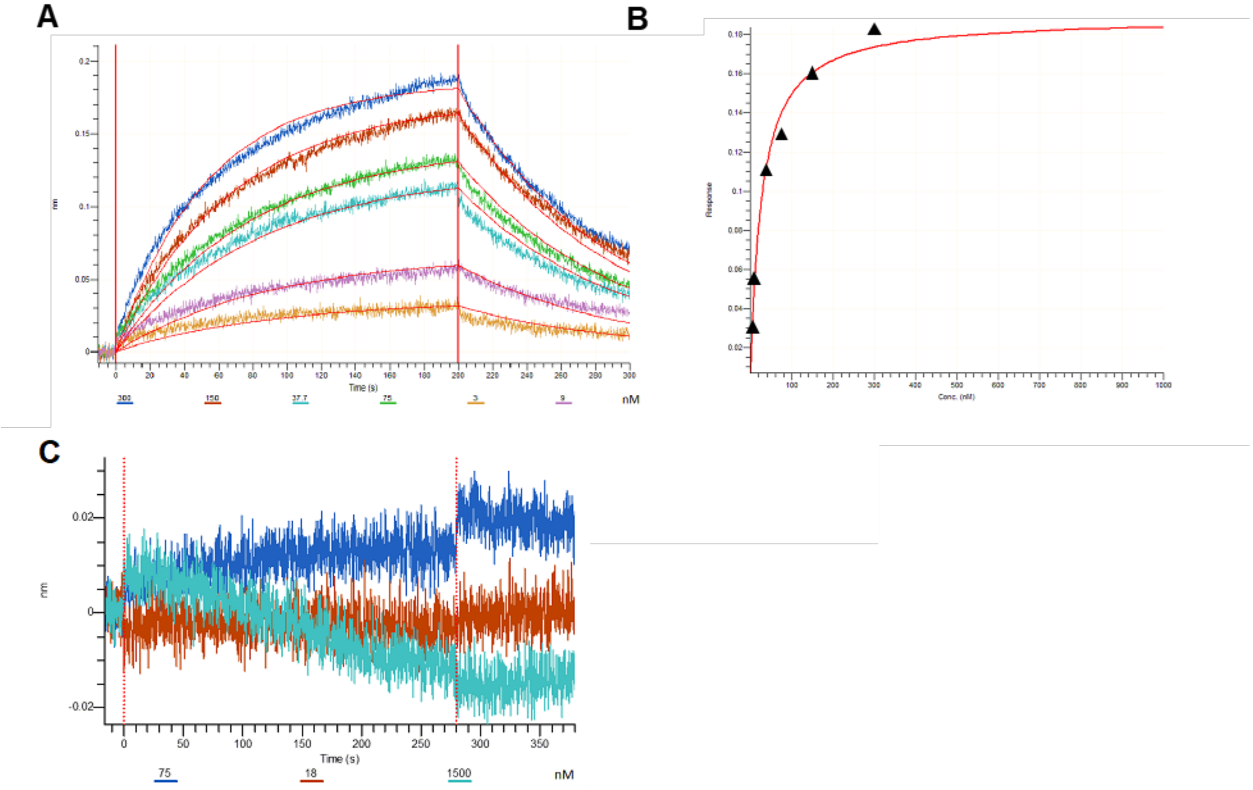


**Supplementary Figure 3**. BLI data for Inactive IDO1 PROTAC **24** (**NU227428**). (A) BLI sensograms showing association and dissociation of compound **24** (**NU227428**) to IDO1 protein (loaded on NI NTA sensors). (B) The equilibrium dissociation constant (Kd) was obtained by fitting the steady state data (Req as a function of compound concentration) with a 1:1 binding model. The measured affinity (standard deviation) for the binary complex formed by IDO1 with **24** (**NU227428**) was determined to be 260 nM. (C) BLI sensorgrams monitoring the interaction of compound **24** (**NU227428**) with CRBN. CRBN was immobilized on AR2G sensors; the sensors were equilibrated in reaction buffer and then dipped into solutions of **24** (**NU227428**) (at concentrations indicated in the legend). No association between CRBN and **24** (**NU227428**) could be detected in these experiments.

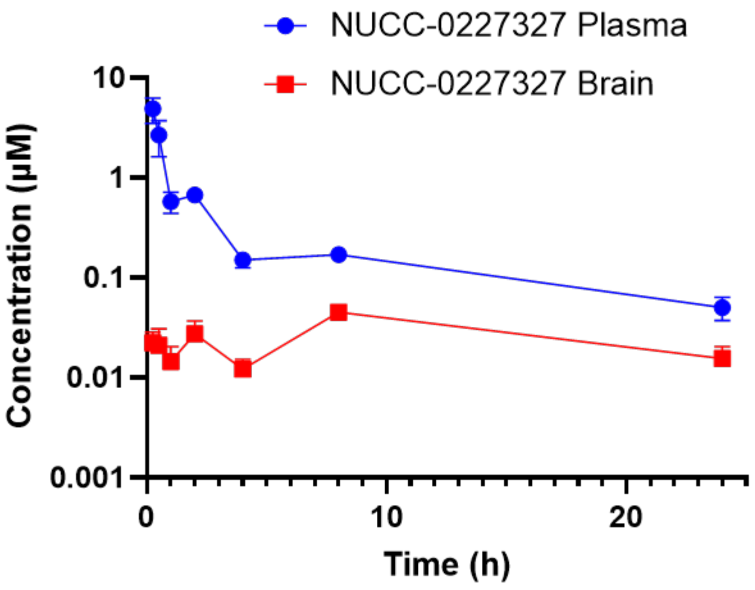


| Compound | Dose (mg/kg) | $T_{1/2}$ (h) | $T_{max}$ (h) | $C_{max}$ (μM) | $AUC_{0-\infty}$ (μM·h) | $AUC_{0-t}$ (μM·h) | $MRT_{INF}$ (h) | Vz/F_Obs (mg/kg)/(μM) | CL/F_obs (mg/kg)/(μM)/h |
|---|---|---|---|---|---|---|---|---|---|
| NUCC-0227327 (plasma) | 50 | 11.4 | 0.25 | 4.9 | 7.1 | 6.2 | 8.8 | 116.5 | 7.1 |
| NUCC-0227327 (brain) | 50 | - | 8.0 | 0.045 | - | 0.68 | - | - | - |

**Supplementary Figure 4**. Pharmacokinetics of **NU227327** (**20**). Several parameters could not be calculated for the brain samples because three points were not detected in the terminal elimination phase.

#### $^{1}H$-NMR, $^{13}C$-NMR, and HPLC of key IDO1 Degraders

#### $^{1}H$-NMR and $^{13}C$-NMR Spectra of PROTAC 7

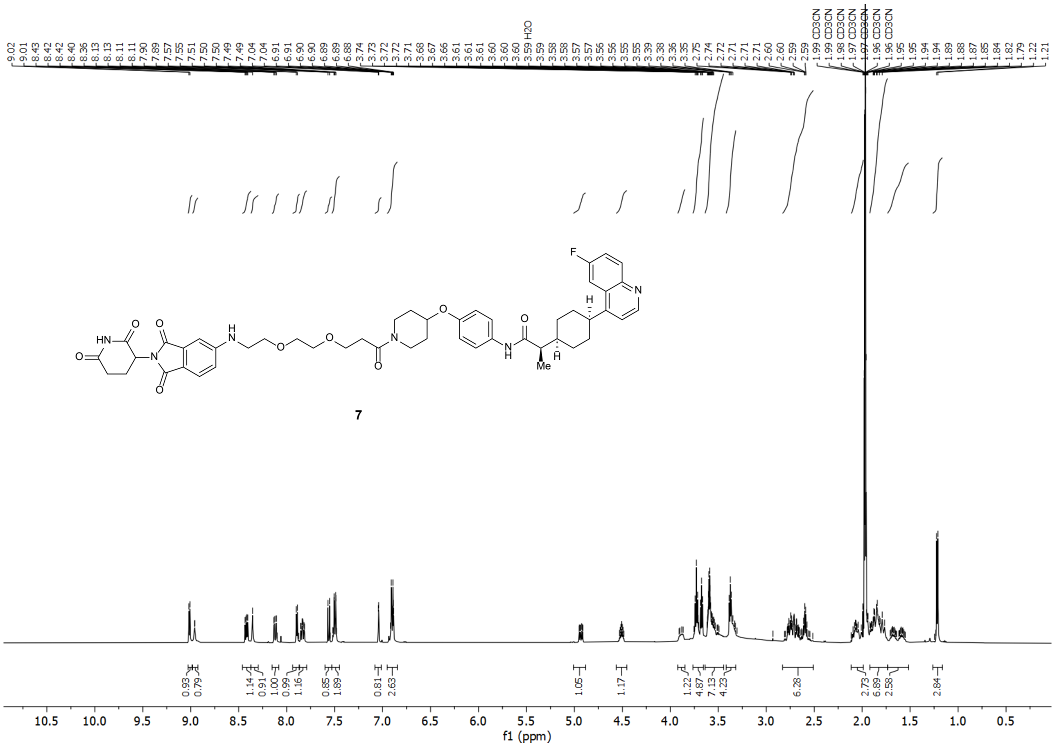


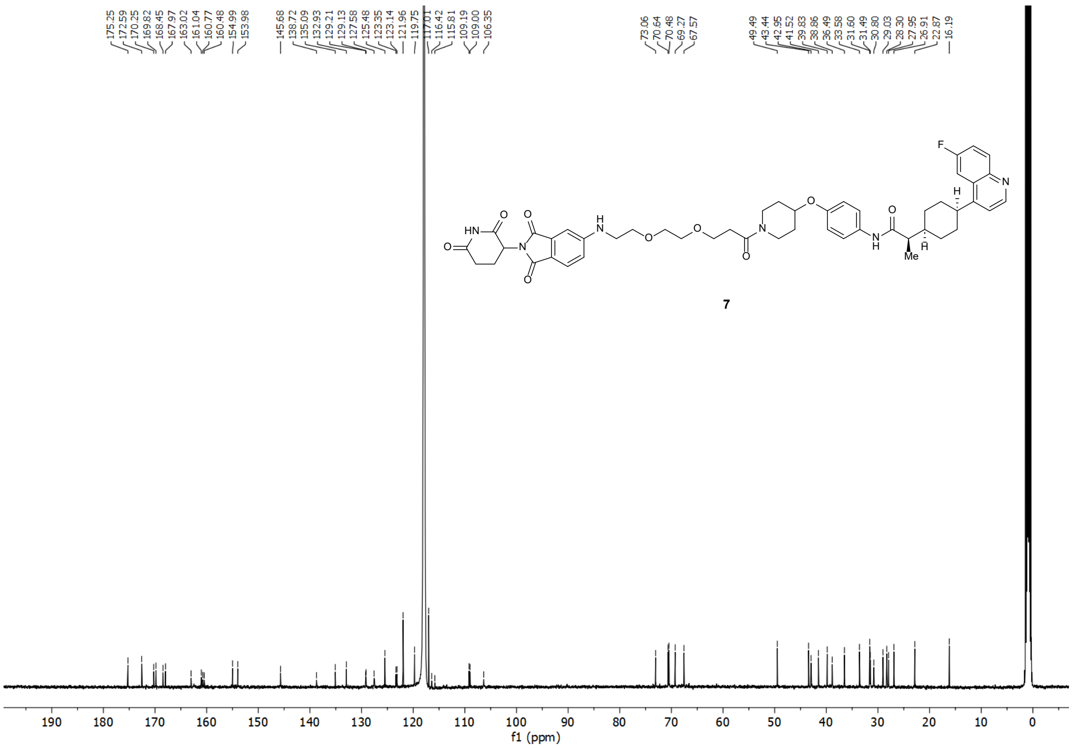

### $^{1}$H-NMR and $^{13}$C-NMR Spectra of PROTAC 8

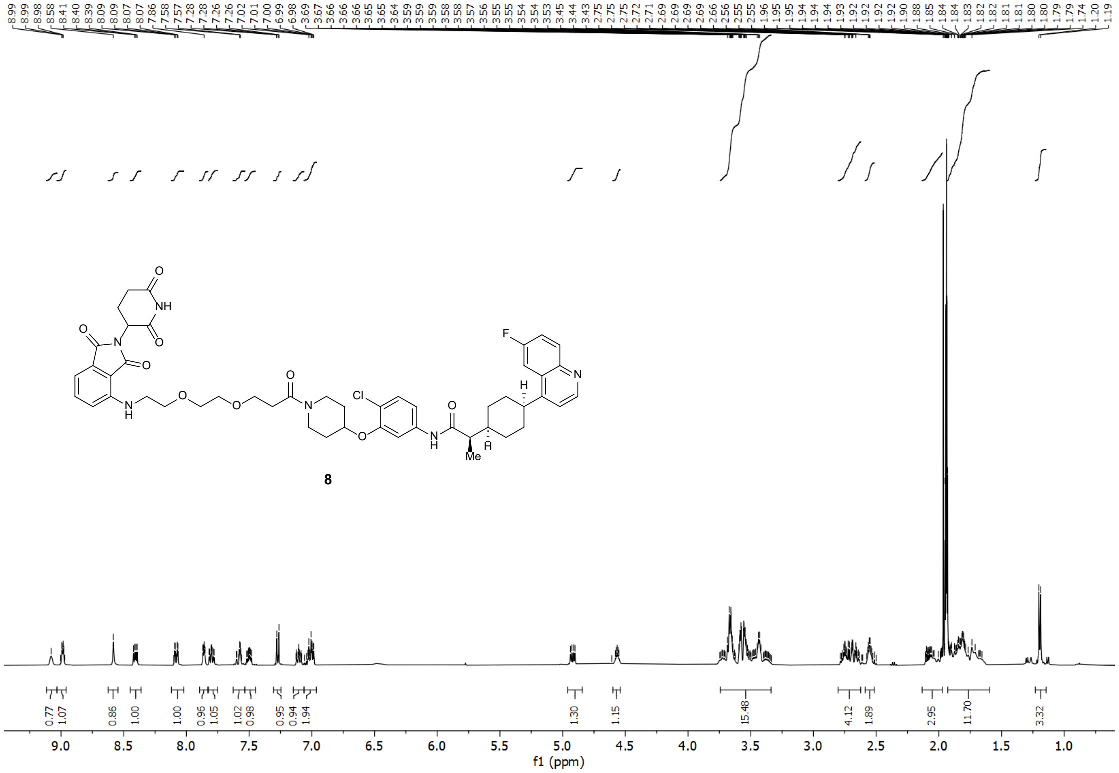


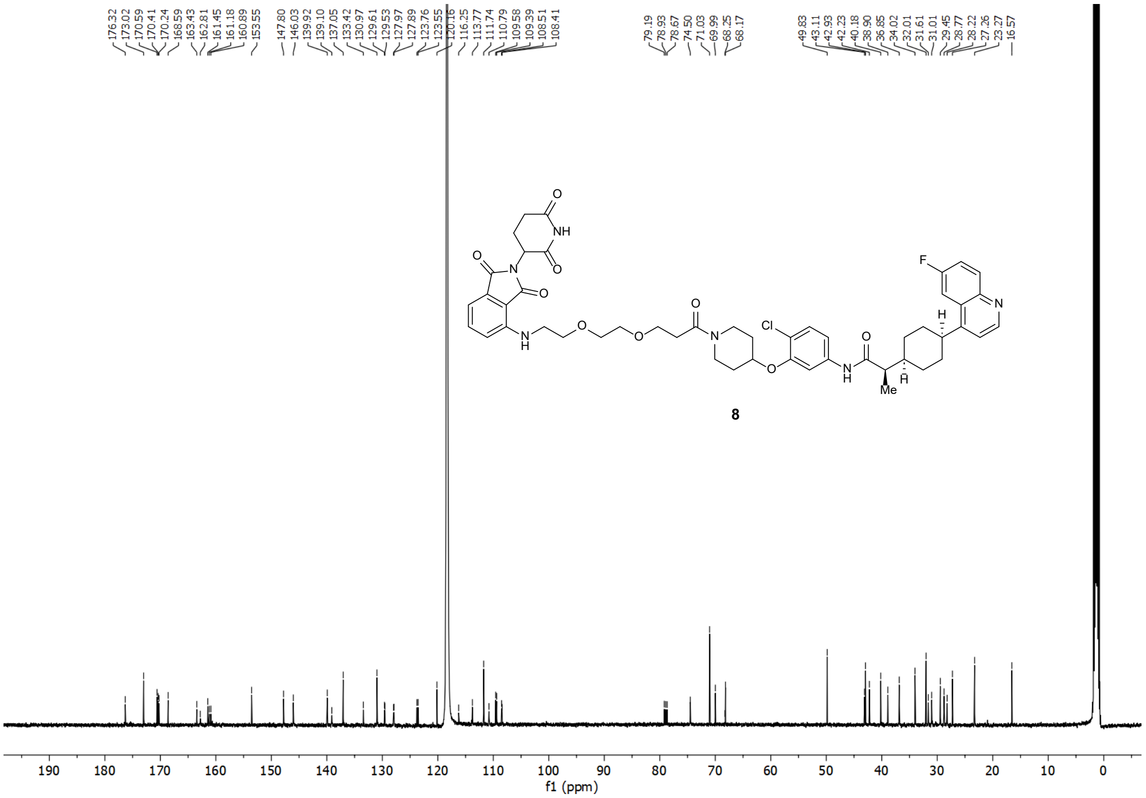

#### [1]H-NMR and [13]C-NMR Spectra of PROTAC 9

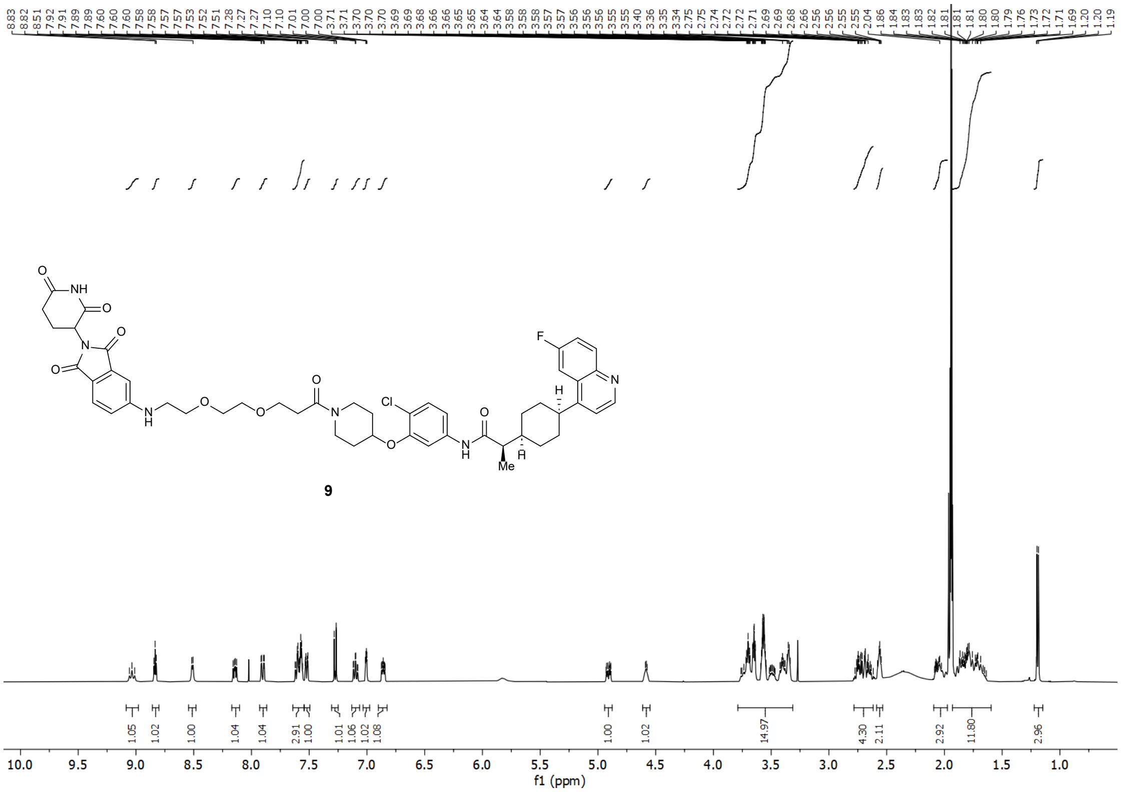


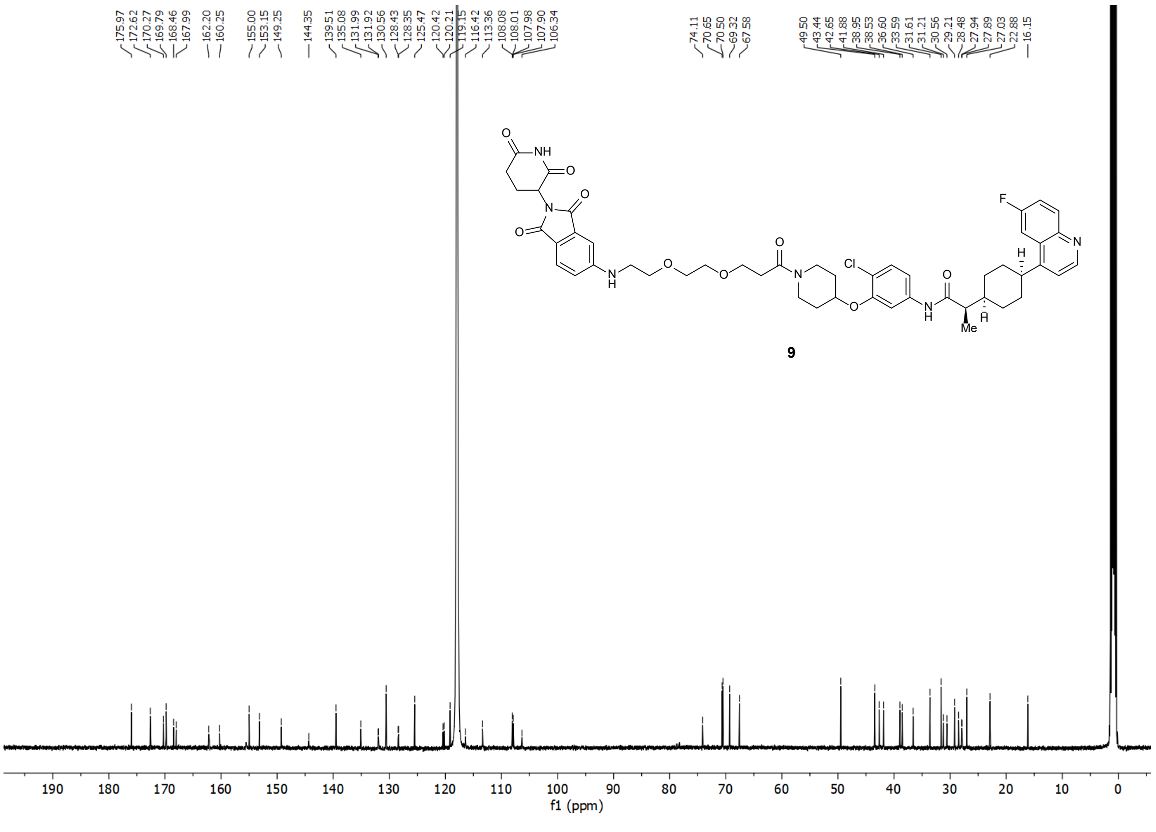

### [1]H-NMR and [13]C-NMR Spectra of PROTAC 10

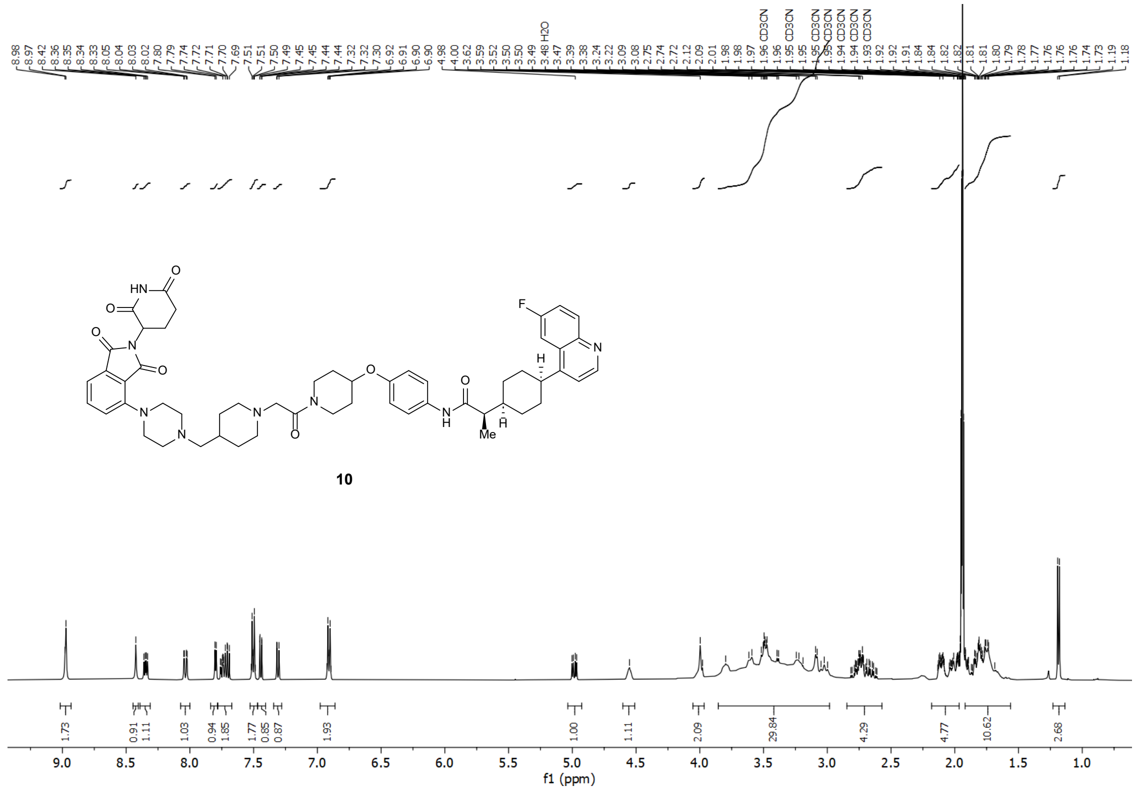


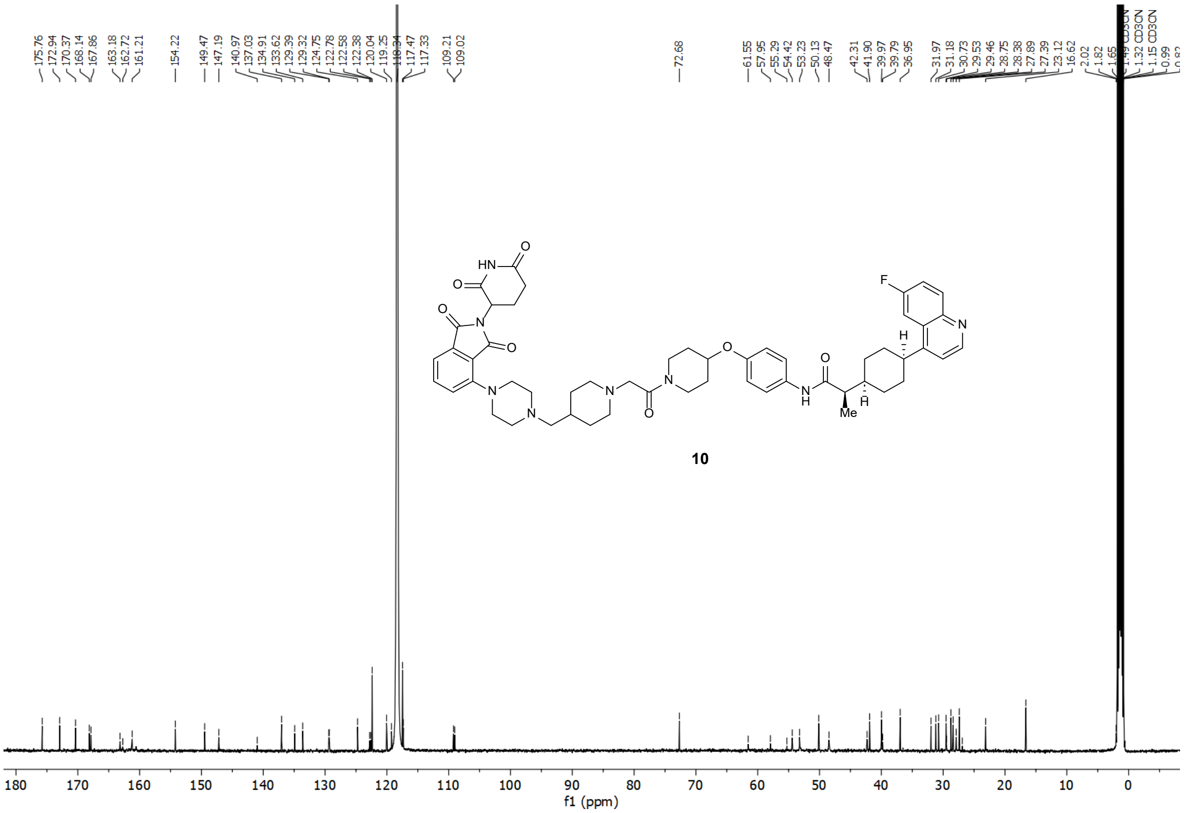

### $^{1}$H-NMR and $^{13}$C-NMR Spectra of PROTAC 11

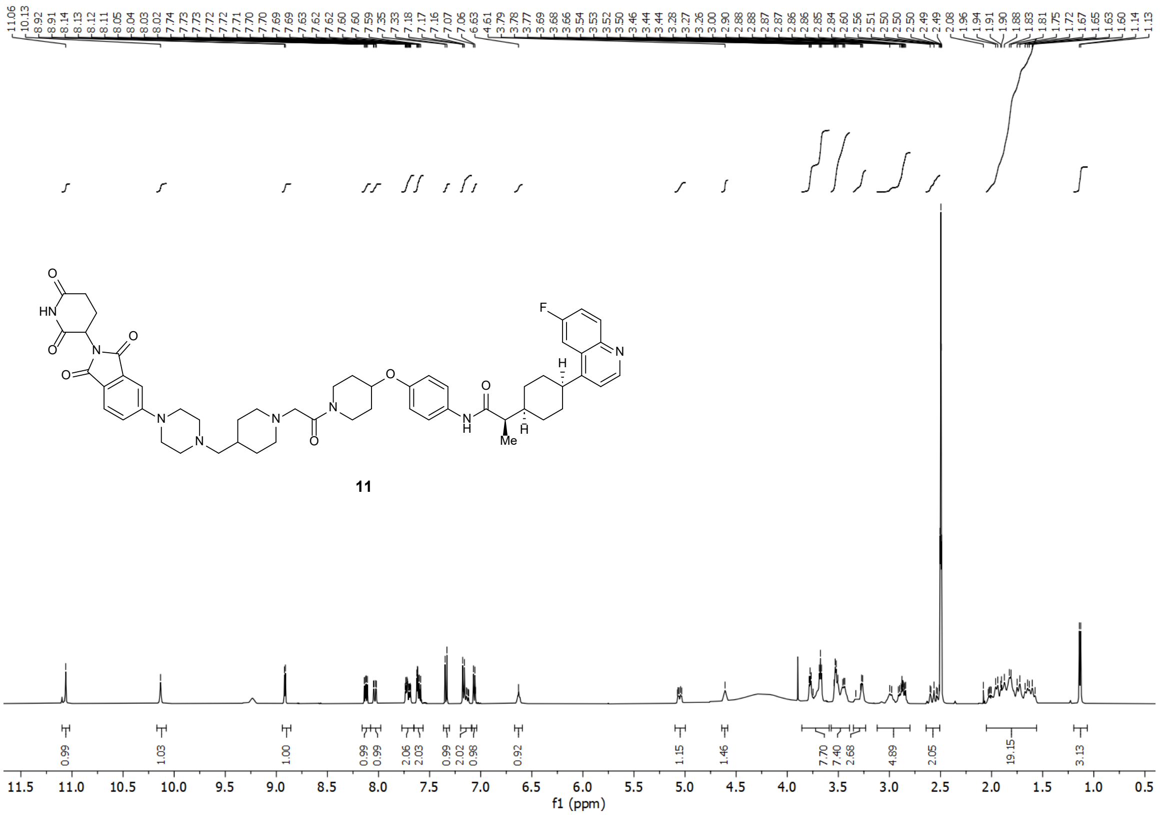


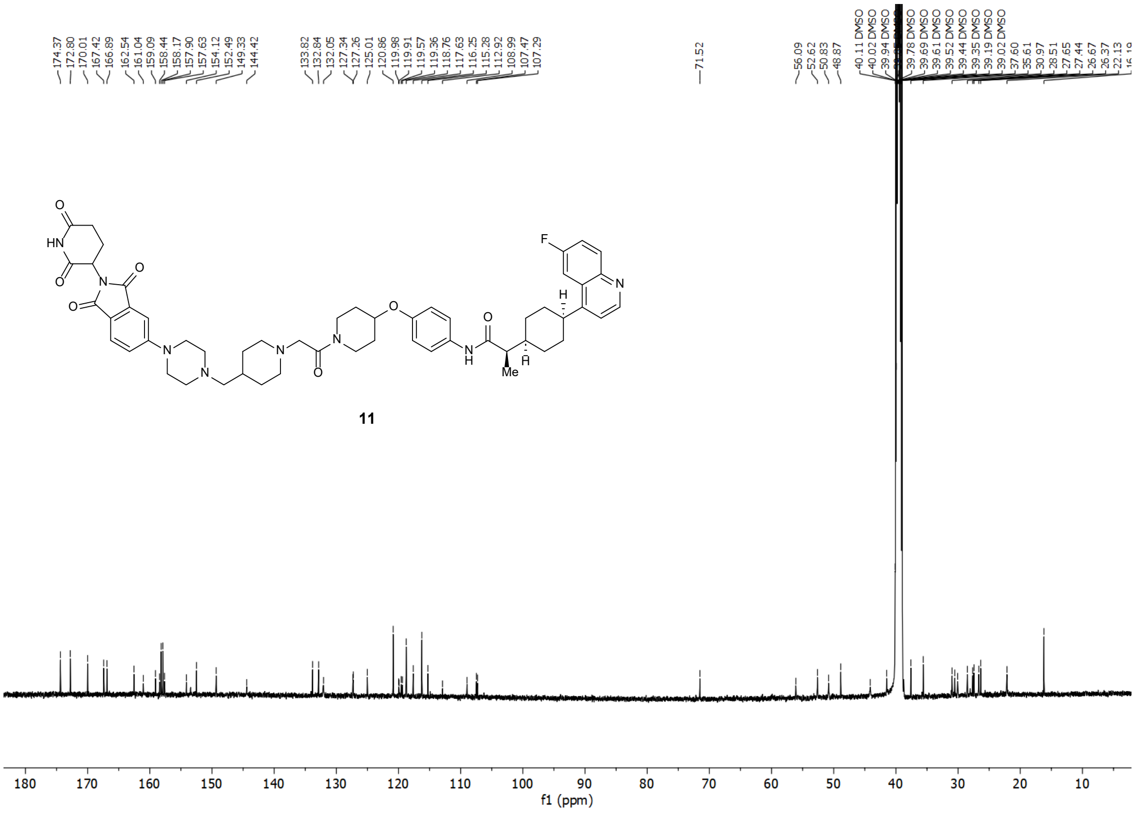

### [1]H-NMR and [13]C-NMR Spectra of PROTAC 12

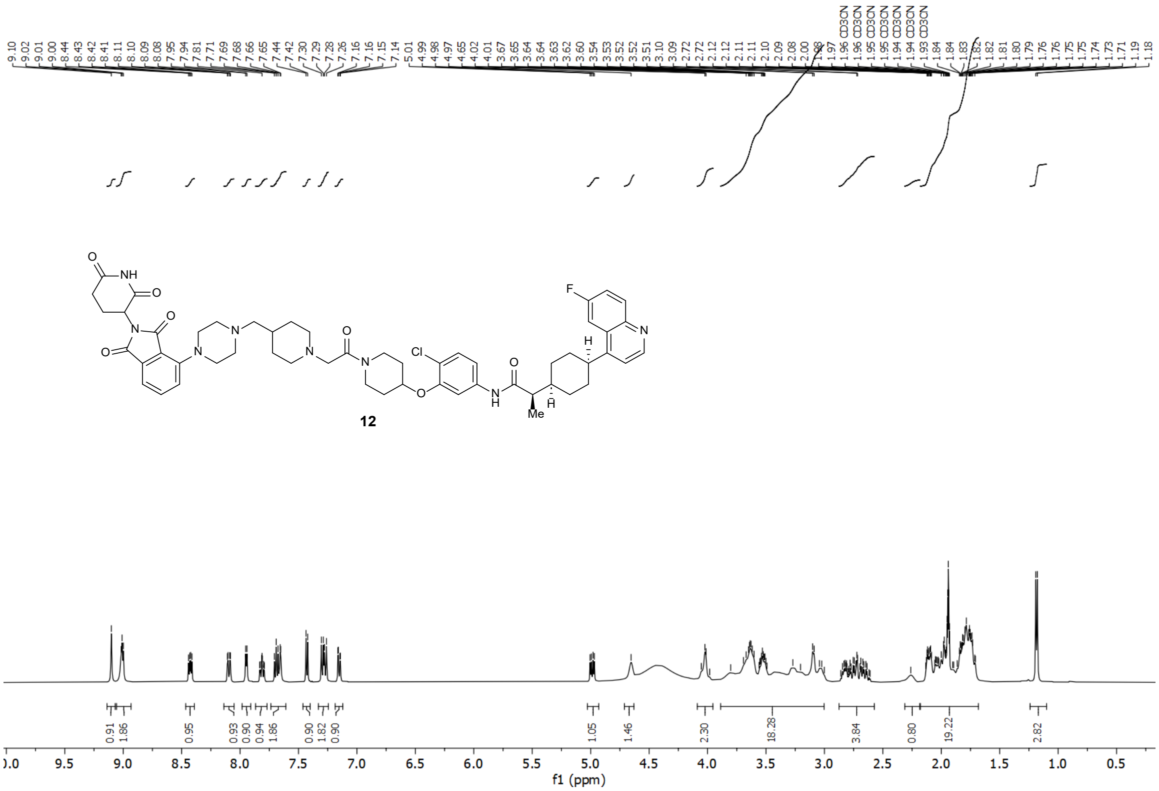


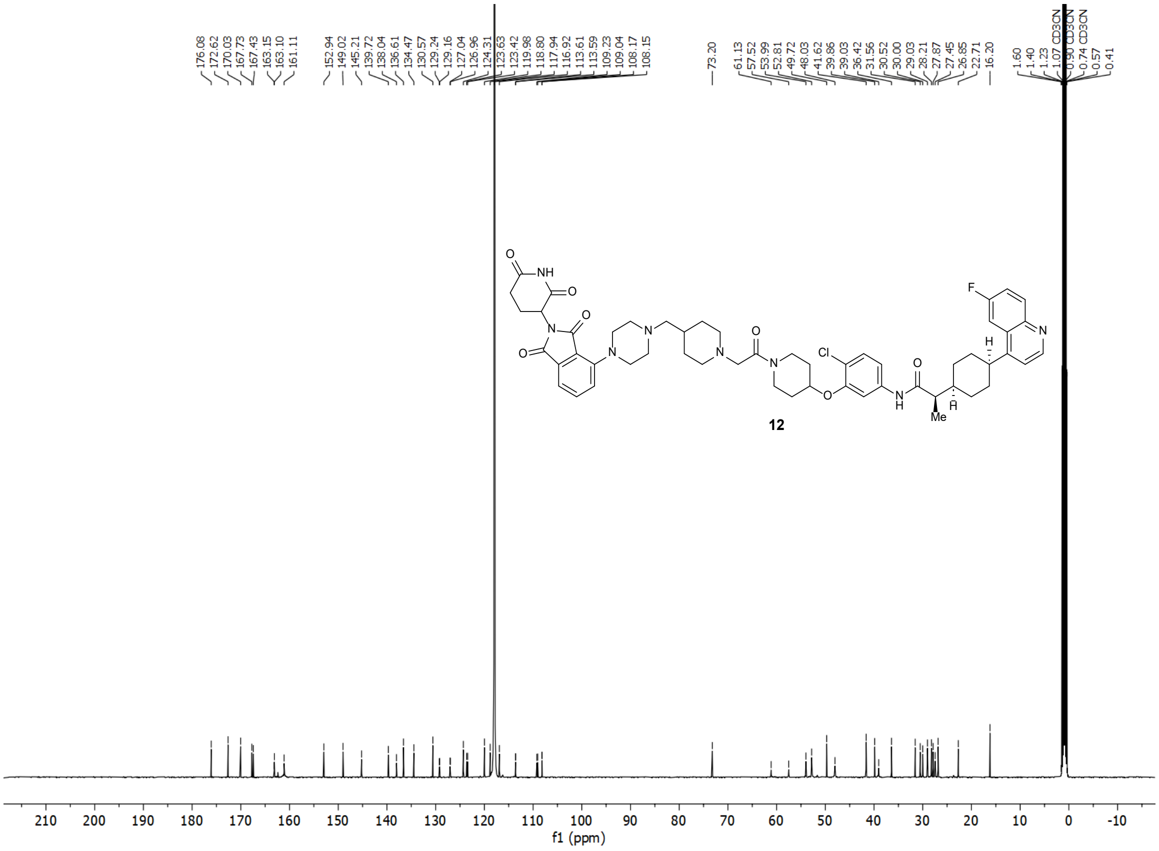

### [1]H-NMR and [13]C-NMR Spectra of PROTAC 13

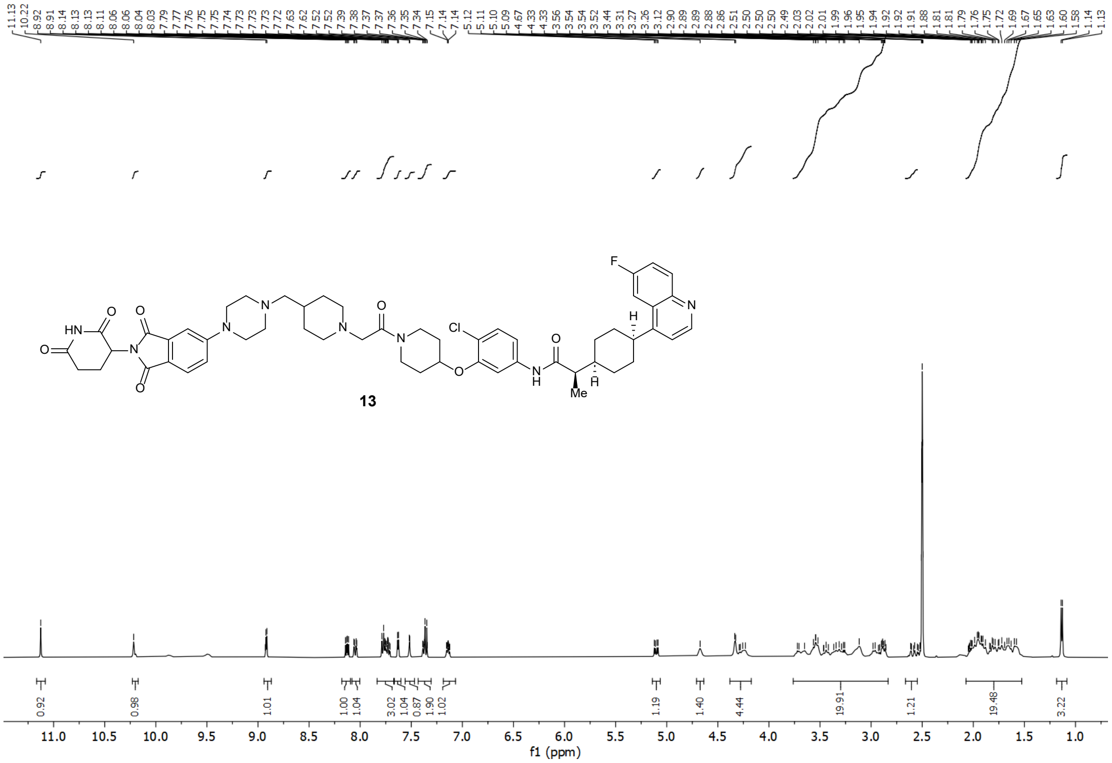


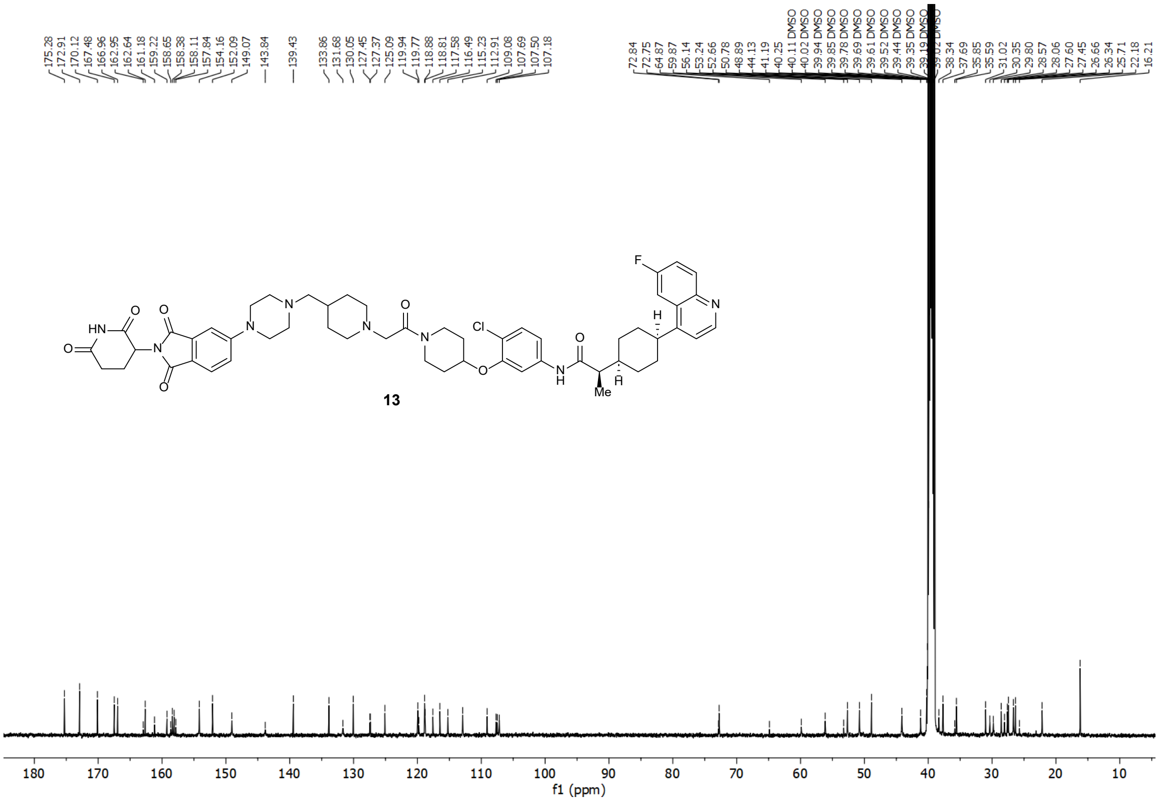

#### [1]H-NMR and [13]C-NMR Spectra of PROTAC 14

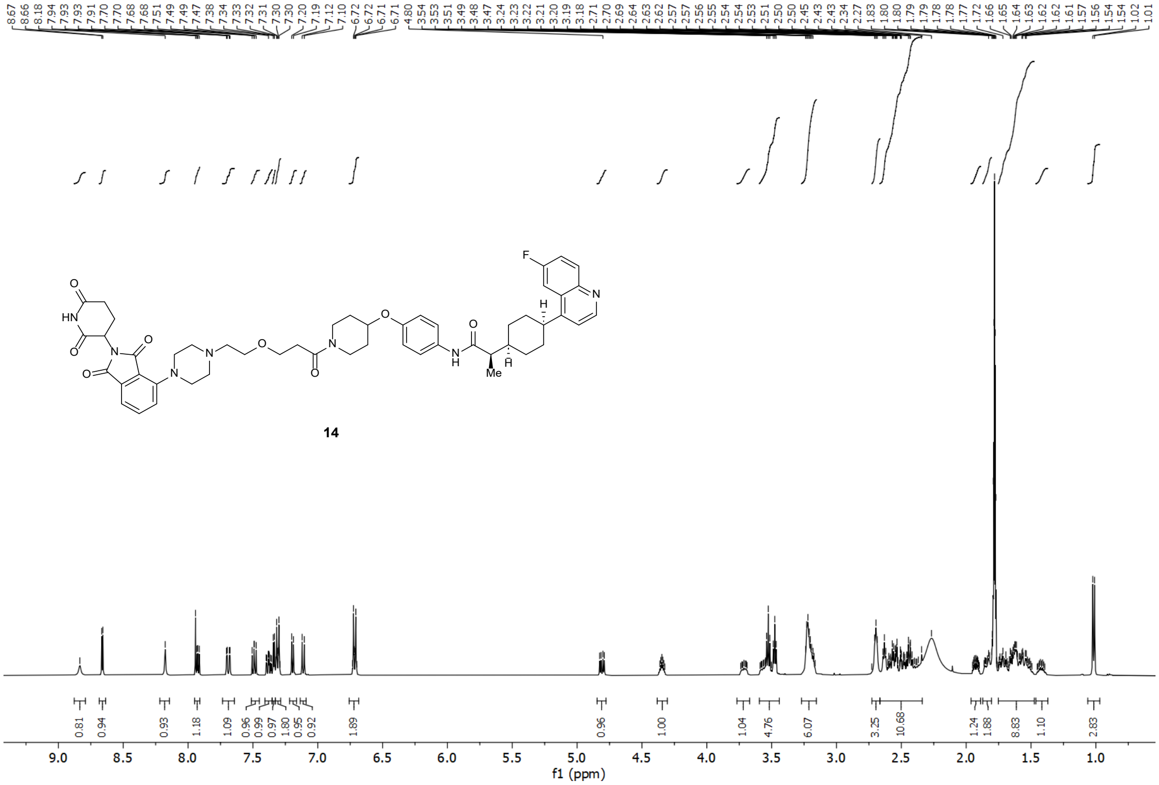


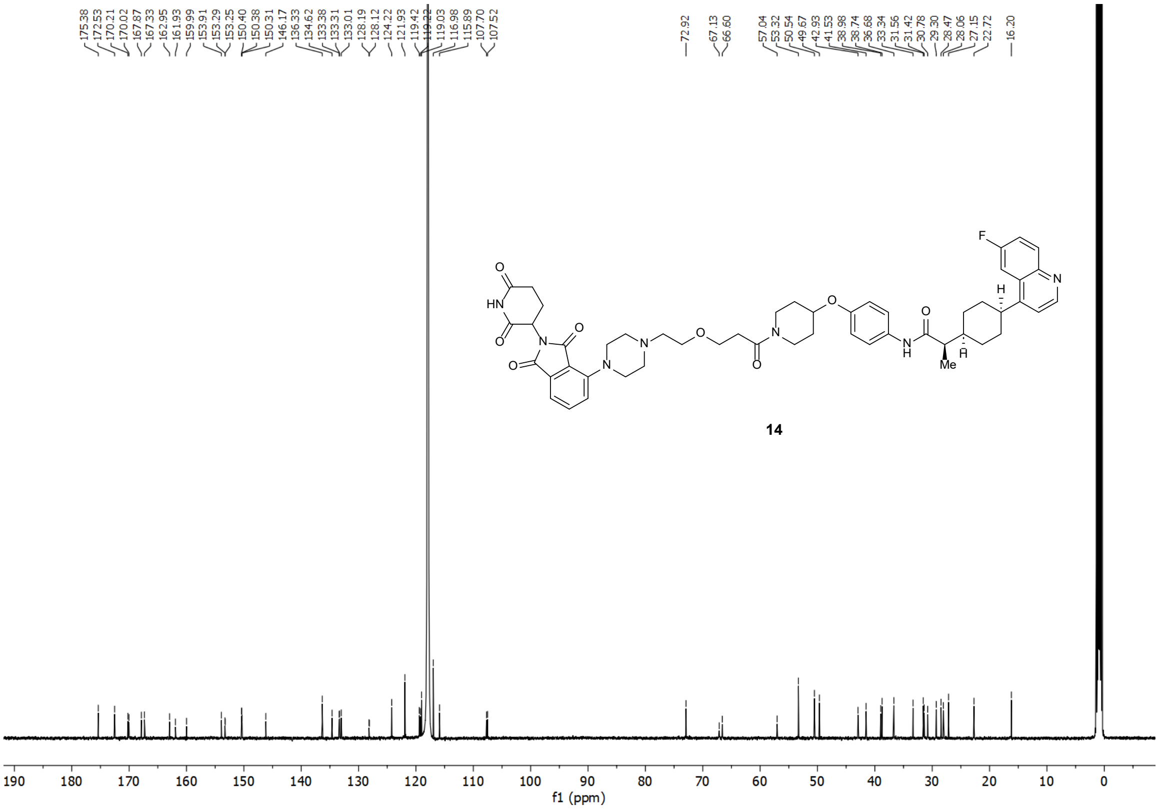

### [1]H-NMR and [13]C-NMR Spectra of PROTAC 15

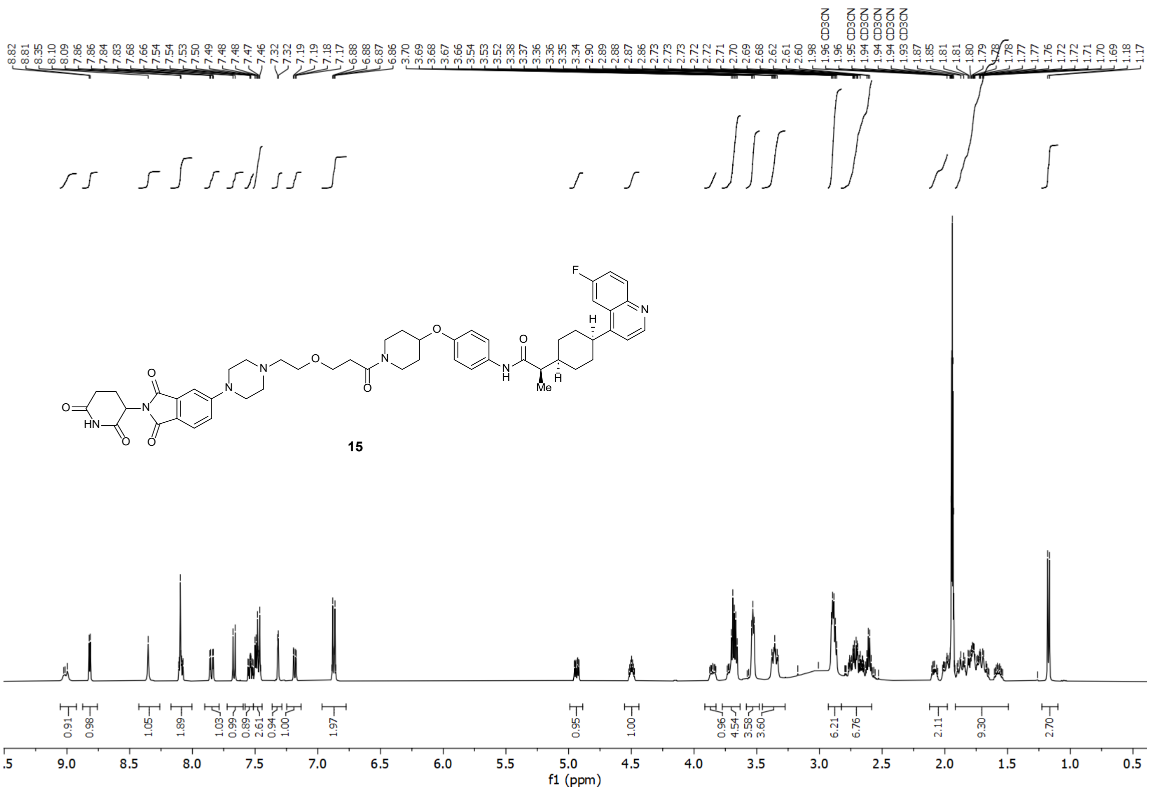


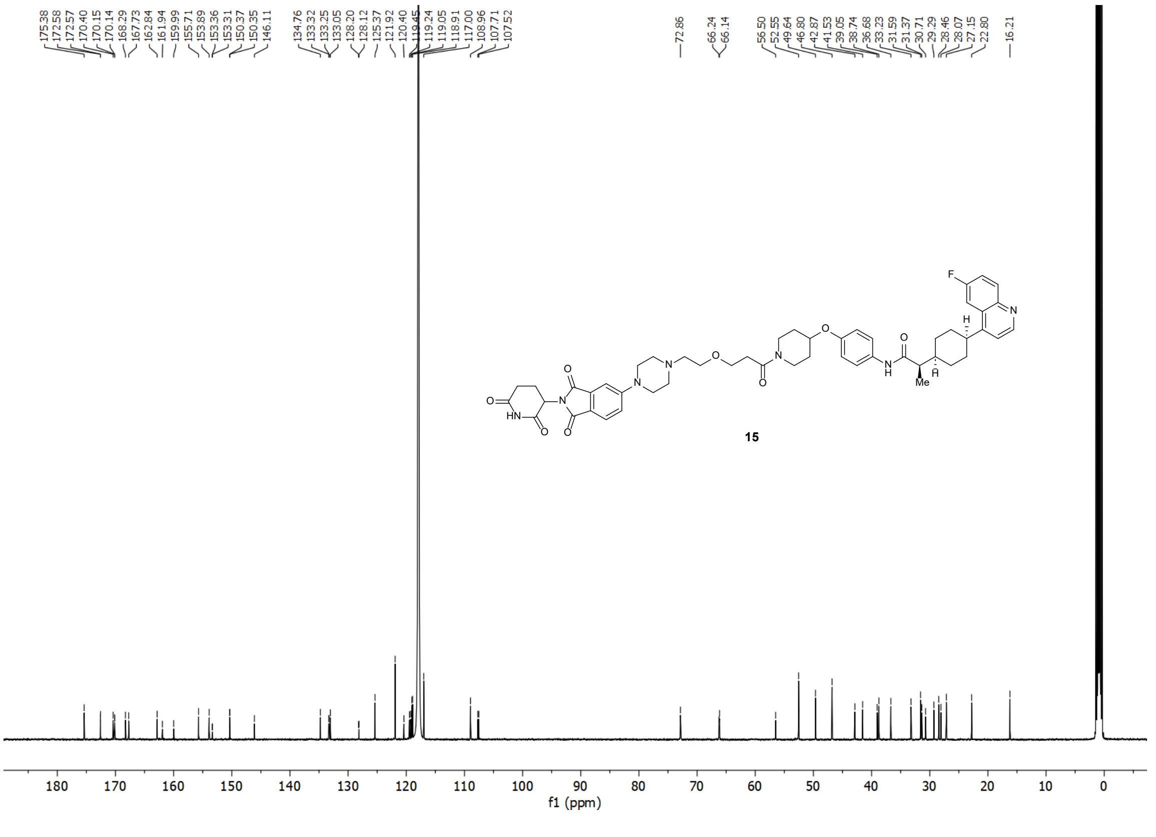

#### $^{1}$H-NMR and $^{13}$C-NMR Spectra of PROTAC 16

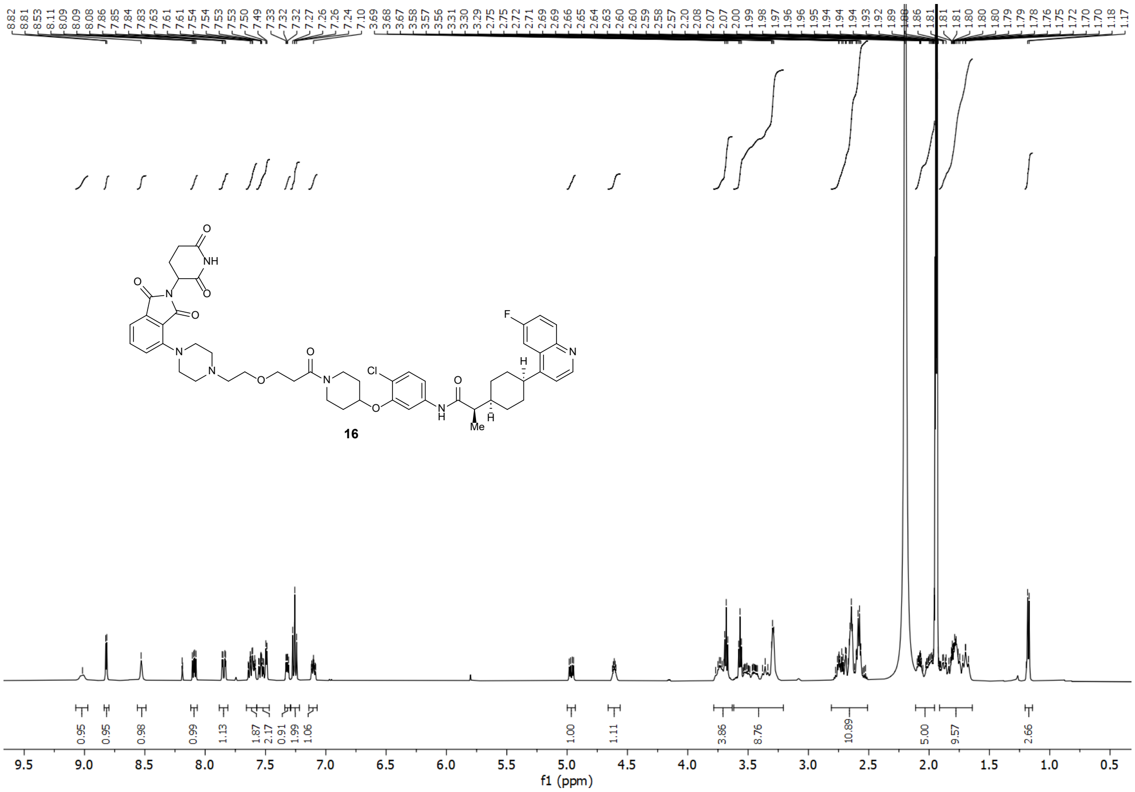


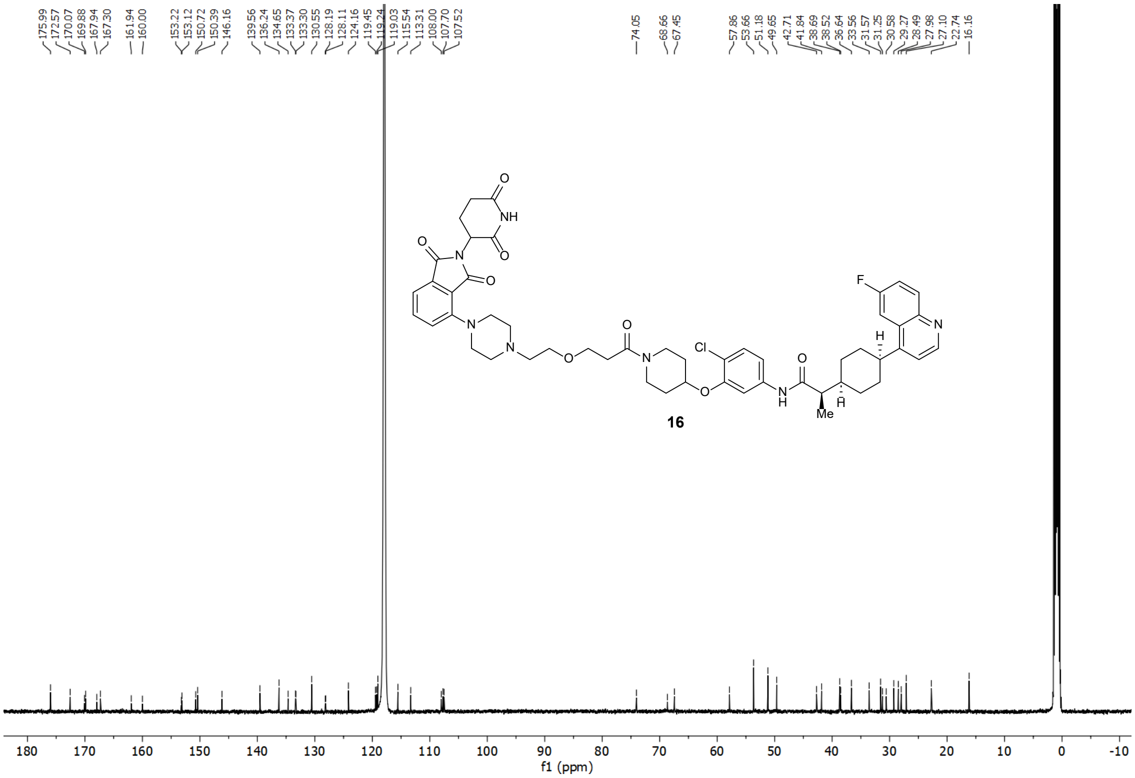

### $^{1}$H-NMR and $^{13}$C-NMR Spectra of PROTAC 17

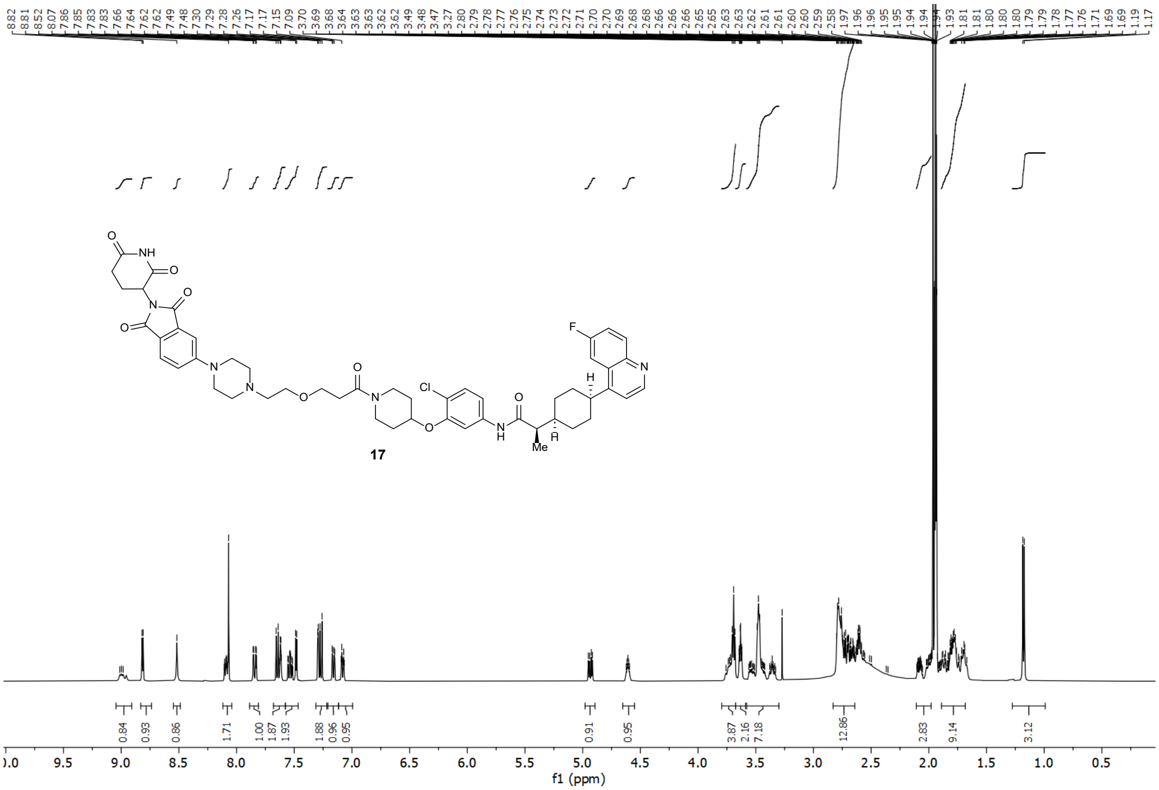


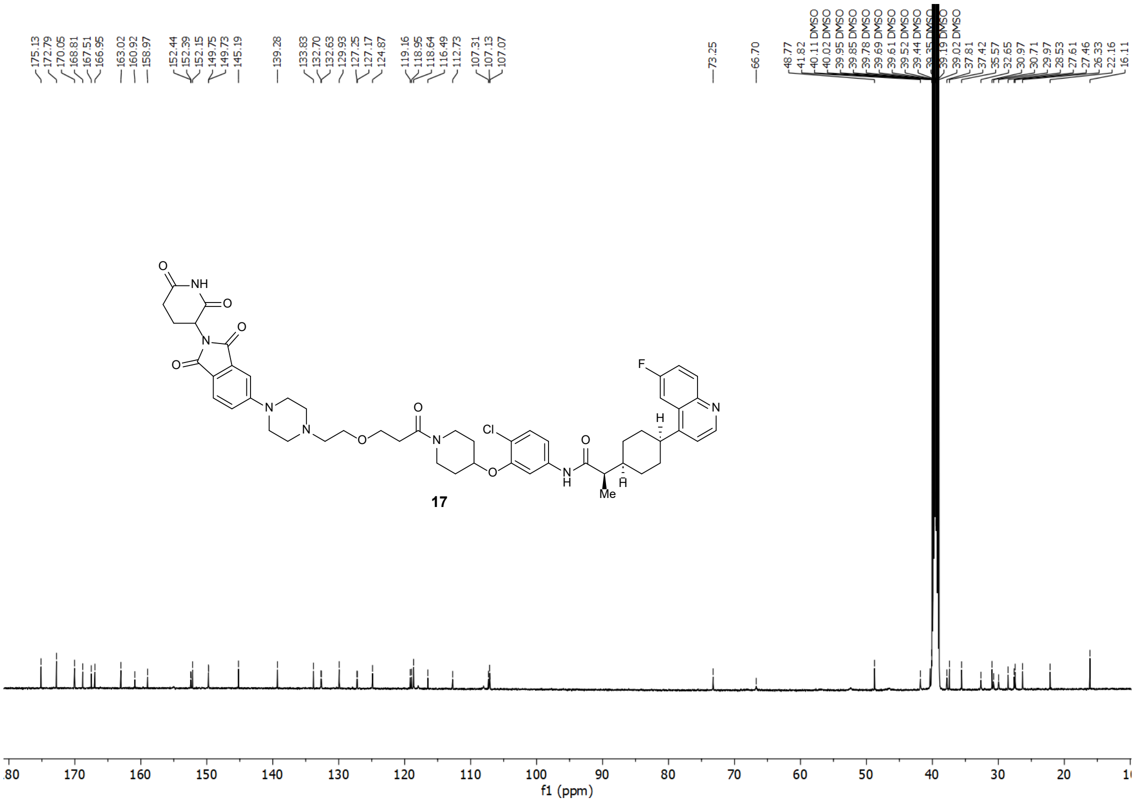

#### $^{1}$H-NMR and $^{13}$C-NMR Spectra of PROTAC 18

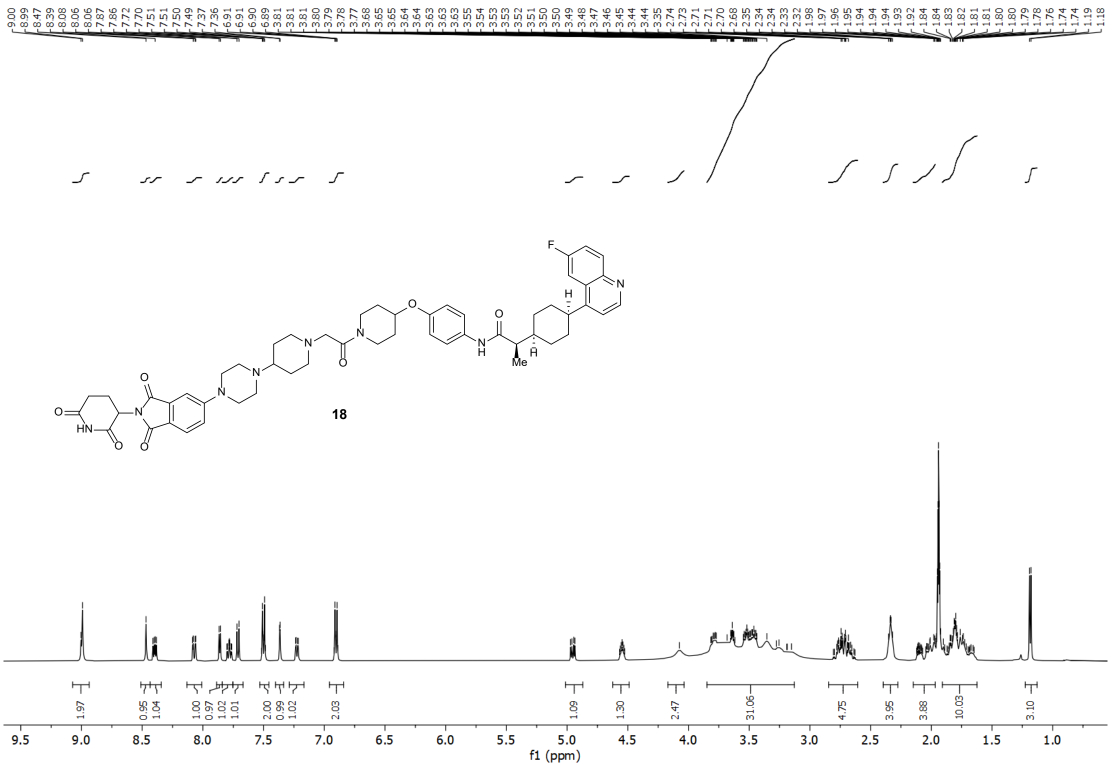


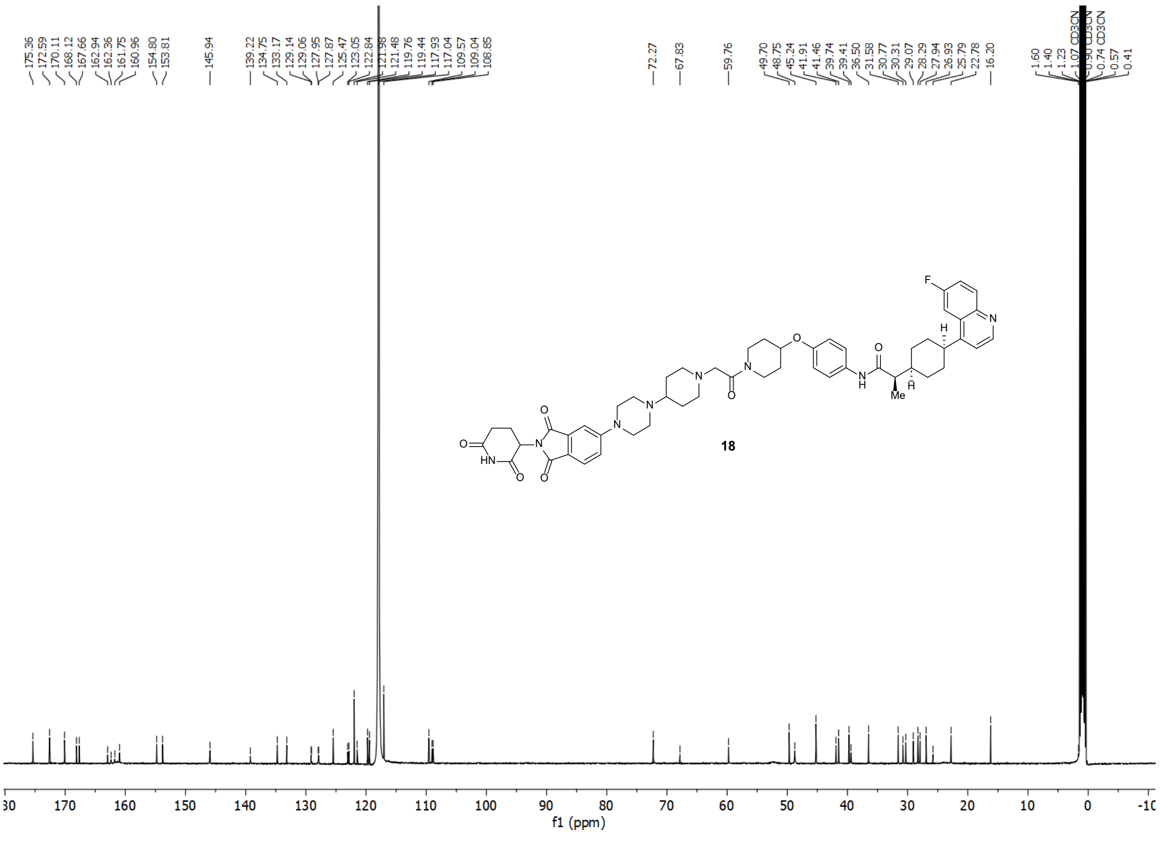

### $^{1}$H-NMR and $^{13}$C-NMR Spectra of PROTAC 19

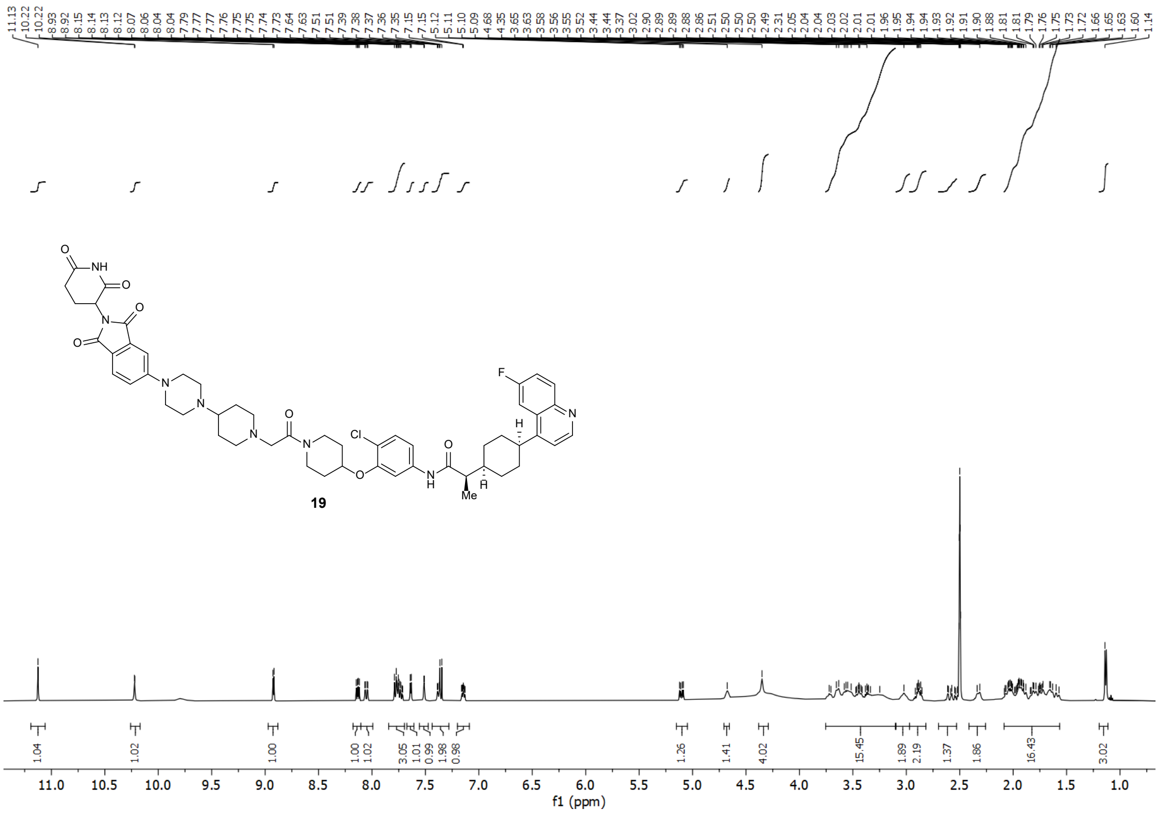

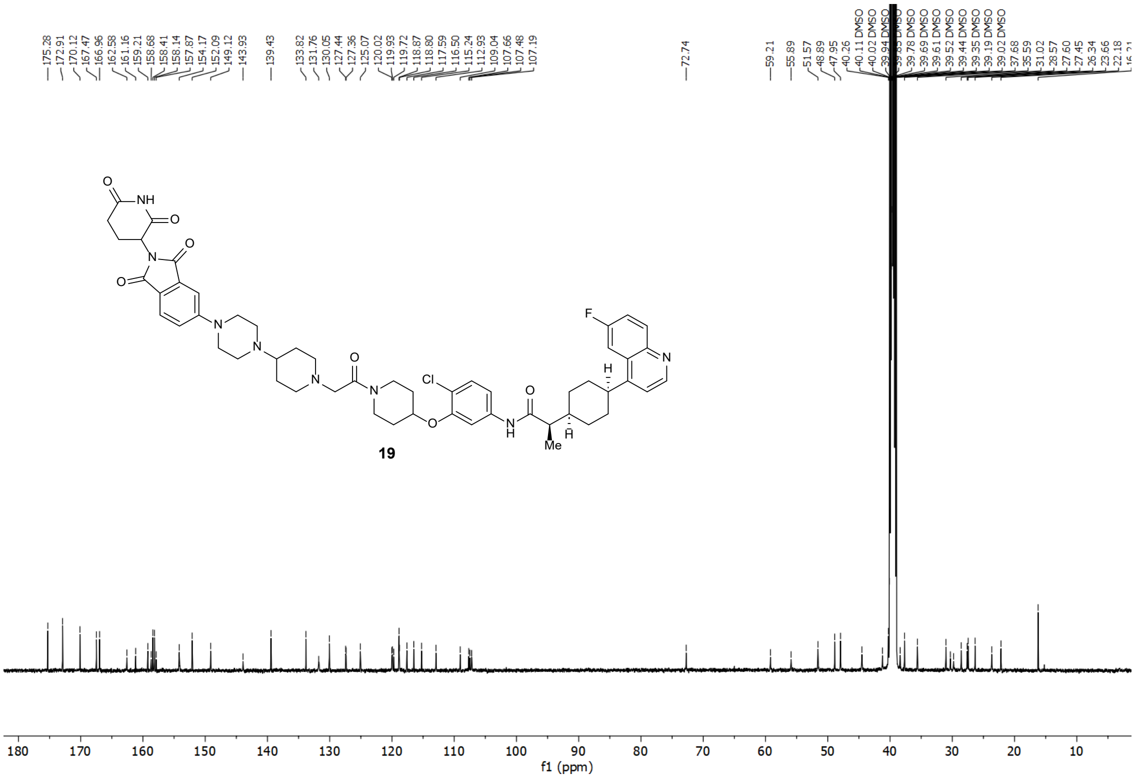

### [1]H-NMR and [13]C-NMR Spectra of PROTAC 20

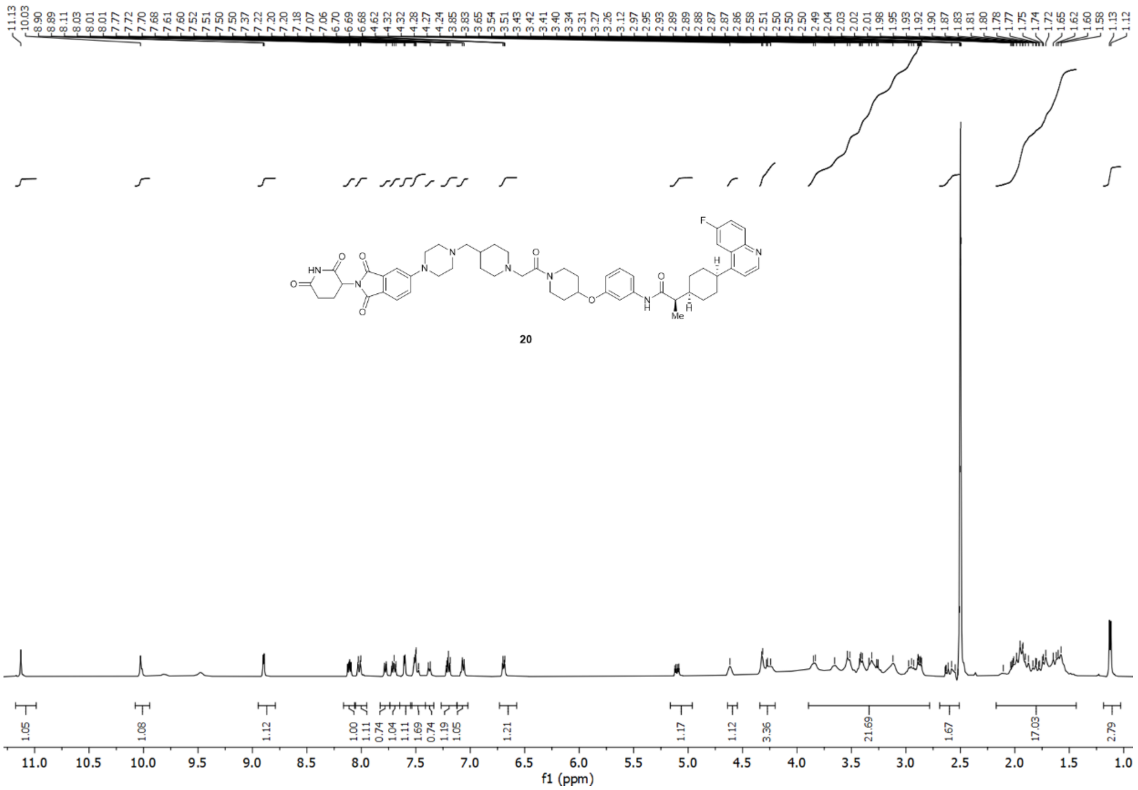


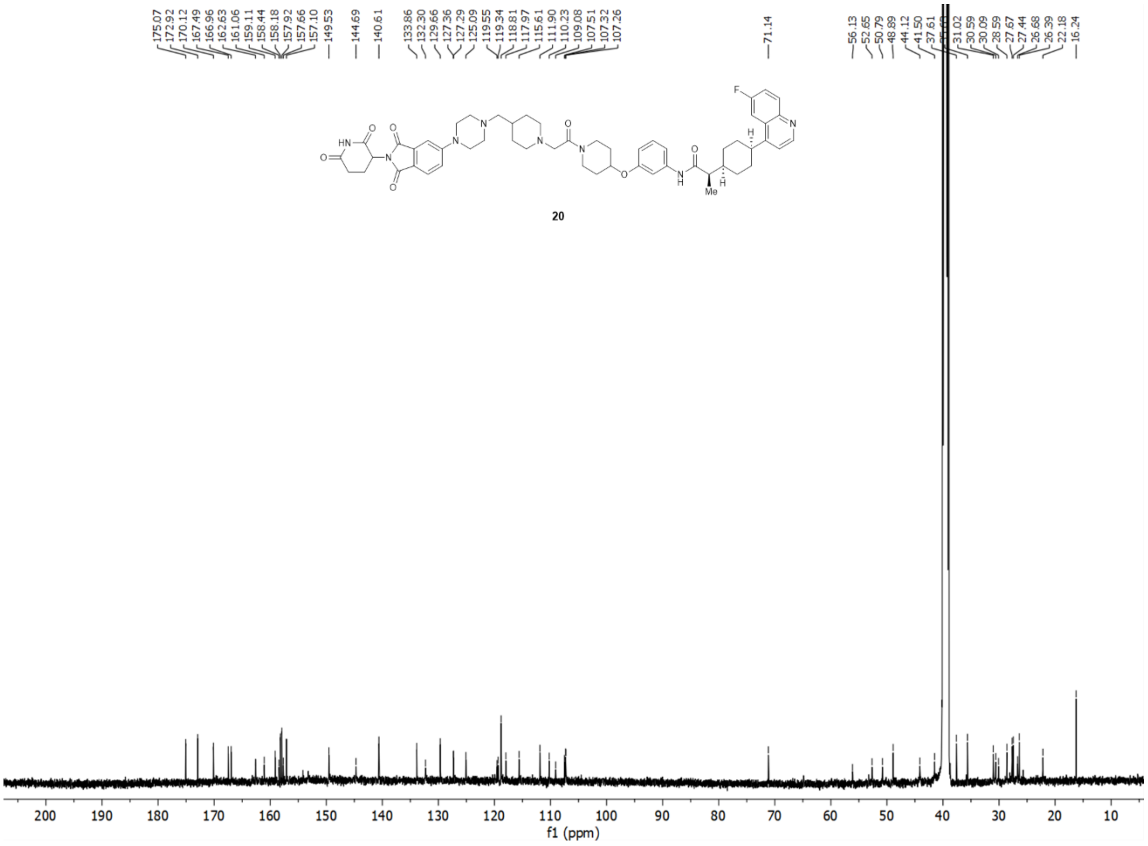

#### HPLC of Compound 20 (NU227326)

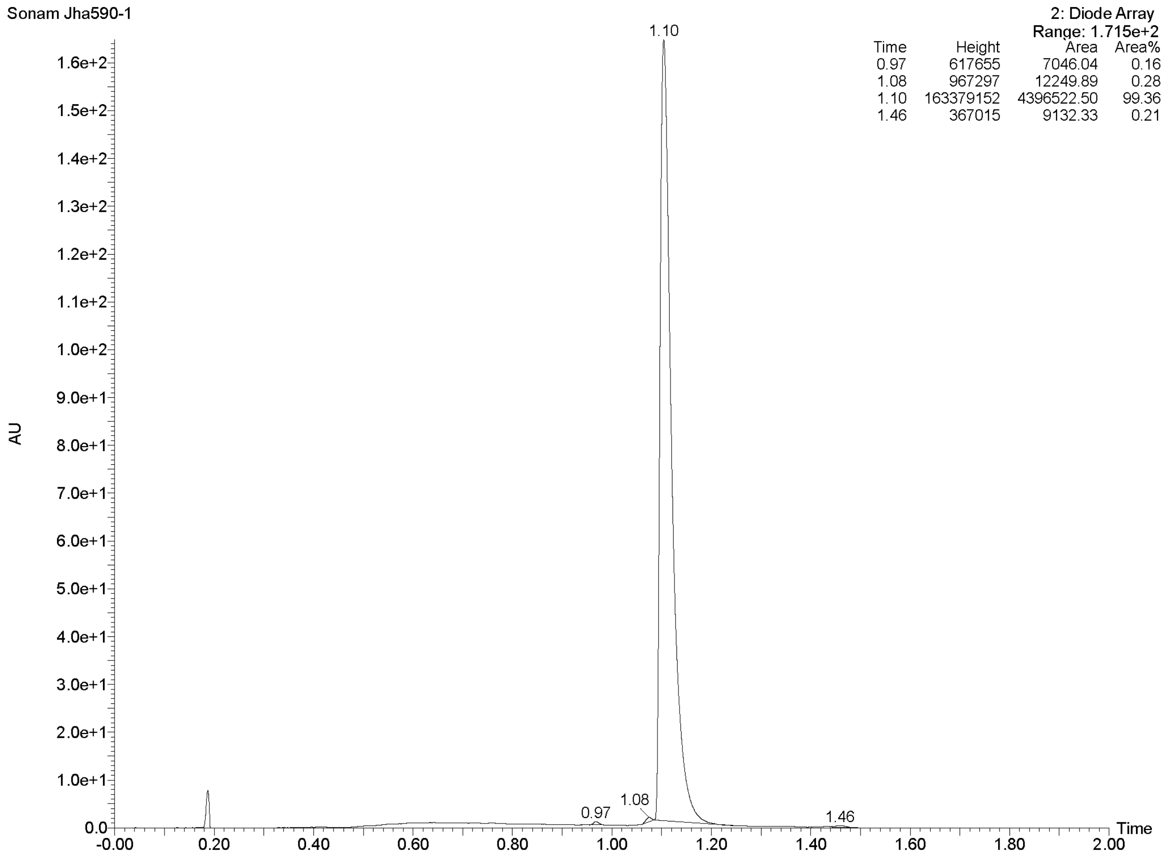

#### $^{1}$H-NMR and $^{13}$C-NMR Spectra of PROTAC 21

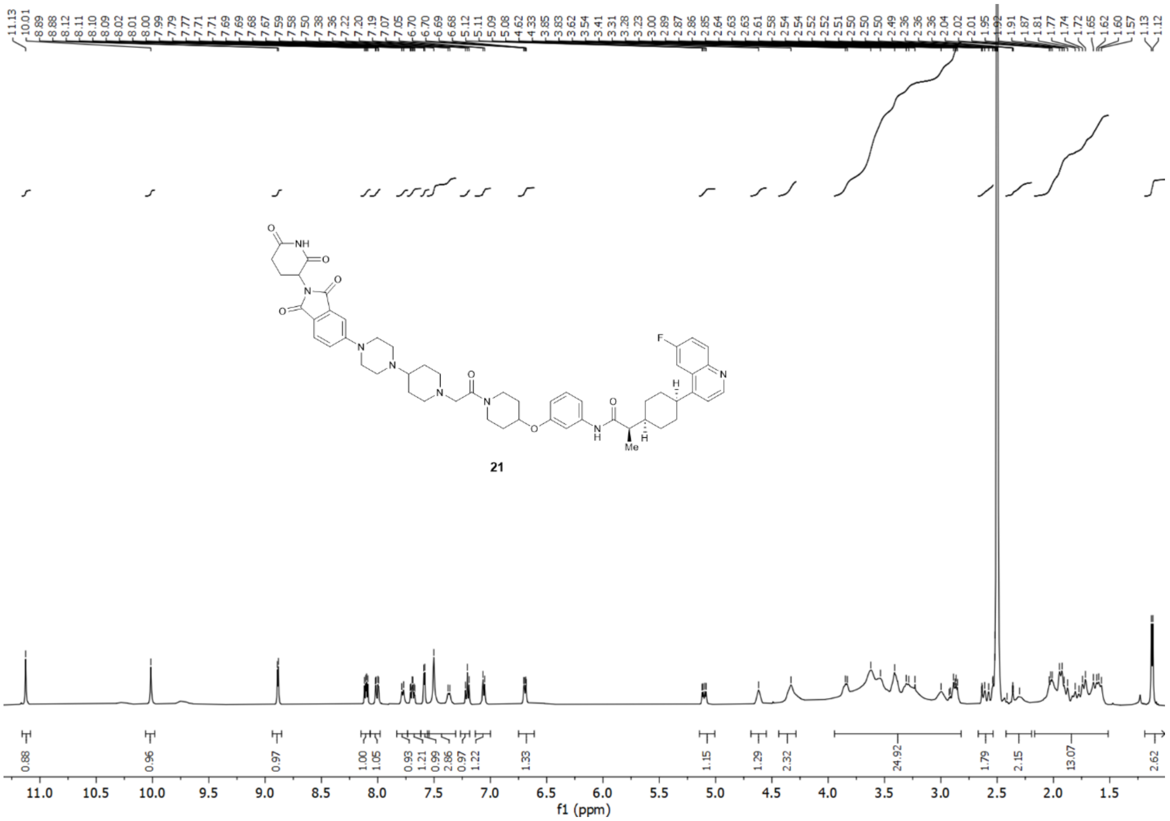


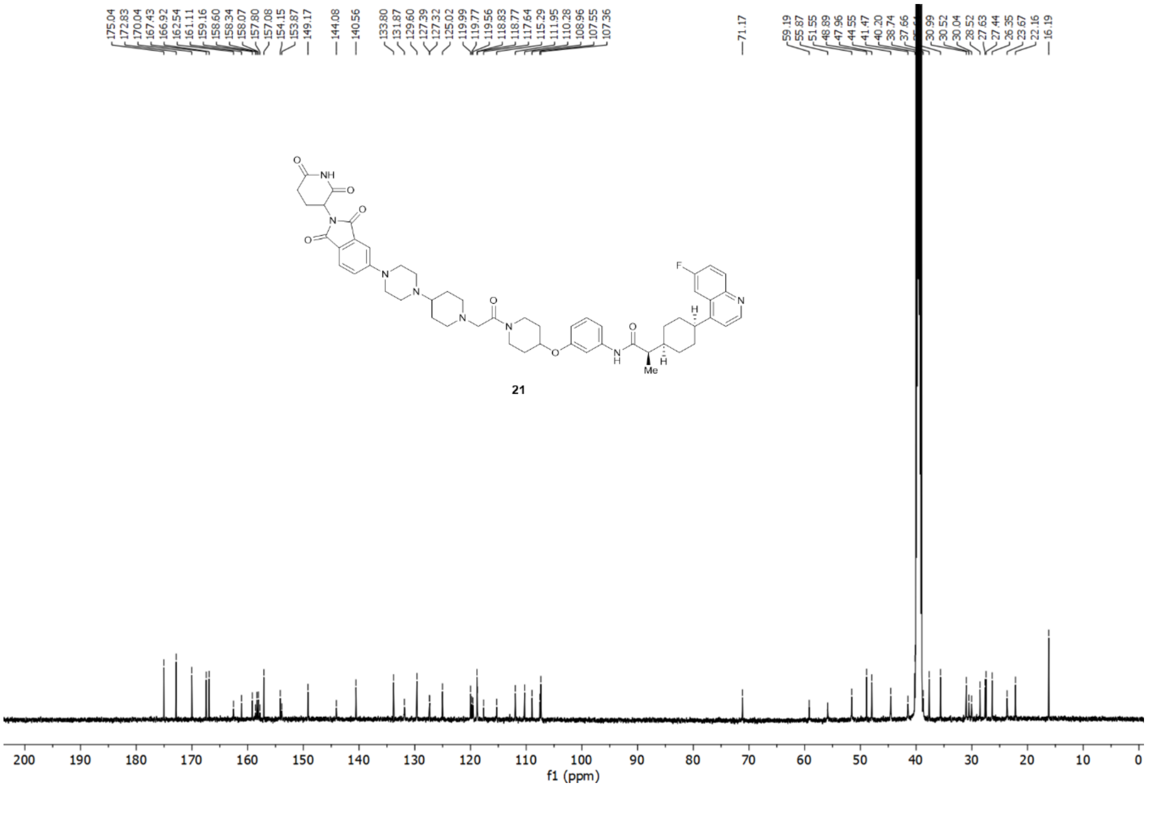

### HPLC of Compound 21 (NU227327)

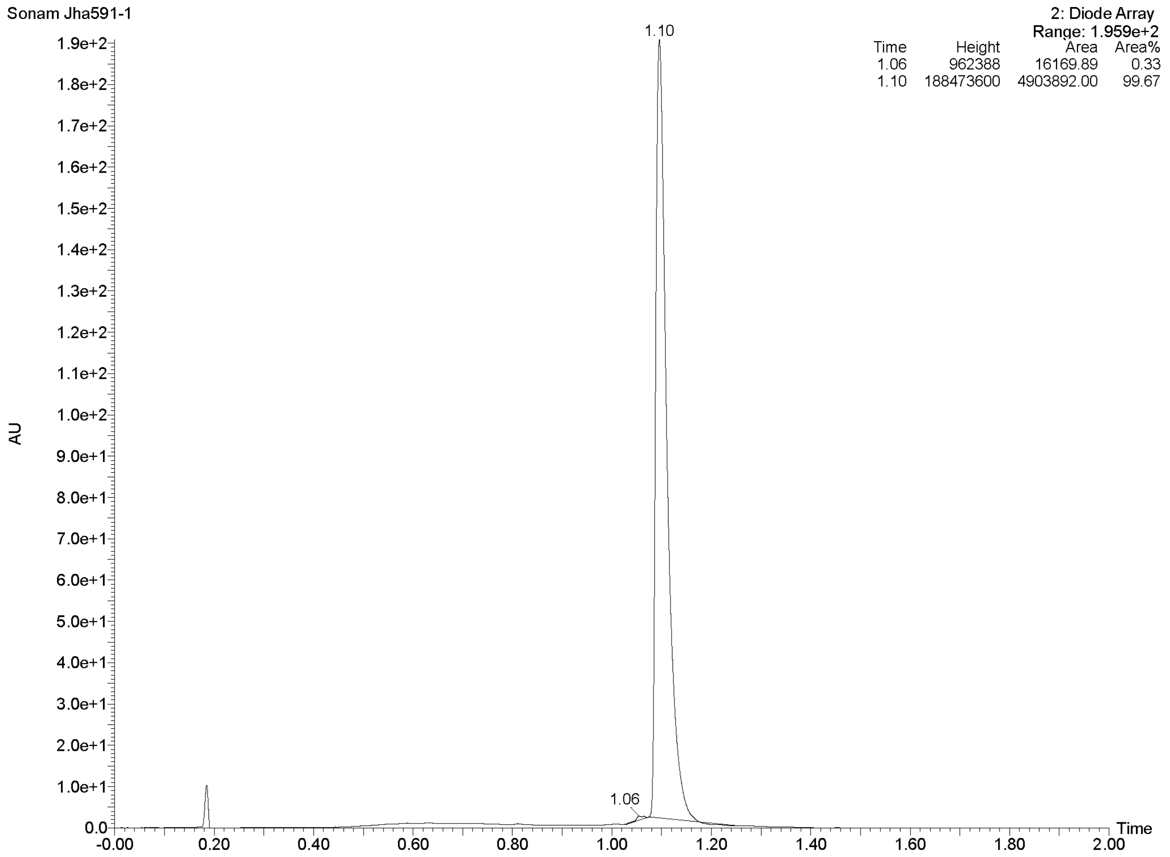

#### $^{1}$H-NMR Spectra of PROTAC 22

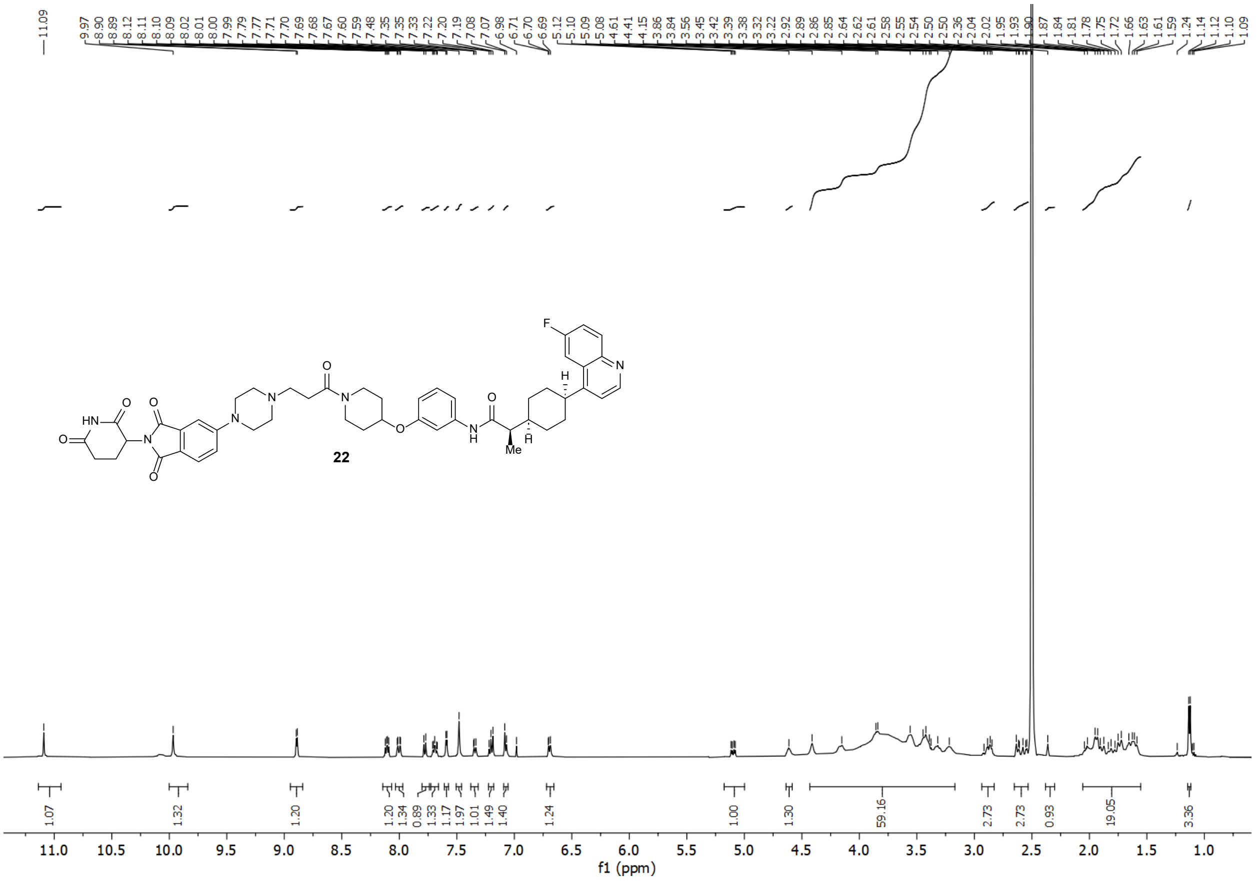

### [1]H-NMR Spectra of PROTAC 23

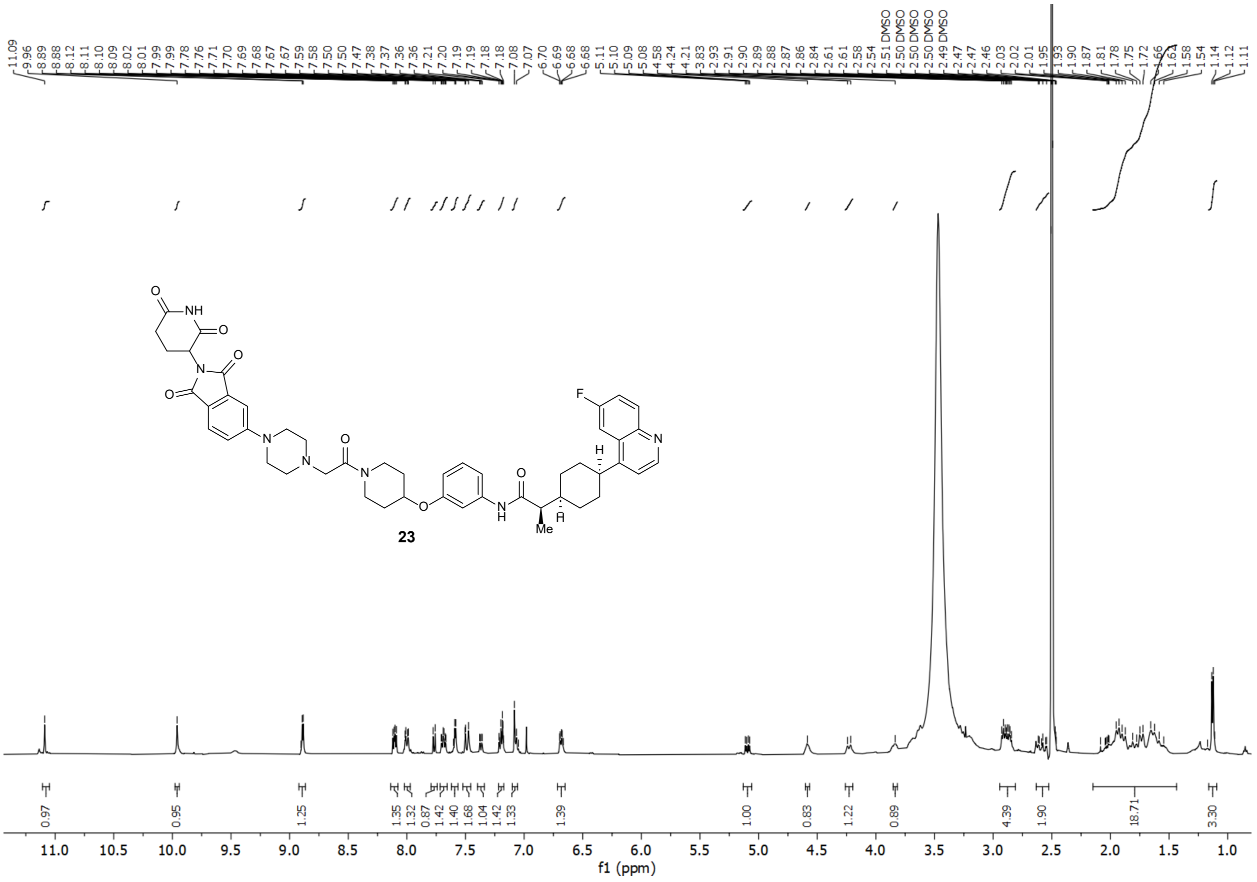

### $^{1}H$-NMR and $^{13}C$-NMR Spectra of PROTAC 24

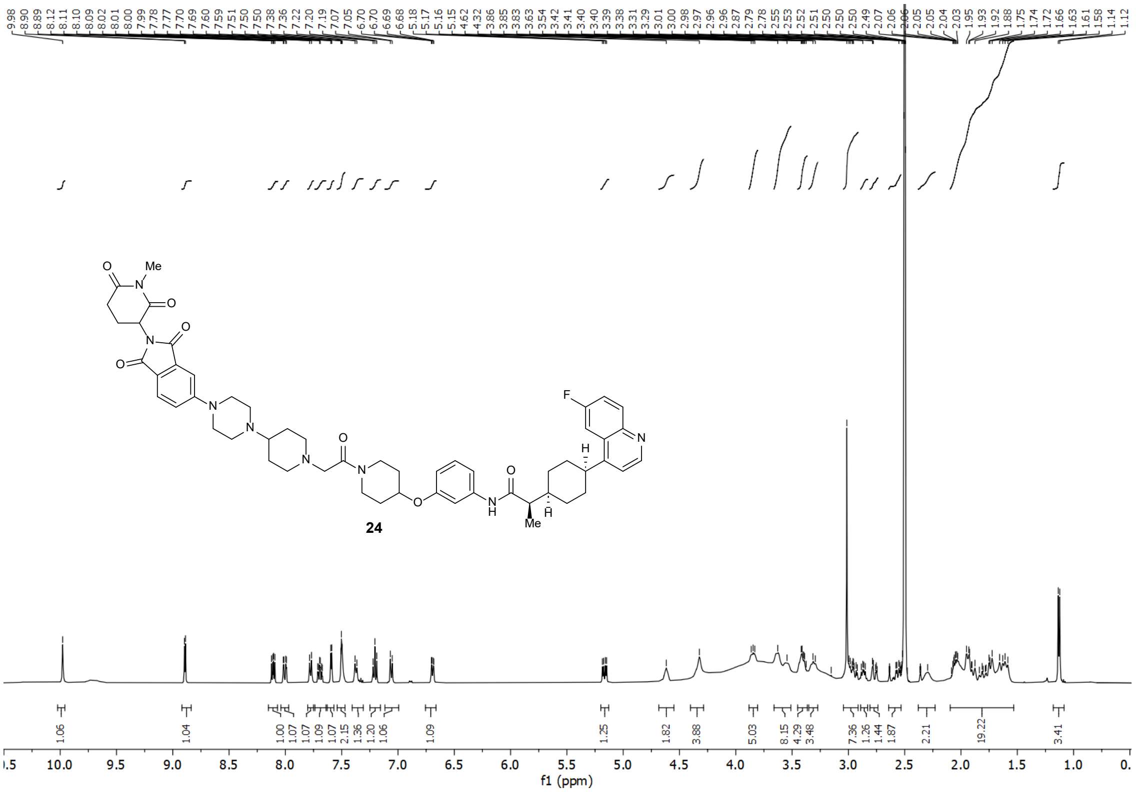


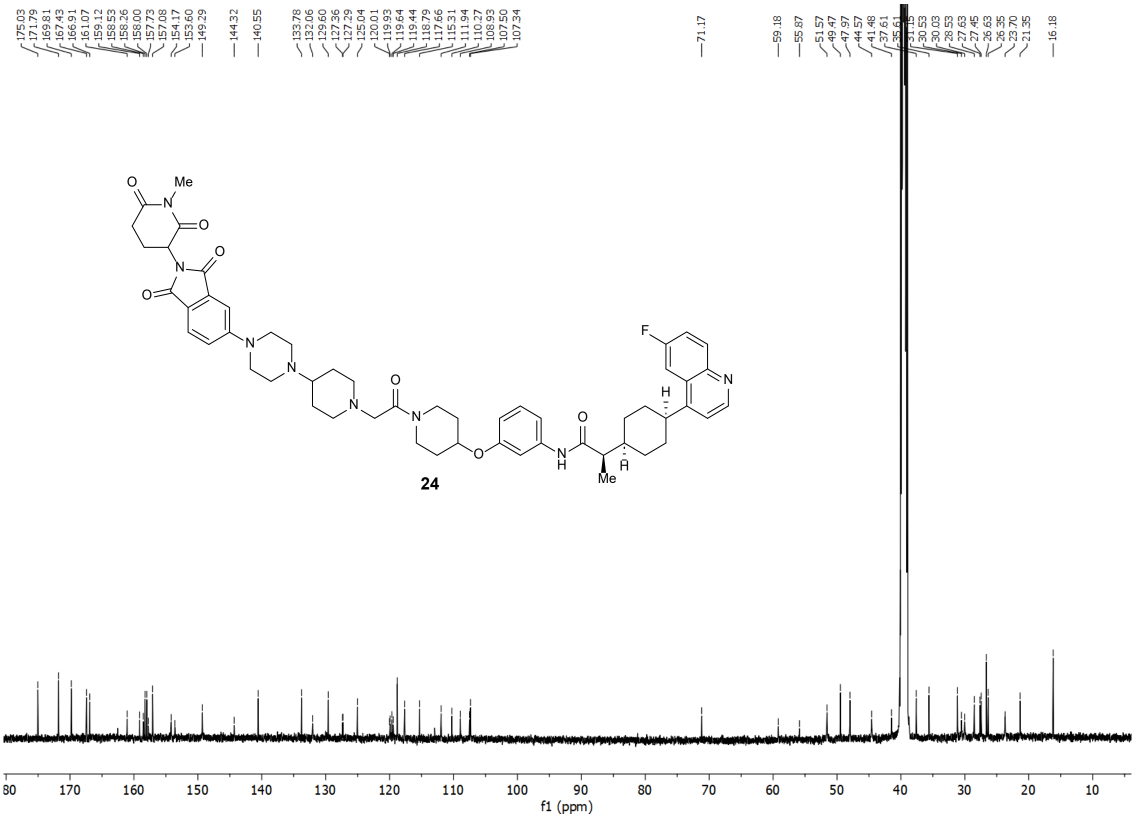

#### Raw western blot images

### Raw data of Western Blot:

Below figures represents the expression of IDO1 and GAPDH obtained by Western Blot in Figure 5.

##### Figure 5: B

Expt: U87 Cells NU227326 at long dose range:
IB – IDO1

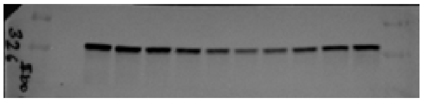

IB – GAPDH

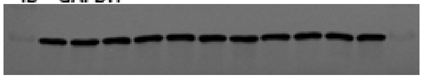

##### Figure 5: D

Expt: GBM43 Cells NU227326 at long dose range:
IB – IDO1

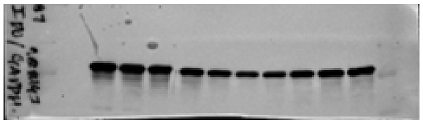

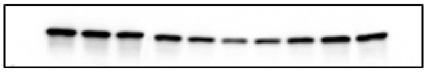

IB – GAPDH

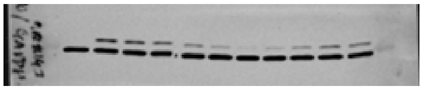

##### Figure 5: B

Expt: U87 Cells NU227327 and NU227326 at long dose range:
IB – IDO1

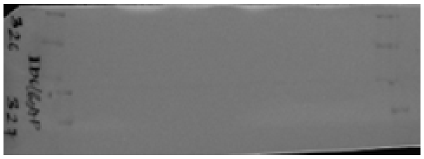

U87 cells NU227326 and NU227327: IB – IDO1

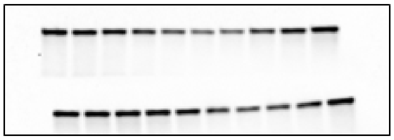

U87 cells NU227326 and NU227327: IB – GAPDH

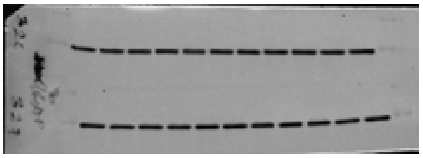

##### Figure 5: G

Expt: U87 Cells treatment with NU227326, NU227428, and NU223618
IB – IDO1

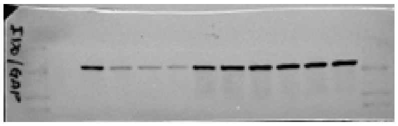

IB – GAPDH

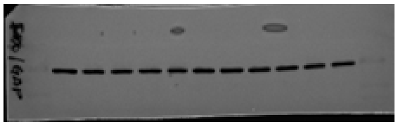

Below figures represents the expression of IDO1 and GAPDH obtained by Western Blot in Figure 6.

#### Figure 6A

Expt: U87 Cells NU227326 degradation kinetics : IB – IDO1

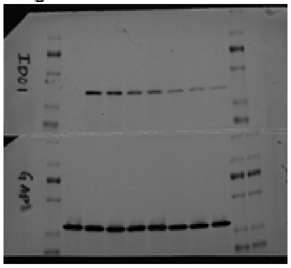

Expt: U87 Cells NU227326 degradation kinetics : IB – IDO1

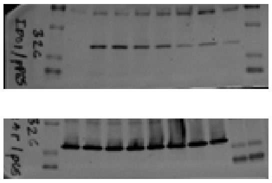

#### Figure 6B and 6C

Expt: U87 Cells NU227326 continuous treatment and post washout : IB – IDO1

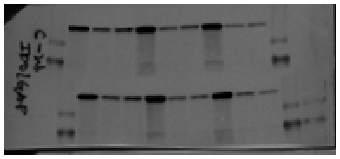

IB – GAPDH

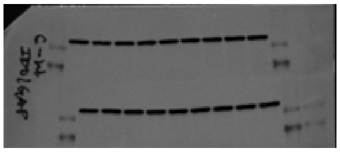

#### Figure 6D

Expt: Competitive inhibition NU223618 and BGB-7204 : IB – IDO1

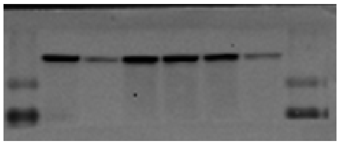

IB – GAPDH

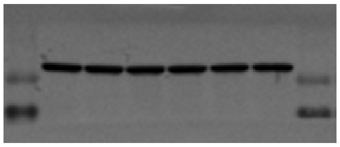

#### Figure 6E

Expt: Competitive inhibition Pomalidomide and MLN4924 : IB – IDO1

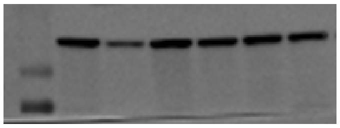

IB – GAPDH

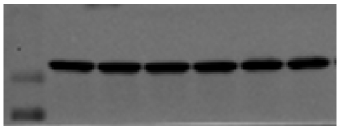

#### Figure 6F

Expt: MG132: IB – IDO1

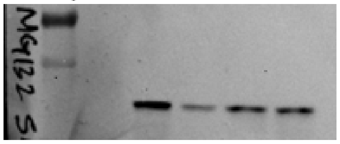

Expt: MG132: IB – GAPDH

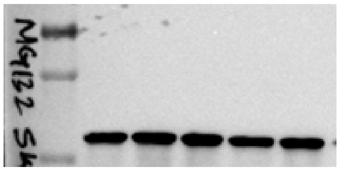

#### Figure 6G and 6H

Expt: PDX-6, PDX-38: IB – IDO1

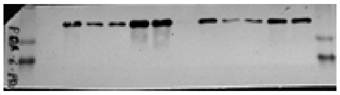

Expt: PDX-6, PDX-38: IB – GAPDH

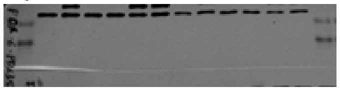

#### Figure 6I

Expt: U87:IDO-GFP: IB – IDO1

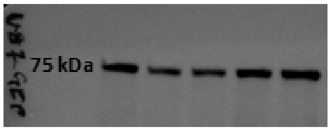


Expt: U87:IDO-GFP: IB – GAPDH

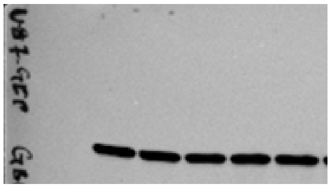

#### Figure 6M

Expt: SW-1990: IB – IDO1

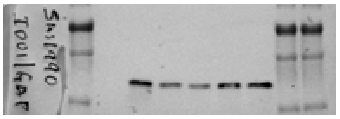

Expt: SW-1990: IB – GAPDH

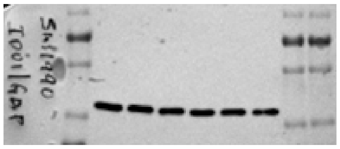

#### Figure 6J, 6K, and 6L

Expt: Tumor cell lines: IB – IDO1

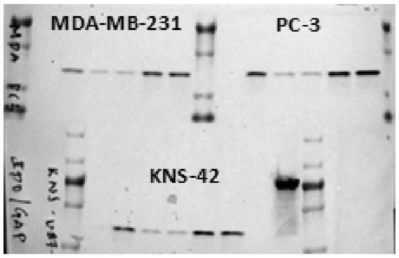


Expt: Tumor cell lines: IB – GAPDH

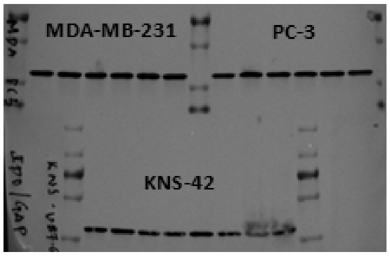


#### Figure 6N

Expt: SKOV-3: IB – IDO1

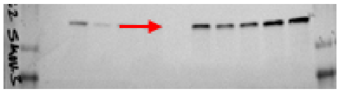

Expt: SKOV-3: IB – GAPDH

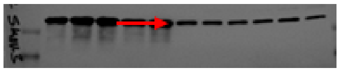
